# Genome-wide data from medieval German Jews show that the Ashkenazi founder event pre-dated the 14^th^ century

**DOI:** 10.1101/2022.05.13.491805

**Authors:** Shamam Waldman, Daniel Backenroth, Éadaoin Harney, Stefan Flohr, Nadia C. Neff, Gina M. Buckley, Hila Fridman, Ali Akbari, Nadin Rohland, Swapan Mallick, Jorge Cano Nistal, Jin Yu, Nir Barzilai, Inga Peter, Gil Atzmon, Harry Ostrer, Todd Lencz, Yosef E. Maruvka, Maike Lämmerhirt, Leonard V. Rutgers, Virginie Renson, Keith M. Prufer, Stephan Schiffels, Harald Ringbauer, Karin Sczech, Shai Carmi, David Reich

## Abstract

We report genome-wide data for 33 Ashkenazi Jews (AJ), dated to the 14^th^ century, following a salvage excavation at the medieval Jewish cemetery of Erfurt, Germany. The Erfurt individuals are genetically similar to modern AJ and have substantial Southern European ancestry, but they show more variability in Eastern European-related ancestry than modern AJ. A third of the Erfurt individuals carried the same nearly-AJ-specific mitochondrial haplogroup and eight carried pathogenic variants known to affect AJ today. These observations, together with high levels of runs of homozygosity, suggest that the Erfurt community had already experienced the major reduction in size that affected modern AJ. However, the Erfurt bottleneck was more severe, implying substructure in medieval AJ. Together, our results suggest that the AJ founder event and the acquisition of the main sources of ancestry pre-dated the 14^th^ century and highlight late medieval genetic heterogeneity no longer present in modern AJ.

## Introduction

Ashkenazi Jews (AJ) emerged as a distinctive ethno-religious cultural group in the Rhineland and Northern France in the 10^th^ century. The AJ population since expanded substantially, both geographically, first to Eastern Europe and recently beyond Europe, and in number, reaching about 10 million today [1–4]. The AJ population today harbors dozens of recessive pathogenic variants that occur at higher frequency than in any other population [5–9], implying that AJ descend from a small set of ancestral founders [10–14]. This Ashkenazi “founder event” is also manifested by four mitochondrial haplogroups carried by as many as 40% of AJ [15, 16]. More recently, studies found high rates of identical-by-descent (IBD) sharing in AJ, that is, near-identical long haplotypes present in unrelated individuals, a hallmark of founder populations [17–21]. Quantitative modeling suggested that AJ experienced a sharp reduction in size (a “bottleneck)” in the late Middle Ages and that the (effective) number of founders was in the hundreds [19, 22–25].

The origins of early Ashkenazi Jews, as well as the history of admixture events that have shaped their gene pool, are subject to debate. In historical research, there are two main hypotheses regarding the identity of the early AJ: either Jews who lived at the Germanic frontiers since late Roman times, or medieval migrants from the established Jewish communities of the Italian peninsula (SI 1). Genetic evidence supports a mixed Middle Eastern (ME) and European (EU) ancestry in AJ. This is based on uniparental markers with origins in either region [15, 16, 26–29], as well as autosomal studies showing that AJ have ancestry intermediate between ME and EU populations [18, 19, 23, 30–34]. Recent modeling suggested that most of the European ancestry in AJ is consistent with Southern European-related sources, and estimated the total proportion of European ancestry in AJ as 50-70% [19, 35]. While the Ashkenazi population is overall highly genetically homogeneous [17, 33, 34], there are subtle average differences in ancestry between AJ with origins in Eastern vs Western Europe [23, 30, 36].

While recent work has advanced our understanding AJ population genetics, open questions remain. For example, can we localize the founder event, or events, in time and space? When and where did admixture events occur in AJ history? Studying the genomes of individuals who lived closer to the time of AJ formation may shed light on these questions.

We present the first DNA study of historical Jews, focusing on AJ from 14^th^-century Erfurt, Germany. The Erfurt Jewish community existed between the late 11^th^ century to 1454, with a short gap following a 1349 massacre [37, 38]. We report 33 genomes from individuals whose skeletons were extracted in a salvage excavation. Our results demonstrate that Erfurt Ashkenazi Jews (EAJ) are genetically highly similar to modern Ashkenazi Jews (MAJ), implying little gene flow into AJ gene since the 14^th^ century. Further analysis demonstrated that EAJ were more genetically heterogeneous than MAJ, with multidisciplinary evidence supporting the presence of two sub-groups, one of which had higher Eastern European affinity compared to MAJ. The EAJ population shows strong evidence of a recent sharp bottleneck, based on the distribution of mitochondrial haplogroups, high levels of runs of homozygosity, and the presence of AJ-enriched alleles, including pathogenic variants.

## Results

### Historical and archaeological context, sample collection, and ethics

The first Jewish community of Erfurt (pre-1349) was the oldest in Thuringia, and its cemetery also served nearby towns [39, 40]. During the 1349 pogrom, most Jews of Erfurt and nearby communities were murdered or expelled [37, 41, 42]. Jews returned to Erfurt around 1354 to form the second community [43], which was one of the largest in Germany [40] (SI 1). The individuals we studied were buried in the south-western part of the medieval Jewish cemetery of Erfurt, which underwent salvage excavations in 2013 (Methods Section 1). The site is shown in Figure S1 and the layout of the cemetery is shown in Figure S2. The excavation site was likely used by the second community, although archaeological evidence is uncertain (Methods Section 1). The cause of death could be determined only for I14904, who was killed by several blows to the head by a sharp object.

Jewish rabbinical law, which was followed by EAJ (SI 1) prohibits exhumation of Jews for most purposes, and also proscribes disturbing the dead. Established rabbinical ruling on ancient DNA studies of Jewish individuals did not exist before the advent of the technology [44], and there is no centralized authority for establishing Jewish rabbinical guidance. As part of this study, we engaged with rabbinical authorities who reviewed our proposed research plan and approved the project under the conditions that only detached teeth are collected and that the analysis is performed only on already-excavated individuals and does not involve excavation specifically for the purpose of ancient DNA research. The study was also approved by the Jewish community of Thuringia, Germany. Following these guidelines, we sampled teeth from 38 skeletal remains.

### Dataset

We followed existing protocols for DNA extraction, library preparation, and enrichment for about 1.24 million single nucleotide polymorphisms (SNPs). We then sequenced the enriched and non-enriched libraries and used multiple metrics to demonstrate very low levels of contamination (Methods Section 2; Table S1). We obtained genome-wide data passing quality control for 33 individuals: 19 females and 14 males. The median proportion of endogenous DNA was 0.03 (range 0.0003-0.67; Table S1), and the median coverage on autosomal targets was 0.45x (range 0.01-2.48x; Table S1). The median number of SNPs covered by at least one sequence was 383k (range 11-842k; Table S1; Figure S3). The estimated ages at death (Methods Section 1) ranged between 5 to at least 60 years old, with 14/33 (42%) estimated to be younger than 20 (Table S2). Children had significantly lower coverage than adults (P=6.7·10^−7^; Methods Section 2).

**Table 1.** Ashkenazi pathogenic variants detected in Erfurt. For each of the 11 high-confidence Ashkenazi pathogenic variants carried by EAJ, we indicate the disease name, the gene, and the variant in HGVS (Human Genome Variation Society) nomenclature. The c.68_69delAG BRCA1 variant is also known as 185delAG. For imputed variants, we provide the marginal posterior probability in *PHCP* for having at least one alternate allele. We further provide the carrier IDs, the allele frequency in MAJ and non-Finnish Europeans (*gnomAD*), and the number (out of four) of Ashkenazi-specific pre-conception screening (PCS) panels where the gene is included (Methods Section 9).

| Disease | Gene | Variant | Carrier ID and *PHCP* probability of carrying at least one alternate allele | Frequency in modern AJ (*gnomAD*) | Frequency in non-Finnish Europeans (*gnomAD*) | Gene in PCS panels |
| --- | --- | --- | --- | --- | --- | --- |
| Retinitis pigmentosa 59 | *DHDDS* | c.124A>G | I14903, 0.985 | 0.52% | 0.004% | 4/4 |
| Gaucher disease, type 1 | *GBA* | c.84dupG | I13861, 1.000 | 0.08% | 0.002% | 4/4 |
| Usher syndrome, type 3 | *CLRN1* | p.N48K | I14897, 0.980 | 0.58% | 0.008% | 4/4 |
| Factor XI deficiency | *F11* | p.E135X | I13870, genotyped | 1.71% | 0.03% | 3/4 |
| Factor XI deficiency | *F11* | p.F301L | I13865, genotyped | 2.38% | 0.03% | 3/4 |
| Cystic fibrosis | *CFTR* | p.G542X | I14736, 1.000 | 0.15% | 0.04% | 4/4 |
| Parkinson's disease | *LRRK2* | p.G2019S | I14739, genotyped | 0.84% | 0.03% | 0/4 |
| Acyl CoA dehydrogenase deficiency | *ACADS* | c.319C>T | I14737, 0.999 | 1.88% | 0.02% | 0/4 |
| Familial Mediterranean fever | *MEFV* | p.V726A | I14739, 0.993 | 3.93% | 0.09% | 2/4 |
| Glycogen storage disease, type 1A | *G6PC* | p.R83C | I13870, 0.970 | 0.66% | 0.05% | 4/4 |
| Breast/ovarian cancer predisposition | *BRCA1* | c.68_69delAG | I13861, 1.000 | 0.41% | 0.009% | 0/4 |

We identified two high confidence families (Methods Section 2): family A, with a mother, a son, and a daughter; and family B, with a father (the one killed by strokes to the head) and a daughter (Figure S4). The two children of family A were buried next to each other, as were the two members of family B (Figure S2). The mother of family A was buried three rows away from her children, in an orientation opposite to all other burials (Figure S2). Further inspection of the DNA data suggested that three more individuals, who were all buried next to each other (Figure S2), might have been second-degree relatives (Figure S5). Two of them were the only ones in our sample to carry the U5a1a2a mtDNA haplogroup (Table S2). However, the data for this trio is also consistent with a first-degree relationship or no relationship, likely due to their low coverage (13k, 15k, and 38k SNPs).

Radiocarbon dating of ten individuals demonstrated that all lived between about 1270-1400 CE (Table S2; Table S3), as expected if these individuals had belonged to the medieval Erfurt Jewish community. The reconstructed date was nearly equally likely to be pre-1349 or post-1349, the year of the pogrom (Figure S6). This occurred due to a wiggle in the ^14^C calibration curve just around 1349 (Figure S7). Therefore, ^14^C dating is not informative on whether the site was used by the first or second community. In accordance with the 14^th^-century Black Death affecting Erfurt at a time when Jews no longer lived in the city (SI 1), we found no traces of *Yersinia pestis* bacteria in the DNA sequences (Methods Section 2, Table S4).

### Ancestry estimation

To analyze the ancestry of the EAJ individuals, we represented their genomes as pseudo-haploids using a single random read for each covered SNP. We merged EAJ with the Human Origins (HO) dataset of modern genomes (about 593k autosomal SNPs, all also enriched in EAJ), which included seven Ashkenazi Jews and 86 other Jews. We projected the EAJ individuals on principal components (PCs) learned from the West-Eurasian individuals of the HO dataset (n=994; Methods Section 3). Eight EAJ had (post-merging) coverage of fewer than 50k SNPs, which did not allow reliable projection (Figure S8). We designated these individuals as *low-coverage*, and excluded them from the principal components analysis (PCA).

In the PCA plot, EAJ individuals clustered with those of MAJ, but had more variability along the European–Middle Eastern cline (Figure 1). Higher variability in EAJ relative to MAJ was also observed when projecting a much larger MAJ sample (Figure S9). Inspection of the PCA plot suggested the Erfurt individuals may be divided into two sub-groups. We used *K*-means to cluster the individuals (with *K*=2) based on their PC1 and PC2 coordinates (see below for statistical tests for the presence of two distinct groups). One group falls closer to individuals from European (EU) populations (“Erfurt-EU”), while the other is closer to Middle Eastern (ME) populations (“Erfurt-ME”).

**Figure 1.**
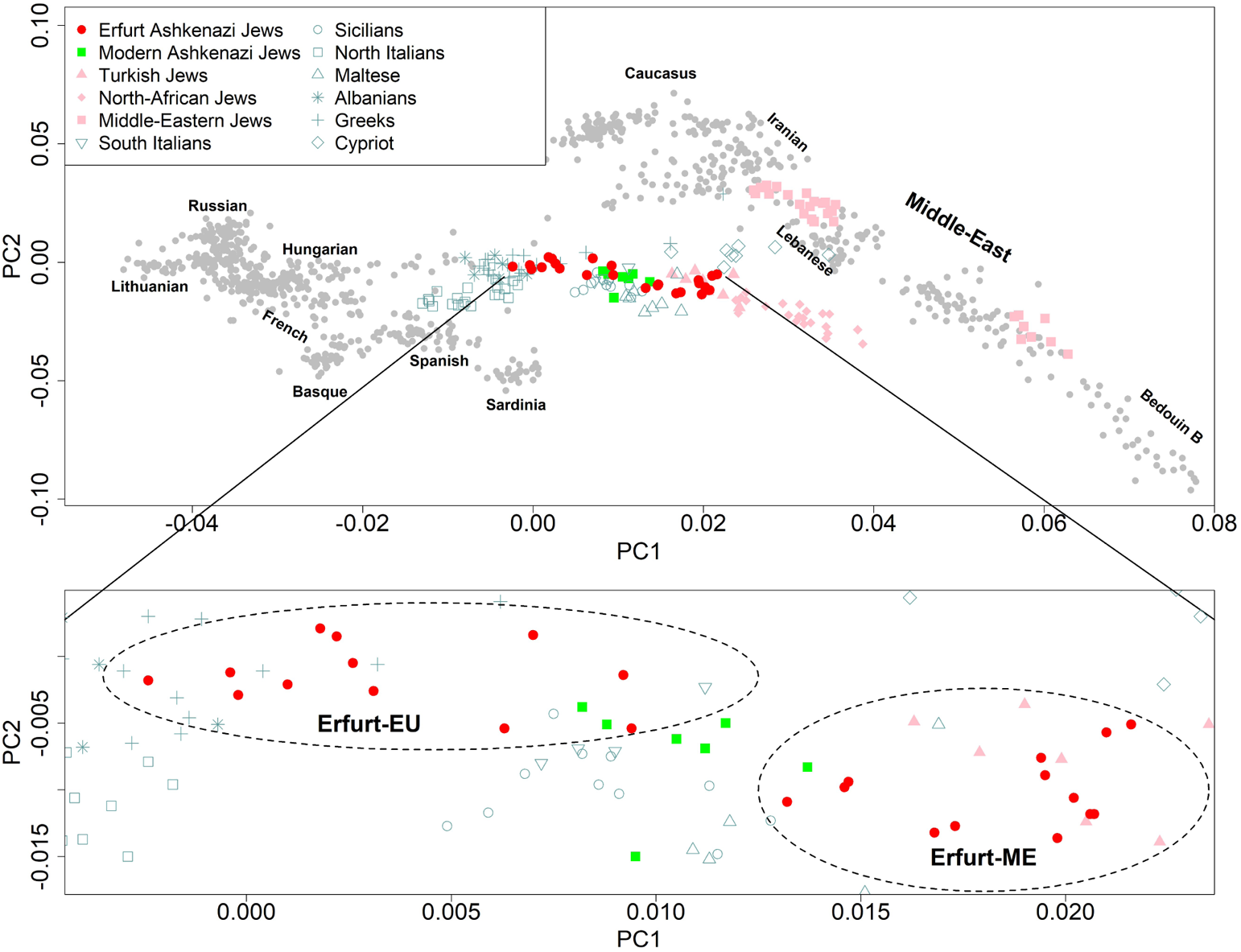
Principal components analysis (PCA). We learned the principal components (PCs) using West Eurasian populations [45] and projected the Erfurt individuals (filled red circles) onto the inferred axes. Modern Ashkenazi Jews (green squares), Jews of non-Ashkenazi origin (pink shapes), and Mediterranean populations (teal shapes) are highlighted. The inset zooms in on the region that contains AJ individuals.

To further characterize the two Erfurt subgroups, we examined whether they correspond to MAJ of Eastern European or Western European origin. While Western MAJ overlapped with Erfurt-ME, both MAJ groups were overall closer to Erfurt-ME than to Erfurt-EU (Figure S10). The Erfurt-ME group overlapped with present-day Turkish (Sephardi) Jews (Figure 1, inset). Erfurt-EU individuals had higher coverage, raising the concern that placement of individuals in PC space is coverage-dependent (Figure S11). However, down-sampling experiments demonstrated that coverage did not affect ancestry assignment (Figure S8). We also verified that there is no correlation between the proportion of European ancestry and the type of library preparation (Figure S8).

An *ADMIXTURE* [46] analysis demonstrated that EAJ are genetically similar to MAJ, but with higher variance (Figure S12), consistent with the PCA findings. Individuals classified based on the PCA as Erfurt-EU had higher EU-related ancestry. The results also revealed a small but consistent East-Asian-related component, especially in the Erfurt-EU group (means of 2.7% and 1.6% in Erfurt-EU and all EAJ, respectively), as previously observed [30]. This suggests either a minor gene flow event from East-Asia, as previously attested by mtDNA [47], or gene flow from Eastern European populations, who carry (at least today) a minor component of this ancestry (Figure S12).

We used f_4_-statistics to test for evidence of gene flow between EAJ, MAJ, and other EU and ME populations (Methods Section 4). We first ran tests of the form f_4_(MAJ, EAJ; X, chimp), where X is any West-Eurasian population. The results showed increasing Z-scores, and hence increased affinity with MAJ as opposed to EAJ, when X changed from Eastern European to Central/Western European, Mediterranean, and Middle Eastern (Figure S13). The same trend was observed for tests f_4_(MAJ, Erfurt-EU; X, chimp), but with the Z-scores being slightly lower. An opposite trend was observed for tests f_4_(MAJ, Erfurt-ME; X, chimp) (Figure S13). These results suggest, in agreement with the PCA and *ADMIXTURE* results, that MAJ have more EU ancestry (particularly Eastern-EU ancestry) than Erfurt-ME but less than Erfurt-EU. Tests of the form f_4_(Erfurt-EU, Erfurt-ME; X, chimp) showed increasing affinity with Erfurt-ME when X changed from Eastern European to Central/Western European, Mediterranean, and Middle Eastern (Figure S14). An Eastern European-related ancestry of some EAJ individuals may accord with recorded migration of families from Bohemia, Moravia, and Silesia into the second Erfurt community (SI 1) [48], assuming that Jews who came from the East have mixed with local populations during their residence there (SI 2). Finally, the tests f_4_(MAJ, X; EAJ, chimp) where X is any Jewish non-Ashkenazi population were positive and very large for all X, suggesting that EAJ are closer to MAJ than to other Jewish groups (Figure S15).

A *qpWave* analysis showed that EAJ and MAJ are consistent with forming a clade with respect to non-Jewish Europeans (P=0.15; Table S5; Methods Section 4). This genetic similarity between EAJ and MAJ, despite living about 700 years apart, suggests a high degree of endogamy over hundreds of years. Using simulations (Methods Section 5), we inferred that any hypothetical admixture event between AJ and Eastern Europeans in the past ≈20 generations must have been limited to replacing at most 2-4% of the total AJ gene pool (Table S6; this would correspond to at most 0.2% replacement per generation; Methods Section 5). The same *qpWave* analysis with Erfurt-EU or Erfurt-ME had lower P-values, particularly for Erfurt-EU, suggesting that each of these groups alone does not fully represent the entire modern AJ gene pool. When replacing EAJ or MAJ with (modern) Turkish Jews, South-Italians, and Germans, all P-values were under 0.05 (suggesting inconsistency with the given pair of populations being a clade), with the highest P-values observed when comparing Erfurt-ME and Turkish Jews (P≈0.01; Table S5). We also repeated the *qpWave* analysis to test whether pairs of populations form a clade with respect to non-Jewish Middle Eastern populations (Table S5). The P-values were >0.05 for tests comparing MAJ and EAJ/Erfurt sub-groups and either of these populations and Turkish Jews, suggesting that these populations have similar sources of Middle Eastern ancestry.

### Quantitative ancestry modeling

We used *qpAdm* to gain insight into the ancestral sources of EAJ (Methods Section 4). We modeled EAJ as a mixture of the following modern sources: Southern European (South-Italians or North-Italians), Middle Eastern (Druze, Egyptians, Bedouins, Palestinians, Lebanese, Jordanians, or Syrians), and Eastern European (Russians). To avoid bias due to ancient DNA damage, we only used transversions. Most of the models with a South-Italian source were plausible (P-value >0.05; Table S7), which would be consistent with historical models pointing to the Italian peninsula as the source for the AJ population. The mean admixture proportions (over all of our plausible models; Table S7) were 68% South-EU, 17% ME, and 15% East-EU (Figure 2A). However, the direct contribution from the Middle East is difficult to estimate given historical ME admixture in Italy [49] (see the Discussion). Indeed, a model with North-Italians as a source (which was only plausible with a Lebanese source; Table S7) had ancestry proportions 44% South-EU, 44% ME, and 12% East-EU (Figure 2A). We validated that the results did not qualitatively change when we tested the same models using all available SNPs, a different outgroup population, or fewer SNPs (Table S7, Figure S16). Models with a Western European source (Germans) instead of Russians were not plausible (Table S7), and there was no support for an East-EU-independent contribution of East-Asians (Methods Section 4). Interestingly, Erfurt-ME could be modeled based on Turkish (Sephardi) Jews (97% admixture proportion) and Germans (3%) as sources.

**Figure 2.**
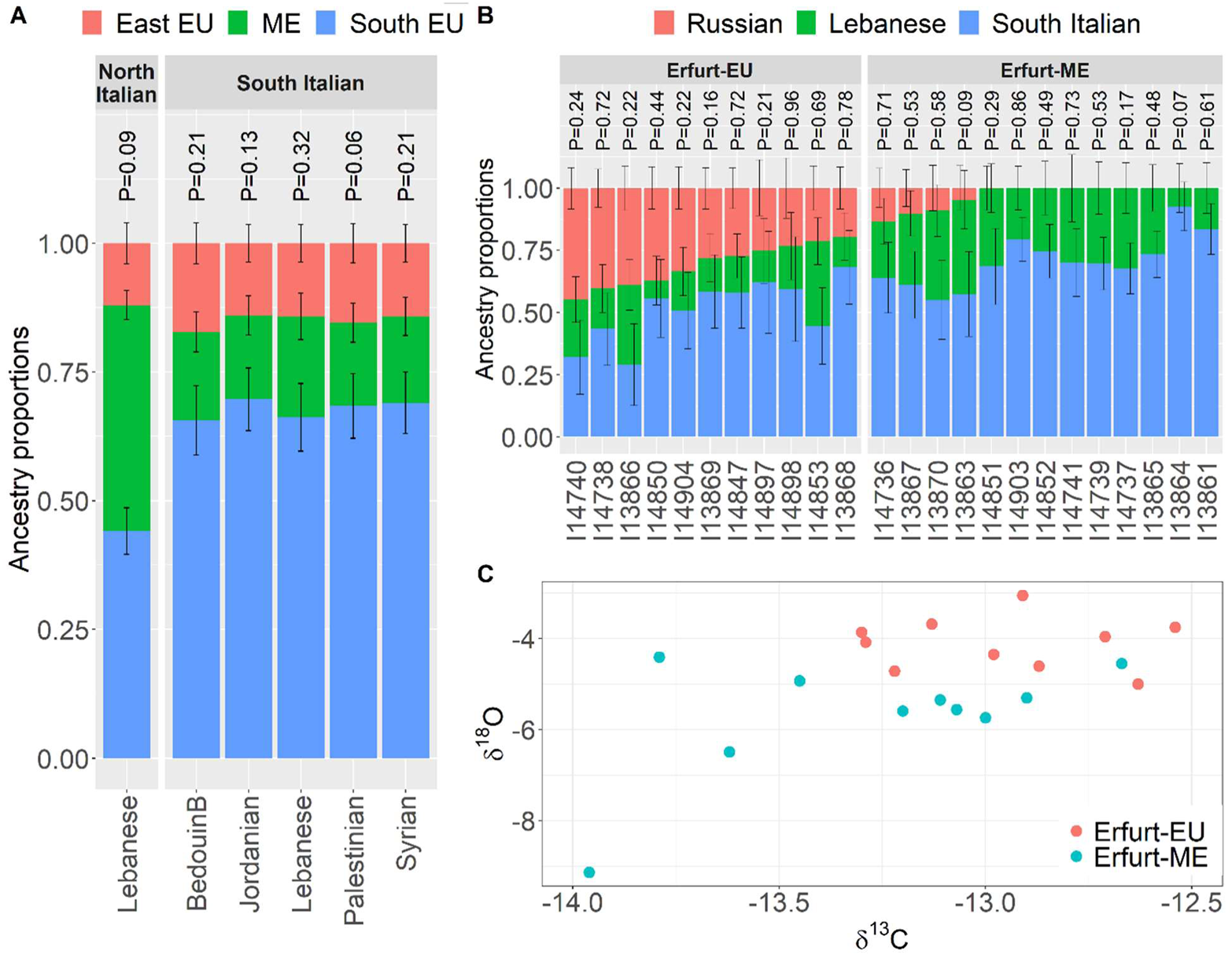
Models for the ancestry of Erfurt Ashkenazi Jews. (A) Each presented *qpAdm* model for the ancestry of Erfurt Jews includes a Middle Eastern, a Southern European, and an Eastern European source (Russians). The Southern European source was either South- or North-Italians, as indicated at the top of each panel. The Middle Eastern source is indicated in the x-axis labels. Only models with a P-value > 0.05 are shown (for full results see Table S7). Error bars represent one standard error in each direction. The P-values of the various models are presented above each model. (B) The ancestry of single Erfurt individuals, labeled by their IDs. We used *qpAdm* with Russian, Lebanese, and South-Italian sources. The individuals are labeled by their Erfurt subgroup (EU/ME). Results are not shown for low-coverage individuals (<50k SNPs), as well as for an additional individual who could not be modeled using these sources (P<0.05). (C) A plot of δ^13^C_enamel_ and δ^18^O_enamel_ isotope values for a subset of 20 Erfurt individuals with >200k SNPs. The Erfurt subgroup affiliation (EU/ME) is color-coded (legend).

We also tried to model EAJ as a mixture of ancient sources [50]. The sources we used were Imperial or late antique Romans [49], Canaanites [51], and early medieval Germans [52]. These models gave poor fit (P<0.01 for both Roman sources), suggesting a missing ancestry component. Alternatively, the poor fit might reflect technical artifacts due to inhomogeneous data types: for example, the Canaanite and EAJ datasets were produced by in-solution enrichment, while the Imperial/ late antique Roman and early medieval German datasets were produced by shotgun sequencing.

To quantify the variance in ancestry within EAJ, we used *qpAdm* to infer the admixture proportions for each EAJ individual in a model with South-Italian, Lebanese, and Russian sources (Figure 2B). We found striking variability in the Eastern European component, which was on average 33% in Erfurt-EU individuals, but was absent from 9 of 13 Erfurt-ME individuals. Similar variability was observed when using a North-Italian source in the *qpAdm* analysis (Figure S16C). This may be consistent with the recorded migration of Jewish families into Erfurt from the East (SI 1). Finally, a *qpAdm*-model suggested that MAJ have a 13% Eastern-EU (or 14% Western-EU) ancestry on top of that of Erfurt-ME.

We next hypothesized that the Erfurt individuals may provide information regarding the timing of admixture in AJ due to their proximity in time to the events. We attempted to estimate the time of Eastern European gene flow into Erfurt-EU using *DATES* [53] (Methods Section 5; Figure S17A). However, our simulations showed that *DATES* estimates in our setting were unreliable, possibly as the source populations are not sufficiently diverged (Methods Section 5; Figure S17B).

### Substructure in EAJ

EAJ have more variable ancestry than MAJ, which we interpreted above as the presence of two subgroups. However, the wider dispersion in EAJ could also reflect ongoing or very recent admixture [54]. Here, we investigate whether the dichotomization of the EAJ individuals is supported by the data. We used the first two PCs and clustered the EAJ individuals using *K*-means for several values of *K*. Based on the gap statistic [55] (Methods Section 6), the optimal number of clusters was *K*=2, providing statistical support to the existence of two sub-groups. As a control, the same method suggested that MAJ, as well as Moroccan Jews, form a single cluster each (Figure S18). The difference between the two EAJ clusters was also significant based on the approach of ref. [56] (P=0.007; Methods Section 6).

We next studied the number of EAJ groups using population genetic simulations (Methods Section 6). We mimicked a single group scenario by simulating admixture between Middle Eastern, Southern European, and Eastern European sources that occurred five generations prior to sampling. For the two-group scenario, we simulated admixture between Middle Eastern and Southern European sources 10 generations prior to sampling, and, five generations later, an admixture event from Eastern Europeans into a subset of the individuals. We ran PCA and *qpAdm* analyses on the simulated genomes under the two scenarios and compared the results to those of the real EAJ genomes (Figure S19, S20). The distribution of the PC1 coordinates and the inferred proportion of individuals without (*qpAdm*-inferred) East-EU ancestry in EAJ were similar to those simulated under the two-group scenario (Figure S19, S20; P=0.19 and P=0.78, respectively). The corresponding distributions under the single group scenario were different from those of EAJ (P=0.03 and P=0.014, respectively).

Altogether, our results provide support to the presence of two genetic sub-groups in EAJ, with one group having higher levels of Eastern European or related ancestry. Given the recorded migration of Jews from the East into the second Erfurt community (SI 1), we hypothesized that some EAJ individuals were migrants. To test this hypothesis, we performed an isotope analysis on dental enamel (Methods Section 1, Table S2, Table S8). The δ^13^C*_enamel_* and δ^18^O*_enamel_* values are plotted for all individuals in Figure 2C, showing distinct distributions of isotope values between the two genetic groups. The differences between the groups were significant for δ^18^O, although not for δ^13^C (P=0.0005 and P=0.1, respectively; two-tailed Wilcoxon test), suggesting average differences in water sources during childhood between the Erfurt-EU and Erfurt-ME populations.

We note that there was no correlation between the locations of the graves in the cemetery and the group affiliation (Erfurt-EU/ME; Mantel test P=0.46) or PC1 and PC2 coordinates (Mantel test P=0.41). This shows that even if these groups were genetically distinctive, they were not culturally or temporally segregated.

### Estimating a demographic model based on mtDNA sequence, IBD sharing, runs of homozygosity, and founder alleles

Previous analyses of identical-by-descent (IBD) haplotypes [19, 22, 23], mtDNA haplogroups [15], and pathogenic variants [11, 12] suggested that AJ have experienced a medieval founder event (a bottleneck). However, the demographic details of the bottleneck are yet to be fully resolved. Here, we used three sources of information — mtDNA haplogroups, runs of homozygosity, and MAJ-enriched variants — to determine whether the EAJ population has already postdated the founder event and estimate the bottleneck parameters.

We list the mtDNA haplogroups of EAJ in Table S2 and report the number of carriers of the four most common Ashkenazi haplogroups (carried cumulatively by about 40% of MAJ [15, 16]) in Table S9. Remarkably, among 31 unrelated individuals, 11 EAJ (35%) carried the K1a1b1a haplogroup. This is greater than the 20% frequency in MAJ (P=0.04; two-tailed binomial test; Table S9). All 11 carriers had a completely identical sequence, except a single C/T polymorphism at position 16223 (C count: 3/11). The same polymorphism also segregated in 107 MAJ K1a1b1a carriers (C count: 48/107; Methods Section 7). Excluding the 16223C/T site, 76/107 MAJ carriers had an identical sequence to that of EAJ. The remaining MAJ carriers were polymorphic at 36 additional sites (most of them (32/36) singletons). A joint Bayesian analysis [57] of MAJ and EAJ K1a1b1a carriers (accounting for the known date of the EAJ individuals; Methods Section 7) suggested a median posterior time to the most recent common ancestor of this haplogroup about 1500 years ago, slightly earlier than previous estimates [15, 16], although with very high uncertainty (95% highest posterior density: 650-6700; Figure S21).

Among the other AJ founder haplogroups, two EAJ individuals carried the K1a9 haplogroup, one carried N1b2, and none carried K2a2a1 (Table S9). The proportion of carriers of founder haplogroups was higher (though not significantly) in Erfurt-ME than in Erfurt-EU (7/13 vs 2/10; P=0.20; Table S9). Overall, the mtDNA results provide evidence that the Erfurt population has already experienced a bottleneck, perhaps even stronger than expected based on modern data.

To quantitatively estimate bottleneck parameters in EAJ and MAJ, we inferred demographic models based on IBD sharing and runs of homozygosity (ROH). Both IBD and ROH represent pairs of haplotypes that descend from a recent common ancestor, and are thus informative on the recent population size [58]. We started with a simple model of an ancestral population of a constant size that has experienced a bottleneck of size *N_b_* starting *T_b_* generations ago and lasting *d* generations, followed by an exponential expansion (Figure 3A). We first inferred the bottleneck parameters using modern IBD segment lengths and a maximum likelihood approach (Methods Section 8). We used whole-genome sequencing data for *n* = 637 individuals [19, 59]. The inferred bottleneck parameters were *N_b_* = 1563 (diploid individuals; 95% confidence interval (CI): [1364–1751]), *T_b_* = 41 (generations; 95% CI: [39–43]), and *d* = 20 (generations; 95% CI: [15–24]; see Figure 3A and Table S10, model (A)). [See Table S10, model (B) for the inferred parameters when fixing *d* = 1, as in previous studies [19, 23].] The predicted counts of IBD segments across length bins (based on the inferred model of Figure 3A) provide good fit to the observed counts (Figure 3B). Assuming 25 years per generation, our model would place the onset of the bottleneck about 1000 years ago, at the time of formation of the early Ashkenazi communities (SI 2).

**Figure 3.**
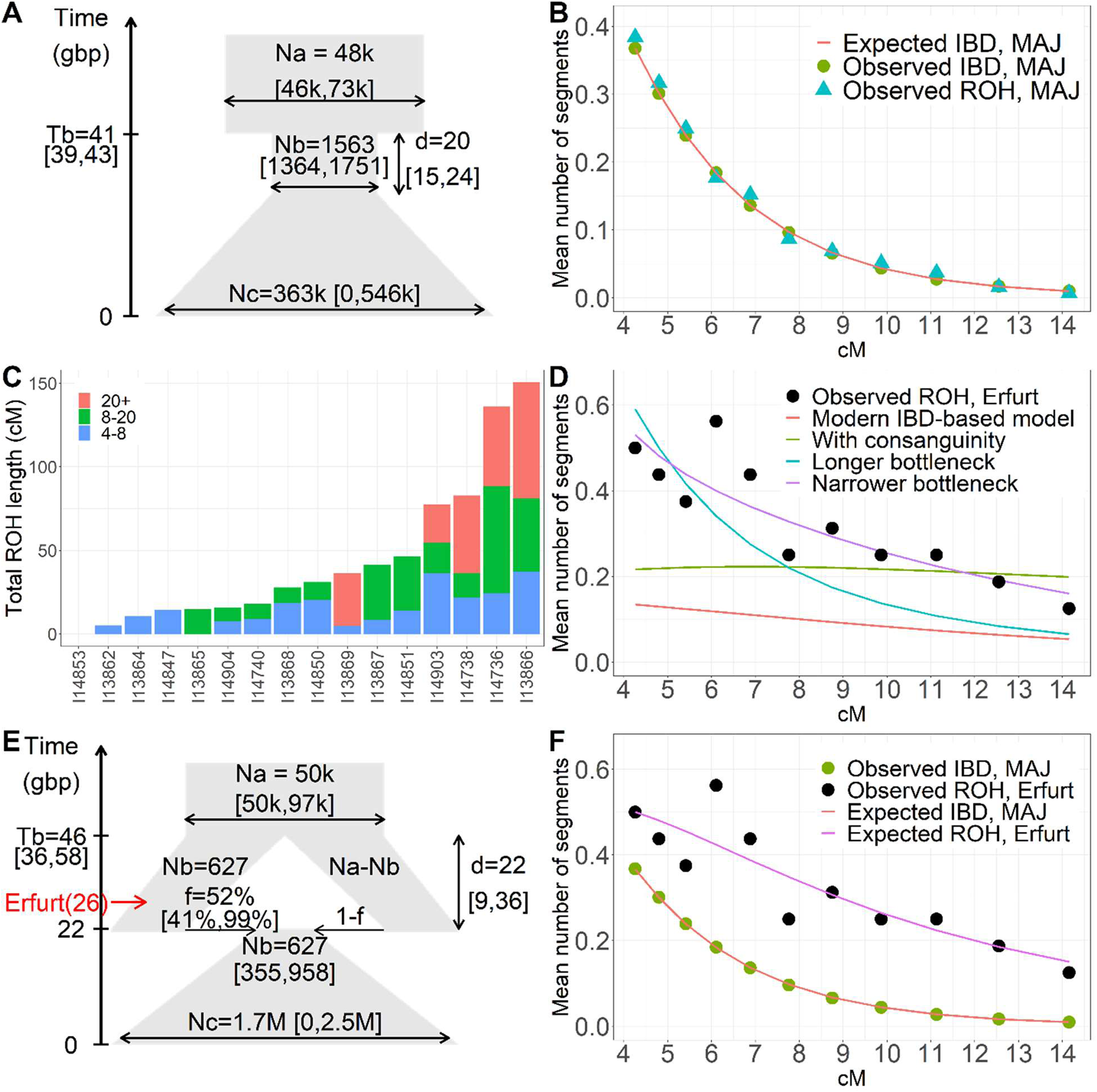
Models for AJ demographic history based on ancient and modern haplotypes. (A) A single-population model for the demographic history of AJ, inferred based on modern IBD sharing (Table S10, model (A)). According to the model, the effective population size has been constant (*N_a_* = 48*k* diploids) until *T_b_* = 41 generations ago. The population then contracted to *N_b_* = 1563 diploids, remaining in that size for *d* = 20 generations. At the end of the bottleneck, the population expanded exponentially until reaching effective size *N*_e_ = 363*k*. In the diagram, the y-axis represents the time in *generations before present* (gbp), and the width is (schematically) proportional to the effective population size. The 95% confidence intervals (CI) were computed using bootstrap and are indicated near each parameter. (B) The observed mean number of IBD and ROH segments (per pair of haploid genomes) in modern AJ across length bins (11 bins between 4-15cM). Each symbol (circles for IBD, triangles for ROH) is placed at the middle of its corresponding bin. The red line shows the expected number of segments per bin based on the demographic model of panel (A). (C) The total length of ROH segments in 16 EAJ individuals with at least 400k SNPs. The bars are colored proportionally to the contribution of segments of different lengths (legend). (D) The observed number of ROH segments in EAJ across length bins (circles) and the expected number based on various models (lines). The expectation based on the model inferred using modern IBD (panel (A)) is shown in red, and the expectation based on the same model but allowing consanguinity in EAJ (Table S10, model (D)) in green. Both models fit poorly to short ROH segments. We then plot the expectation based on a model similar to that of panel (A), but having either a longer or a narrower bottleneck (Table S10, models (E) and (F)) in teal and purple, respectively. These models fit the data better. (E) A two-population model inferred jointly using IBD in MAJ and ROH in EAJ (Table S10, model (H)). According to the model, the ancestral population of effective size *N_a_* = 50*k* split *T_b_* = 46 generations ago into one population of size *N_b_* = 627 (representing Erfurt, indicated in red) and another of size *N_a_* − *N_b_*. The bottleneck lasted *d* = 22 generations, and was followed by merging of the two populations with proportions *f* = 52% and 1 − *f*, respectively. The combined population then expanded exponentially as in the single-population model, reaching a present effective population size of *N*_e_ = 1.7*M*. The time of sampling of the Erfurt population is shown at 26 generations ago (assuming 25 years per generation). (F) The observed number of IBD segments in MAJ (green circles; same data as in panel (B)) and ROH segments in EAJ (black squares; same data as in panel (D)) across segment length bins, and the expectations based on the two-population model of panel (E) (MAJ: red line, EAJ: pink line). The two-population model fits the data well.

We detected ROH segments in the modern individuals and used them to infer the bottleneck parameters in a similar manner (Methods Section 8). The inferred model (Table S10, model (C)) was similar to that inferred based on IBD, and the counts of observed ROH segments fit well those predicted by the IBD-based demographic model (Figure 3B). We henceforth used the IBD-based model, which was based on more observations, and which did not rely on assumptions regarding consanguinity.

We next sought to determine whether our inferred demographic model provides a good fit to the ancient EAJ data (after accounting for the ≈650-year difference; Methods Section 8). We detected ROH segments in EAJ using a *hapROH* [60] (Methods Section 8; Figure S22). Due to the low coverage of the ancient genomes, we focused on 16 individuals covered in at least 400k SNPs. The EAJ individuals had substantially higher levels of ROH compared to other ancient populations (Figure S23), with an average of 44cM per individual in segments greater than 4cM (30cM in segments >8cM and 14cM in segments >20cM; Figure 3C). The total ROH length per genome was similar between Erfurt-EU and Erfurt-ME (P=0.43; two-tailed Wilcoxon test; Figure S24A). Interestingly, carriers of the K1a1b1a mtDNA haplogroup had higher levels of ROH compared to the other individuals (average per genome 76cM vs 25cM; P=0.03; one-tailed Wilcoxon test; Figure S24B). Overall, these results provide additional support to the hypothesis that EAJ have post-dated the bottleneck event.

To refine our inference of the bottleneck parameters, we compared the observed ROH counts in EAJ to those expected under the model we inferred using modern IBD (Figure 3A). The number of ROH segments in EAJ exceeded that expected based on the modern data, in particular for short and intermediate segments (Figure 3D). One possible explanation for the gap between the EAJ and MAJ data is a high rate of false positives for short ancient ROH segments. However, previous simulations argue against this hypothesis [60] (see also Figure S22), and we therefore attempted to identify models that fit both ancient and modern data. We first observed that a few EAJ individuals had very long ROH segments (five individuals with an average of 43.7cM in ROH segments of length >20cM; Figure 3C), which may result from their parents being related. We thus hypothesized that modeling consanguinity in EAJ may better fit the expectation based on the modern data (Methods Section 8; Table S10, model (D)). However, while this improved the fit for segments longer than 10cM, the observed number of short segments remained higher than expected (Figure 3D).

Our second hypothesis was that the Erfurt history involved a more intense or a more prolonged bottleneck compared to MAJ. We therefore tested whether a narrower or a longer bottleneck in the history of EAJ (compared to the model inferred based on modern IBD) could explain the gap in the number of ancient ROH segments (Methods Section 8). We found that a model with about 3.0-fold narrower bottleneck fit the EAJ ROH data (Figure 3D; Table S10, model (E)). However, this model did not fit the modern data (Figure S25A). The best-fit model with a (2.4-fold) longer bottleneck still deviated from both the ancient ROH and modern IBD data (Figure S25A; Figure 3D; Table S10, model (F)). To identify a model that would fit both ancient and modern data, we inferred the demographic parameters using both data types, giving equal weight to each (Methods Section 8; Table S10, model (G)). However, the inferred model did not provide a good overall fit to the modern data (Figure S25B). Taken together, our results suggest that the single-population model of Figure 3A is missing certain components of the AJ demographic history.

Motivated by our results regarding EAJ sub-groups, we hypothesized that a missing component in our model is substructure within AJ during its early history. We therefore attempted to fit the model shown in Figure 3E, in which the AJ population split during the bottleneck into two groups experiencing different bottleneck intensities, one of them represented by EAJ. According to the best fit model (Figure 3E and Table S10, model (H)), the bottleneck started *T_b_* = 46 generations (95% CI: [36, 58]) ago and lasted *d* = 22 generations (95% CI: [9, 36]). The group represented by EAJ experienced a narrow bottleneck of size *N_b_* = 627 (95% CI: [355, 958]) and contributed 52% (95% CI: [41, 99]%) to the MAJ gene pool. The remaining contribution came from a second group that did not experience the initial bottleneck but contracted to size *N_b_* only *T_b_* − *d* generations ago (Figure 3E). This model fits well both modern and ancient data (Figure 3F). We used parametric bootstrap to show that rejection of the single-population model is not due to overfitting (P<0.01; Methods Section 8).

We were able to show by simulations that our method can infer the parameters of a two-population model (Figure 3E) even in the presence of extreme imbalance between the amount of modern and ancient data (Methods Section 8). However, even this expanded model relies on several simplifying assumptions: that the non-Erfurt sub-group experienced no initial bottleneck; that there was no gene flow from non-AJ populations or between the AJ sub-groups; that population splitting and merging coincided with the start and end of the bottleneck, respectively; and that the bottleneck remained of a constant size throughout its duration. Thus, our results should not be interpreted as a statement on the correct form of the demographic model, and we cannot rule out alternative models, particularly more complex ones. Nevertheless, our results illustrate how a model of substructure, with different groups undergoing different bottleneck intensities, can reproduce the modern and ancient haplotype sharing data.

The third source of information regarding the bottleneck is variants that are specific (or near-specific) to MAJ and are also present in EAJ. We define AJ *founder alleles* as minor alleles (not necessarily disease-causing) in SNPs targeted by our in-solution enrichment that have frequency >0.5% in MAJ, frequency <0.01% in Europeans (both in *gnomAD* [61]), and frequency ≲ 1% in the Middle East (Methods Section 8). Overall, we identified 216 AJ founder alleles. Among these alleles, 15 were present in at least one EAJ individual (Table S2). To determine whether this number of observed alleles is expected under the scenario that EAJ post-dated the bottleneck, we used MAJ allele frequencies and ran binomial simulations to estimate the expected number given the EAJ sample size and per-individual coverage (Methods Section 8). The [2.5,97.5]-percentiles of the simulated allele counts in EAJ-like individuals were [14, 32] (Figure S26). Those percentiles are likely overestimated due to the conditioning on alleles whose MAJ frequency was above a cutoff (Methods Section 8). The presence of 15 founder alleles in EAJ is thus within the range expected if EAJ has already experienced the AJ bottleneck.

The proportion of individuals who carry AJ founder alleles was similar between Erfurt-EU and Erfurt-ME (44% and 46%, respectively). The proportions remained similar even when accounting for coverage (P=0.45; quasi-Poisson regression; Methods Section 8). In contrast, the proportion of founder allele carriers was higher in K1a1b1a mtDNA carriers compared to other individuals (73% vs 17%; P=0.005; Fisher’s exact test; although the evidence weakened when accounting for coverage (P=0.038)). Thus, as also implied by the ROH data, K1a1b1a carriers may have been particularly affected by the bottleneck.

### Pathogenic variants

If EAJ were affected by the same founder event as MAJ, we expect EAJ to carry some of the pathogenic variants common in present-day AJ. We compiled a list of 62 pathogenic variants based on [19], after excluding variants with high frequency in Europeans and East-Asians (Methods Section 9; Table S11). However, as our dataset is based on SNP targeting, only six variants were enriched. Imputation based on whole-genome sequences can be error-prone for ancient populations that are genetically distinct from any modern population. However, as we showed above, EAJ are genetically close to MAJ, and we therefore attempted to impute the EAJ genomes based on a reference panel of 702 MAJ whole-genomes [19, 59].

As current imputation software cannot handle pseudo-haploid input, we developed an imputation method based on the *PHCP* framework [51] (Methods Section 9; see also [62]). Briefly, *PHCP* (Pseudo-Haploid *ChromoPainter*) uses the Li-Stephens hidden Markov model (HMM) [63], as implemented in *ChromoPainter* [64], but with the hidden state representing a pair of haplotypes from the reference panel. As in other imputation methods, transitions are due to ancestral recombinations, and the emission probabilities represent the possibility of imperfect copying from the reference haplotypes. The imputed diploid genotypes are based on the posterior marginal probabilities of the HMM.

We validated the *PHCP*-based imputation results using three approaches. First, we calculated the proportion of SNPs with Mendelian inconsistency in the three children of families A and B (Methods Section 9). The proportion of inconsistent SNPs was 0.19-0.23% for the two children who were covered in >500k SNPs, and 0.59% for the child with 113k SNPs (Table S12). In comparison, the inconsistency rate was 2.13-2.15% in unrelated individuals (Table S12). Second, we masked the 216 founder alleles (and three pathogenic variants), imputed them, and evaluated the imputation accuracy (Methods Section 9).

Across the 219 SNPs and all EAJ individuals, in 20 cases the alternate allele was genotyped. After masking, an alternate allele was imputed in 15 cases, implying a false negative rate of 5/20=25%. To estimate the false-positive rate, we considered 2541 cases where the (pseudo-haploid) genotype was the reference allele. After masking, the alternate allele was imputed only 13 times. In some of these cases, the alternate allele may have been truly present but not reported. Therefore, 13/2541=0.51% is a conservative estimate of the false positive rate. Finally, we imputed the EAJ genomes using *GLIMPSE* [65] and compared the results to *PHCP*. This approach does not provide a fully independent validation, as *GLIMPSE* is also based on the Li-Stephens model and we used the same MAJ reference panel. However, its implementation is distinct from that of *PHCP*. Among pathogenic variants confidently imputed by *PHCP* (see below; Table 1; Table S11; Methods Section 9), only 1/7 was missed by *GLIMPSE*. In the other direction, 0/6 variants confidently detected by *GLIMPSE* were missed by *PHCP*. Overall, these results support the reliability of our imputation framework.

In the imputed EAJ genomes, we discovered with high confidence 11 Ashkenazi pathogenic variants. These variants were either genotyped or imputed with marginal posterior probability >97% in *PHCP* and >50% in *GLIMPSE* (Table 1). Five other variants were detected with low confidence (Methods Section 9; Table S11). The high confidence variants were carried by eight EAJ individuals, with each variant appearing once (Table S13). Seven carriers belonged to Erfurt-ME and four carried the K1a1b1a mtDNA haplogroup (Table S13).

Several of the discovered variants are of notable medical importance, and some were dated using modern genomes. We discovered two dominant variants. The *BRCA1* 185delAG (also known as c.68_69delAG) is one of three common variants in *BRCA*1/2 genes in AJ [6] and is known to increase the lifetime risk of breast and ovarian cancer by more than 70% and 44%, respectively [66]. It is also present in Iraqi and other Jews, and was previously dated to 61 generations ago [67]. The G2019S variant on *LRRK2* (which was genotyped) increases the risk of Parkinson’s disease [68]. The variant is common in North-Africans, and it was found in about 20% and 40% of Parkinson’s disease cases in AJ and North-Africans, respectively. The variant was previously dated to a few thousands of years ago [69, 70]. The remaining variants were recessive. The 84dupG variant in *GBA* is an AJ-specific variant for Gaucher disease (along with the more common N370S variant). It was previously dated to 56 generations ago [71]. Two variants on *F11*, leading to Factor XI deficiency, were genotyped. The type II variant (E135X; also known as E117X) is present in other Jewish and Arab populations [72] and was previously dated to 120-189 generations ago [73]. The Type III variant (F301L; also F283L) is AJ-specific and was dated to at least 31 generations ago [73]. The familial Mediterranean fever *MEFV* variant V726A is found in multiple Middle Eastern populations and is dated to a few thousands of years ago [74–76]. The discovery of all of these variants in 14^th^-century EAJ is consistent with previous estimates of their time of origin.

Other recessive pathogenic variants we detected include the cystic fibrosis *CFTR* variant G542X [77], the retinitis pigmentosa *DHDDS* variant 124A>G [78], the Usher syndrome (type 3) *CLRN1* variant N48K [79], and the glycogen storage disease (type 1A) *G6PC* variant R83C [80]. All of the above recessive variants are in genes included in pre-conception screening panels (Table 1). Finally, we identified a female child carrier of the *ACADS* c.319C>T variant, who had a 44% probability to be homozygous (Table S11) and thereby affected by acyl CoA dehydrogenase deficiency. While the disease may be associated with failure to thrive [81], we did not notice any pathologies in her skeleton that might be associated with this phenotype. An important caveat of this analysis is that imputation demonstrates the presence of the *haplotypes* carrying the variants, but the variants themselves may have been missing from EAJ. However, the low false positive rate we observed above with genotyped founder alleles suggests that such a scenario should be rare.

### Other phenotypes

The lactase persistence dominant allele rs4988235/T [82] is known to have a much lower frequency in MAJ compared to Europeans (10.0% vs 60.1%, respectively; *gnomAD*; Table S11). The T allele frequency in (unrelated) EAJ was 11.7% (7/60; 95% CI: [6, 22]%; Methods Section 10; Table S11), similar to the MAJ allele frequency. The blue eye recessive allele rs12913832/G [83] had frequency 55% in EAJ (33/60; 95% CI: [42, 67]%; Table S11), again similar to the MAJ frequency (54.8%). The red hair recessive alleles rs1805007/T, rs1805008/T and rs1805009/C [84] were present in 8.3% of EAJ (5/60; 95% CI: [3.6,18.1]%, Table S11) compared to 12.4% in MAJ. No homozygous carriers were observed.

To test the ability of polygenic scores to predict stature in EAJ, we used osteological estimates of height in 13 adults (Methods Section 1; Table S2, Table S14). These estimates were correlated (*r* = 0.48, 95% CI: [−0.10,0.81]; Figure S27) with genetically-predicted heights (Methods Section 10). While our sample size is too small to reach a definitive conclusion, the ability to genetically predict the stature of ancient individuals, even if at reduced accuracy, agrees with recent studies [85, 86]. Finally, a recent study [87] found a sharp change in allele frequency for rs17514136 and rs10839708 between 16^th^-century plague victims in Ellwangen, Germany, and modern individuals from the same town. In AJ, allele frequencies were similar between 14^th^-century EAJ and MAJ (P=0.18 and 0.81, respectively; one-tailed binomial test; Table S11).

## Discussion

We have presented the first genome-wide data from historical AJ individuals. We used the data to refine the picture of early AJ origins. The ancestry of EAJ was closely related to that of modern AJ, as evidenced by the *PCA*, *ADMIXTURE*, and *qpWave* analyses, suggesting overall genetic continuity of AJ over the past ≈700 years. However, EAJ individuals had more variable ancestry than MAJ and were possibly stratified by the presence of a minor Eastern European ancestry component. Multiple lines of evidence suggest that the EAJ population had already experienced a “bottleneck” shared with MAJ: the high frequency of Ashkenazi founder mtDNA haplogroups; and the presence of Ashkenazi-specific pathogenic variants, other AJ-enriched alleles, and long runs of homozygosity. Carriers of the K1a1b1a mtDNA founder haplogroup seem to have descended from an even smaller set of founders. In agreement with previous studies [19, 23, 25], we date the onset of the expansion in AJ population size to about 20-25 generations ago (see additional discussion in SI 2).

Our ancient DNA data allowed us to identify patterns in the history of AJ that would not have been otherwise detectable from modern genetic variation. Specifically, our genetic results suggest that the AJ population was structured during the Middle Ages. Within Erfurt, one group of individuals had an enrichment of Eastern European-related ancestry (Figure 1 and Figure 2B), while the other had ancestry very close to that of MAJ of Western European origin and modern Sephardi Jews (Figure 1 and Figure S10). The two groups also had significantly different levels of enamel δ^18^O (Figure 2C). Medieval AJ may have been structured even beyond Erfurt, based on our inferred demographic model (Figure 3E). In contrast, present-day AJ is a remarkably homogeneous population [17, 23, 33]. This suggests that even though the overall sources of ancestry remained very similar between medieval and modern AJ, endogamy and within-AJ mixture since medieval times have contributed to the homogenization of the AJ gene pool.

We found that a plausible model for the ancestral sources of EAJ (Figure 2A) include groups related to people in South-Italy (about 70%, who themselves plausibly might harbor Middle East-related ancestry), the Middle East (about 15%), and Eastern Europe (about 15%). Models with North-Italians as a source were also plausible, with an ancestry proportion of about 45% to each of North-Italians and Middle Easterners. The ancestry proportion estimates using North-Italians are closer to previous estimates using modern SNP and sequencing data [19, 35], but a North-Italian source was less favored by *qpAdm* (Table S7). While these results could be consistent with a model where the Middle Eastern ancestry in AJ has not been as large as previously thought, complicating the picture are (i) our inability to identify a satisfactory model for modern AJ; (ii) the historically variable levels of Middle Eastern ancestry in Italy [49, 88–90] (SI 2); and (iii) the possible problems when modeling an ancient population with modern sources using *qpAdm* [50] (although see our robustness tests in Table S7). Therefore, the direct contribution of ME sources to AJ ancestry may be higher than estimated (SI 2). Either way, the substantial Southern European ancestry we inferred adds weight to the evidence that early AJ descended, at least partly, from Italian Jews (SI 2). The estimate of about 15% Eastern European-related ancestry is consistent with a previous study [35]. The identification of this source as Eastern European relies on the f_4_ results (figs. S13, S14) and the *qpAdm* models (Table S7); however, this ancestry might derive from a broad area across Central or Eastern Europe, which may accord with recorded migration into Erfurt from Bohemia, Moravia, and Silesia (SI 1). For an additional discussion on the historical interpretation of these results, see SI 2.

As with other ancient DNA studies, our historical inferences are based on a single site in time and space. This implies that our data may not be representative of the full genetic diversity of early AJ, as we have indeed inferred (Figure 3E). However, even for a single site, our sample size was relatively large (>30), and we were able to capture substructure not present in MAJ. Another limitation is the reliance of our demographic models on a relatively small number of runs of homozygosity, which are difficult to infer from pseudo-haploid data. In particular, several models were disqualified due to mismatch with observed counts of short ROH or IBD segments, which are difficult to accurately call (Figure 3D; Figure S25). Our inferred demographic model (Figure 3E) should not be interpreted as a complete and precise demographic reconstruction; rather, it should be viewed as a simplified model (perhaps one among many) that captures the main features of the observed genetic data. Similarly, our models for the ancestry of EAJ (Figure 2A) may not be the only plausible models, and the ancestral sources we inferred should be interpreted as proxies, distant in time and space, of the true ancestral populations.

Our radiocarbon dating definitively timed the EAJ individuals to the 14^th^ century. However, it did not resolve whether the section of the cemetery that we studied was used before or after the 1349 pogrom. Skeletal evidence for a violent cause of death was observed only in one EAJ individual, implying that the individuals we studied were unlikely to be victims of the 1349 events. As we found no traces of *Yersinia pestis* in the DNA (Table S4), and given that the Black Death has arrived in Erfurt only in 1350 [42], the plague is also not likely to have been a major cause of death. The historical records on the migration of families from the East into the second Erfurt community (SI 1) [48] and the presence of EAJ individuals with increased Eastern European ancestry provides evidence in favor of the EAJ individuals belonging to the second community. Such an inference would be consistent with archaeological evidence and the isotope analysis (Figure 2C).

The discovery of pathogenic Ashkenazi founder variants in the EAJ genomes sheds light on their origins. The background haplotypes of six variants – in the genes *BRCA1*, *LRRK2*, *GBA*, *F11* (two variants), and *MEFV* – were previously studied. For all six variants, the estimated coalescence time of the haplotypes of carriers was >25 generations ago, in agreement with the haplotype already being present in EAJ. We detected additional variants of medical importance in the genes *CFTR*, *DHDDS*, *CLRN1*, and *G6PC*; a finding that provides important information on the natural history of these variants.

## Acknowledgements

We are thankful to Ephraim Shoham-Steiner for consultations when planning this study, and to Rabbi Ze’ev Litke, who undertook a ruling on contexts in which ancient DNA studies in Jews could be permissible under strict rabbinical Jewish law. We thank the Jewish community of Thuringia for considering and approving the study. We thank Douglas Kennett, Nick Patterson, and Maria Stürzebecher for discussions; Shaul Stampfer, Sergio DellaPergola, and Itsik Pe’er for commenting on the manuscript; Kim Callan, Fatma Zalzala, Kristin Stewardson, Nicole Adamski, and Ann Marie Lawson for wet laboratory work; Rebecca Bernardos for sample management; and Iosif Lazaridis, Matthew Mah, Adam Micco, and Zhao Zhang for bioinformatics support. The study was funded by the Israel Science Foundation grant 407/17 and the United States–Israel Binational Science Foundation grant 2017024 to SC, by the National Science Foundation (USA) grants 1912776 and 0922374 to VR, and by the following grants to DR: NIH grants GM100233 and HG012287; the Allen Discovery Center program, a Paul G. Allen Frontiers Group advised program of the Paul G. Allen Family Foundation; John Templeton Foundation grant 61220; a private gift from Jean-François Clin; and the Howard Hughes Medical Institute.

This article is subject to HHMI’s Open Access to Publications policy. HHMI lab heads have previously granted a nonexclusive CC BY 4.0 license to the public and a sublicensable license to HHMI in their research articles. Pursuant to those licenses, the author-accepted manuscript of this article can be made freely available under a CC BY 4.0 license immediately upon publication.

## Data availability

We will post the genotype data for the ancient Erfurt individuals upon publication at https://reich.hms.harvard.edu/datasets and the sequencing data at the European Nucleotide Archive. All other ancient genomes and the Human Origin array data for modern populations are available at https://reich.hms.harvard.edu/allen-ancient-dna-resource-aadr-downloadable-genotypes-present-day-and-ancient-dna-data. The modern Ashkenazi Jewish genomes are available at https://ega-archive.org/datasets/EGAD00001000781. We will post the isotope data at Isobank upon publication.

## Code availability

The *PHCP* imputation method is available at https://github.com/ShamamW/PHCPImpute. All other software packages used in this study were previously published.

## Author contributions

SC and DR conceived, designed, and supervised the study. KS performed the archaeological excavation. SF performed the physical anthropology analyses. DB and JCN performed the mitochondrial DNA analysis. NCN, GB, KMP, and VR performed the isotope analysis. ML provided details on the history of the Erfurt community. LVR provided historical context on Jewish origins. NR supervised the laboratory experiments. SM supervised the bioinformatics analyses. AA performed imputation using *GLIMPSE*. ÉH performed the pathogen DNA analysis. HR performed the detection of the runs of homozygosity. HF provided information on carrier screening. SW performed all other data analyses. HO, NB, GA, IP, and TL provided the Ashkenazi genomic reference data. JY performed quality control on the reference data. YEM provided input on the mtDNA analysis. SS provided input on the population genetic analyses. SW and SC drafted the manuscript with input from other authors and revised it together with DR. HR, HO, TL, LVR, and SS provided extensive feedback on the manuscript.

## Methods

### 1. Archaeology and physical anthropology of the Erfurt site

#### 1.1. The archaeological excavation

Traditional Judaism imposes very strict regulations on the management of Jewish cemeteries, including the directive that the dead should be left in peace—relocation is possible only under very particular circumstances. As a result, archaeological investigations in areas where Jewish cemeteries have survived (or are suspected to exist) are not allowed as a matter of principle. While we know of quite a few medieval Jewish cemeteries in Europe, only few have been excavated properly [1, 2]. All published examples are the result of rescue excavations. In most cases, the human bones were reburied as quickly as possible, in consultation with the Jewish communities. The excavations in Erfurt took place under similar circumstances.

The medieval Erfurt Jewish cemetery was located, following religious regulations, outside the city of Erfurt itself. It is unknown when the excavated section of the cemetery was used for burial by the Erfurt Jewish community. However, some hints arise from examining the fortifications around the site and from archaeological evidence from elsewhere in the city. Our excavated section is located between the first city wall (12^th^ century) to the south and an outer wall to the north, in an area where a moat used to lie in front of the first city wall (Figure S1; Figure S2). It is conceivable that the excavated section was used as a cemetery only after the construction of the outer wall, as prior to constructing that wall, the area was used for fortification and the original cemetery must have extended beyond the outer wall to the north. In Brühl, a site in the western part of Erfurt, wood retrieved from moat in front of the first city wall was dated by dendrochronology to 1324/1325. Several years later – at an unknown date – the moat was filled up and a second fortification wall was built with a new moat in front of it. It is plausible that the construction of an outer wall in the Jewish cemetery and in Brühl happened at around the same time. We consequently hypothesize that the outer wall in the area of the Jewish cemetery was likely constructed only in the second half of the 14^th^ century. If correct, this would imply that the excavated section of the cemetery was used only by the second community, during the second half of the 14^th^ century. Indeed, radiocarbon dating of the teeth we sampled dated them to the 14^th^ century (Table S2; Table S3). However, the radiocarbon results could not exclude origins in the first half of the 14^th^ century or even slightly earlier, which would place the samples in the first community (Figure S6; Figure S7).

After the expulsion of the Jews from Erfurt in 1454, a barn and a granary were built by the city in the years 1465-1473 on top of the cemetery. The granary (Kornhofspeicher) still exists today. The southern and northern walls of the granary were constructed on top of the inner and outer city walls, respectively (Figure S1A). In earlier investigations of the area surrounding the granary (over a period of several years), numerous gravestones and human bones were recovered. Burials *in situ* (in graves) were only observed in the area a little further north of the outer wall. These could not be recovered due to safety reasons.

The conversion of the granary into a multi-story car garage in 2013 required the construction of an external ramp, which then necessitated an archaeological rescue investigation. The excavation was carried out between 8 March and 17 April 2013. Since graves were not recognizable, a planum was first laid out by machine, at which point wood remains and some bones started to become visible. From then on, excavation was only done by hand. In the course of the excavation, at the suggestion of the builder, the recovery of the skeletons was restricted to areas where the constructions were expected to destroy the graves. In other areas, the skeletons remained in the ground.

The number of graves exposed during the excavation is likely a small fraction of the total in the cemetery. The size of the excavation area was about 16 x 12 meters. It is certain that the cemetery continued to the west, up to an unknown boundary. To the east, burials were partially destroyed by the construction of the granary. The ground level inside the granary is so low that its construction in the 15^th^ century destroyed all burials along a length of more than 80 meters. If one assumes an overall occupancy of a similar density as in the excavation field, about 1000 graves were destroyed by the construction of the granary. This assumes burial only on one level, as was encountered in the excavation area.

The archaeological documentation includes 47 burials. Six further graves were documented only after construction was underway and could only be partially recovered. Remains of wooden coffins were found in almost all graves. Tombs were located remarkably close to one another (Figure S2), and followed medieval Jewish funerary practice in that the integrity of tombs is always preserved. With one exception (I14850), all the burials lay parallel to the city wall, with the legs of the interred pointing roughly to the east – i.e., roughly in the direction of Jerusalem (Figure S2). No grave goods were observed, but there is evidence that some of the dead were buried with their clothes. This is suggested by the presence of buckles (I14904, who was violently killed), a piece of jewelry on one of the women (I14850 again; the piece has a close parallel in the treasure trove from Weißenfels from 1349), and a silk ribbon on the head of a child. A full report detailing the physical anthropology of the remains will be published shortly [3].

In 2018, we (K.S. and S.F.) collected detached teeth (mostly molars) for the DNA study. Overall, we found 38 teeth, one per individual (see teeth numbers in Table S2). In 2021, all skeletons were reburied in the recently recovered Jewish cemetery of the 19^th^-century community. This cemetery has signs that explain the history of the cemetery and provide information about the reburied Medieval dead.

A full description of the site, including an excavation report, will appear in the *Die mittelalterliche jüdische Kultur in Erfurt* (volume 6).

#### 1.2. Age at death estimation

We estimated the age at death for non-adult individuals by assessing the developmental stage of the dentition [4] and the length of the long bones [5]. In adult individuals, we estimated the age at death by established methods based on the stage of degeneration of the pubic symphysis face, the auricular joint face, the sternal rib ends, and additionally the cranial suture closure, as summarized e.g., in [6]. Adult age was determined in an individual when the epiphyses were fused.

#### 1.3. Height estimation

We reconstructed body height for two individuals using the anatomical method [7], and compared the results with those obtained using the mathematical method [8] based on several long bone measurements. As the results matched well, we used Pearson’s regression formulae to reconstruct body height in all adult individuals whose long bones were sufficiently preserved. When comparing the estimated height to the genetic data, we used the mean over all estimates of each individual.

#### 1.4. Radiocarbon dating

We performed accelerator mass spectrometry radiocarbon dating on purified collagen extracted from tooth dentin at the Pennsylvania State University Accelerator Mass Spectrometry (AMS) laboratory using previously described methods [9, 10]. We calibrated dates using OxCal 4.4 [11] and the IntCal20^48^ calibration curve [12]. We assessed sample quality using stable isotope analysis. We found that carbon-to-nitrogen ratios for all collagen samples fell between 3.19-3.24, which is well within the range of 2.9-3.6 expected for good collagen preservation [13].

The estimated dates are reported in Table S2 (95.4% probability intervals) and the full measurements are reported in Table S3. Our first attempt at dating the father and daughter from Family B (I14904 and I13869, respectively) yielded results inconsistent with their relationship (95.4% CI 1266-1298 calCE for the daughter and 1288-1398 calCE for the father). We radiocarbon dated these samples again, obtaining the same result for the father (1297-1395 calCE). For the daughter, the second run suggested a range of dates consistent with the relationship (1278-1380 calCE), due to the appearance of a small post-1350 peak (Figure S6).

We note that we dated teeth, which form in childhood and early adulthood. Thus, all dates should not be interpreted as representing date of death, but instead a date during an earlier time of life.

#### 1.5. Isotope analysis

To test the hypothesis that some Erfurt individuals were migrants, we selected enamel samples for δ^13^C and δ^18^O isotope analysis at the University of New Mexico Center for Stable Isotopes. Among samples with sufficient material, we considered 20 samples with coverage >200k SNPs, which were confidently assigned to an Erfurt subgroup (EU/ME).

Enamel surfaces were cleaned with a rotary tool and a 200µm endmill used to remove about 30 mg of enamel, avoiding any dentin. Samples were crushed in an agate mortar with UltraPure water (resistivity=18.2 MΩ cm) to keep chips from shattering. About 15 mg of sample was transferred to a 2ml centrifuge tube with about 0.1M Acetic Acid and agitated for about 30 seconds using a VortexGenie to sufficiently mix the sample and solution. Samples were reacted for about 4 hours, then rinsed to neutrality with UltraPure water and microcentrifuged for 1 min at 3000 rpm between each rinse. After the final rinse samples were freeze dried overnight (about 12 hours). Next, 7-8 mg of sample was weighed into 12ml glass exetainers for analysis. An additional 1mg aliquot of sample is separated and placed on the ThermoFisher Nicollet Summit FTIR diamond ATR for QC/QA analysis. Samples were analyzed on a Thermo Scientific Delta V IRMS with a GasBench carbonate device for δ^13^C_en_*_a_*_mel_ and δ^18^O_en_*_a_*_mel_.

The full results of the isotope analysis are reported in Table S8, and the final values of δ^13^C_en_*_a_*_mel_ and δ^18^O_en_*_a_*_mel_ appear in Table S2.

### 2. DNA sequencing

#### 2.1. DNA extraction and sequencing

Of the 38 teeth, four did not have root material and were not further processed. For the remaining 34 samples, we performed the following. In dedicated clean rooms at Harvard Medical School, we drilled into the roots of the teeth to obtain powder from cementum and dentin. We extracted DNA using a protocol meant to retain short molecules [14]; converted the DNA into individual barcoded and/or indexed double-stranded and single-stranded libraries treated to remove characteristic ancient DNA damage [15–18]; enriched for approximately 1.24 million single nucleotide polymorphisms (SNPs) [19] and mitochondrial DNA [20]; and sequenced the enriched and non-enriched libraries on Illumina instruments (Table S1).

#### 2.2. Bioinformatics and quality control

We merged read pairs, requiring at least 15 overlapping base pairs, and allowing no more than one mismatch if base quality was at least 20, and as much as three mismatches if base quality was less than 20, and chose the nucleotide of higher quality when we observed mismatches. We mapped the sequences to the human genome reference sequence hg19 (GRCh37, https://www.ncbi.nlm.nih.gov/assembly/GCF_000001405.13/) and the inferred mitochondrial ancestral sequence RSRS [21] using the *samse* command of *BWA* version 0.7.15 using parameters -n 0.01, -o 2, and -l 16500 [22]. We removed duplicate molecules that mapped to the same start and stop positions and (for double-stranded libraries) that had the same molecular barcodes.

We analyzed the data to assess ancient DNA authenticity based on the following metrics (Table S1). First, the ratio of Y to X+Y chromosome sequences: uncontaminated females should have a very low (<0.030) and males should have a high (>0.35) ratio in the type of data we produced. Second, the estimated rate of variation in the mitochondrial genome and the X chromosome in males at known polymorphisms: uncontaminated individuals should be consistent with very little variation. We estimated the degree of contamination with *contamMix* version 1.0-12 [23] for the mitochondrial DNA and *ANGSD* for the X chromosome [24]. Third, an appreciable rate of cytosine to thymine damage in the final nucleotide, as expected for genuine ancient DNA. We initially obtained DNA data for 34 samples. However, one sample was covered only in 52 SNPs, and was omitted from all analyses.

We determined the mitochondrial haplogroups using *HaploGrep2* [25]. The procedure for Y-chromosome haplogroup determination is described in Supplementary Text S7 of [26], using the YFull YTree v. 8.09 phylogeny (https://github.com/YFullTeam/YTree/blob/master/ytree/tree_8.09.0.json), obtaining information about SNPs from ISOGG YBrowse (https://ybrowse.org/gbrowse2/gff/snps_hg38.csv; accessed Oct 18, 2020), lifting coordinates from hg38 to hg19 using liftOver and intersecting with the SNPs present in the v. 8.09 tree. The haplogroup calls were converted into letter-number haplogroup designations (e.g., J2a1) using the phylogeny of the International Society of Genetic Genealogy v. 15.73. We could not infer the haplogroup of four males due to insufficient coverage over the informative Y-SNP targets.

#### 2.3. Lower coverage in children

We initially observed that all EAJ samples covered at <100k SNPs were under the age of 13. To formally test whether children had lower coverage, we used the mid-range of the estimated age at death (see Methods 1.2) and classified all individuals of estimated age ≤20 as children and all others as adults. We excluded two samples whose ages were not estimated. We then used a two-tailed t-test to compare the number of covered SNPs between children and adults. The significantly lower coverage in the children raises the possibility that DNA may be less well preserved (on average) in teeth that are not fully developed.

#### 2.4. Detecting relatives

We used the method reported in refs. [9, 27, 28] to identify first-degree relatives. To detect additional relatives, we used *READ* [29] with the default parameters.

#### 2.5. Pathogen DNA scan

We screened the Erfurt individuals for the presence of pathogens using *MALT* [30, 31]. A custom *RefSeq* genomic dataset containing bacteria, viruses, eukaryotes, and the human reference sequence GRCh38 was used to construct the *MALT* database using default parameters [32], with an index step size of 6. For each Erfurt sample, we screened all merged, de-duplicated sequences, applying a minimum complexity filter with a threshold of 0.3. We ran *MALT* (version 0.3.8), using the parameters --mode BlastN, --alignmentType SemiGlobal, --minPercentIdentity 0.85, --topPercent 1, --minSupport 1, -maxAlignmentsPerQuery 100. Results were then screened using the *HOPS* workflow [33] to determine whether there was evidence of authentic DNA from a set of 348 pathogens of interest among the Erfurt samples (Table S4). We assessed authenticity using a standard three step screening pipeline that considers (1) the edit distance distribution of all sequences that align to the pathogen of interest; (2) the presence of C-to-T (or G-to-A) sequence damage (which is characteristic of authentic ancient DNA); and (3) the edit distance distribution of the subset of aligned sequences that contain C-to-T damage [33].

Only a single pathogen, *Enterobius vermicularis,* passed all three authenticity screening steps (Table S4). However, *E. vermicularis*, commonly known as a pinworm, is a human intestinal parasite that has previously been sampled from ancient latrines [34]. It is therefore unlikely that this pathogen would have been present in the teeth of the Erfurt individuals at their time of death, and the most likely source of this DNA is contaminated groundwater in the Erfurt cemetery after burial. The groundwater contamination hypothesis is further supported by the presence of reads aligning to *E. vermicularis* in as many as 23/33 Erfurt individuals (Table S4), and by the fact that most cases (21/23; Table S4) failed tests for the authenticity of the ancient DNA. Similarly, five Erfurt individuals showed weak evidence for the *Schistosoma mansoni* pathogen, another water-borne human intestinal parasite that could have been introduced in the sampled teeth via environmental contamination after death. Very weak evidence was detected for pathogens associated with periodontal disease (*Parvimonas micra*, *Fusobacterium*, and *Fusobacterium nucleatum*) in a single individual, I14850 (the mother of family A, who was also buried in opposite orientation to all other individuals and possibly with her clothes on; Methods 1.1). Fewer than 30 reads aligned to each of these pathogens, and no evidence of C-to-T damage was detected in any of the aligned reads, however, these are all common oral pathogens [35], therefore it is possible that the reads represent authentic DNA that was present in the oral microbiome during individual I14850’s lifetime, but that was not well preserved in this sample.

Overall, our pathogen screening analysis did not find convincing evidence of any pathogens of interest among the Erfurt individuals. Particularly, we found no evidence of *Yersinia pestis*, the pathogen responsible for the plague, among any of the Erfurt individuals. The lack of evidence cannot completely rule out the possibility of *Y. pestis* infection in any given individual, as the preservation rate of *Y. pestis* DNA in teeth from individuals who are known to have died from plague has previously been estimated at only 37% [36]. However, the failure to detect any evidence of this pathogen among any of the Erfurt individuals suggests that the Erfurt cemetery is unlikely to have been a mass burial site for victims of a plague epidemic.

### 3. Qualitative ancestry analyses

#### 3.1. Principal components analysis (PCA)

We used *smartPCA* [37], with the option “lsqproject”, which enables projection of samples with high missingness when the PCs were learned from modern samples. The following populations from the Human Origins (HO) dataset were used : Armenian, Iranian, Turkish, Albanian, Bergamo, Bulgarian, Cypriot, Greek, Italian_South, Maltese, Sicilian, Italian_North, English, French, Icelandic, Norwegian, Orcadian, Scottish, BedouinA, BedouinB, Jordanian, Palestinian, Saudi, Syrian, Abkhasian, Adygei, Balkar, Chechen, Georgian, Kumyk, Lezgin, Ossetian, Jew_Ashkenazi, Jew_Georgian, Jew_Iranian, Jew_Iraqi, Jew_Moroccan, Jew_Tunisian, Jew_Libyan, Jew_Turkish, Jew_Yemenite, Basque, Spanish, Spanish_North, Druze, Lebanese, Belarusian, Croatian, Czech, Estonian, Hungarian, Lithuanian, Ukrainian, Canary_Islander, Sardinian, Finnish, Mordovian, Russian, and Polish.

Erfurt samples were projected on the PC space learned by these populations. Eight EAJ individuals with <50k SNPs were excluded from all PC analyses.

We also ran PCA with a larger sample size of MAJ, as follows. We merged the HO dataset with whole-genomes of *n* = 544 modern AJ [38]. These genomes were generated in Phase 2 of the Ashkenazi Genome Consortium (TAGC) sequencing project, and relatives were removed [39]. The merged dataset had about 470k SNPs. We used the same HO populations as above to learn the PC space, except that here we removed the HO MAJ population. Unlike the above analysis, here we projected both EAJ and the TAGC MAJ on the PC space. Due to the large MAJ sample size, we plotted the positions of these samples as a 2-dimenional kernel-density plot (Figure S9) using the function *stat_density_2d()* from the package *ggplot2* in *R*. The density was scaled to 1 using the argument *contour_var = “ndensity”*. To evaluate the effect of coverage on placement in PC space, we down-sampled a randomly selected subset of *n* = 525 MAJ individuals to match the set of SNPs covered by the EAJ samples. Each of the 25 (non-low-coverage) EAJ samples was used to match 525/25=21 MAJ individuals. We then selected a single allele at random from each down-sampled MAJ individual and repeated the projection of the MAJ and EAJ samples on the same PC space.

For the analyses of MAJ of Eastern European vs Western European origin, we merged the EAJ genomes with those from Behar et al. (2013) [40]. The merged dataset included 245,792 autosomal SNPs. The following modern populations from the Behar et al. dataset were used to learn the PCs: Abkhasian, Adygei, Algerian_Jewish, Armenian, Balkar, Bedouin, Belarusian, Bulgarian, Chechen, Croat, Cypriot, Druze, Estonian, French, French_Basque, Georgian, Georgian_Jewish, Greek, Hungarian, Iranian, Iranian_Jewish, Iraqi_Jewish, Italian, Jordanian, Kumyk, Lebanese, Lezgin, Libyan_Jewish, Lithuanian, Mordovian, Moroccan, Moroccan_Jewish, North_Ossetian, Orcadian, Palestinian, Russian, Saudi, Sephardi_Jewish, Spanish, Syrian, Tunisian_Jewish, Turkish, Polish, and Ukranian. The total resulting sample size was *n* = 882. We projected the Erfurt and MAJ (Ashkenazi_Jewish_Eastern and Ashkenazi_Jewish_Western) samples on the resulting PC space. Due to the smaller number of SNPs after merging the datasets, three additional EAJ samples had less than 50k SNPs and were excluded from the PC analysis.

#### 3.2. ADMIXTURE analysis

We ran *ADMIXTURE* version 1.3.0 [41] using the default parameters and using only SNPs that were covered in at least 18 Erfurt genomes (about 86k SNPs). We used the populations that were included in the PCA and the following populations: Erfurt, Egyptian, Han, Hazara, Kalash, Mbuti, Mandenka, Yoruba, Pima, China_Lahu, She, Adygei, Oromo, Somali, Dinka, Mala, Saami_WGA, Burbur_WGA, Ain_Touta_WGA, Azeri_WGA, Shaigi_WGA, Kurd_WGA, Assyrian_WGA, Naro, Shua, Nogai, Altaian, Dolgan, Tajik, Turkmen, Luo, Savo, Tunisian, Jew_Ethiopian, Algerian, Mansi, Jew_Cochin, Turkish_Balikesir, Saharawi, Irish, Moroccan, German, and Yemeni.

Similarly to the PCA, individuals with less than 50k SNPs were not included in the *ADMIXTURE* analysis.

### 4. f_4_ statistics, *qpWave*, and *qpAdm*

For the f_4_ and *qpWave* analyses, we merged the Human Origins dataset with whole-genomes of *n* = 544 modern AJ [38] (See also Methods 3.1). We used *AdmixTools* [42] version 5.1 for running the analyses. To avoid bias due to ancient DNA damage, we used only transversions SNPs in *qpWave* and *qpAdm* analyses (about 110k SNPs). In all population-level analyses, we (1) omitted one of each pair of first-degree relatives, keeping the individual with the higher coverage; and (2) included the low-coverage individuals (<50k SNPs), except in analyses that required the Erfurt-EU and Erfurt-ME group affiliation.

#### 4.1. f_4_ statistics

When inspecting our f_4_ statistics, we noticed unexpected results for f_4_ tests of the form f_4_(ME1, EU; ME2, Outgroup), where ME1 and ME2 are two Middle Eastern populations, EU is a European population, and the outgroup is chimpanzee. Naively, we expect ME2 to share more alleles with ME1, and thereby the statistic to be positive. In contrast, the statistic often came out negative. In fact, even when we split MAJ into two random groups and ran f_4_ tests of the form f_4_(MAJ1, ancient Germans; MAJ2, Chimp), the Z-score was −0.57.

A speculative explanation may be African gene flow into Middle Eastern populations [43, 44]. Denote by *α* the proportion of African ancestry in two Middle Eastern populations (assumed, without loss of generality, to be equal due to an admixture event that has pre-dated their split). A schematic admixture graph is shown in Methods Figure 1.

**Methods Figure 1.**
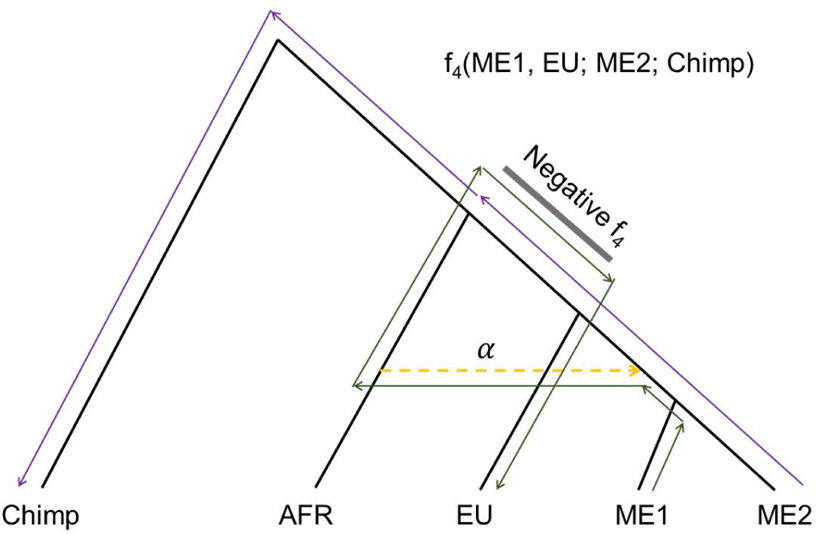
A schematic admixture graph for two Middle Eastern populations (ME1 and ME2), a European population (EU), an African population (AFR), and chimpanzee as an outgroup. The underlying tree is shown in black lines. A dashed orange line shows gene flow from Africans to the ancestors of Middle Easterners that has replaced a proportion *α* of these ancestors. The statistic f_4_(ME1, EU; ME2, Chimp) can be seen as a signed product of drift parameters (mean squared allele frequency differences) along branches shared between paths connecting ME1→EU and ME2→Chimp [42]. Given the admixture event ancestral to Middle Easterners, EU can be reached from ME1 in two paths, one of them involving admixture from Africans (green line). Similarly, Chimp can be reached from ME2 in two paths, one of them *not* involving the African admixture event (purple line). The gray bar labeled as “Negative f_4_” highlights a branch where the two paths proceed in opposite directions, hence contributing a negative term to the total f_4_ statistic. All other three possible pairs of ME1→EU and ME2→Chimp paths contribute non-negative terms to the statistic (see Eqs. (1), (2), and (4)). However, the total f_4_ statistic may be negative. For more details, see the “outgroup case” in ref. [42].

Next, write uME1 and uME2 to represent the genetic ancestry of the two Middle Eastern populations with their African ancestry excluded, and aME1 and aME2 to represent the African ancestry in these populations, such that an allele in ME1 has probability *α* to descend from aME1 and probability (1 −*α*) to descend from uME1 (and similarly for ME2). The statistic f_4_(ME1, EU; ME2, Outgroup) can be decomposed into a sum of four terms [42] (see also Methods Figure 1),

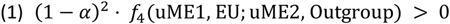

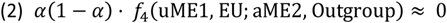

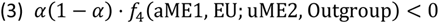

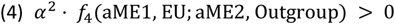

It can be seen that only the third term (Eq. (3)) is negative. (A graphical representation of this term is demonstrated in Methods Figure 1.) If this term is very large in absolute value, it may lead an overall negative f_4_ statistic. We note that this scenario is analogous to the “outgroup case” in the context of f_3_ statistics [42].

Given the problem with the interpretation of the numerical values of this type of f_4_ statistics, in further analyses we only considered the relative order of the z-scores from tests using this statistic.

#### 4.2. qpWave

We ran *qpWave* with the option “allsnps:YES”, which means that in each f_4_ test, the program uses all SNPs that were non-missing in all four populations, but a different set of SNPs can be analyzed for each underlying f_4_-statistic. We ran *qpWave* tests separately against reference (“right”) European and reference Middle Eastern populations. We selected the right populations as follows. In our *qpWave* analyses with European reference populations, we chose populations that represent main European ancestries: (modern) Russian, Norwegian, French, Spanish, Bulgarian, and Italian_North, with Primate_Chimp as an outgroup (first right population). Middle Eastern populations are closely related, and we therefore chose the right populations as follows. We ran f_4_ tests of the form f_4_(EAJ, MAJ; ME1, ME2) where ME1 and ME2 represent all possible pairs of populations from: BedouinA, BedouinB, Palestinian, Lebanese, Syrian, Jordanian, Egyptian, Saudi, and Druze. We used in *qpWave* all populations that were involved in tests with Z-score > 1.64: Druze, Lebanese, Jordanian, and BedouinA. We again used Primate_Chimp as an outgroup.

In tests with South-Italians we used samples of Sicilian and Italy_South together as one group.

#### 4.3. qpAdm

Here too, we used the option “allsnps:YES”. The reference populations (right populations) for the *qpAdm* analyses were: Mbuti, Ami, Basque, Biaka, Bougainville, Chukchi, Eskimo_Naukan, Han, Iranian, Ju_hoan_North, Karitiana, Papuan, Sardinian, She, Ulchi, and Yoruba. Mbuti was used as the outgroup (provided to *AdmixTools* as the first in the list of reference populations) in all analyses. In robustness tests, we replaced Mbuti with Ami as the outgroup. As in the *qpWave* analyses, in models with South-Italians we used samples of Sicilian and Italy_South together as one group. In models with ancient Germans, we used samples from [45], not including individuals with elongated skulls or with Southern European ancestry. The ancient Levant (Canaanite) samples included samples from [46] of Bronze-Age Megiddo (Megiddo_MLBA) and the ancient Rome samples included samples from [47] of Late Antiquity (Italy_LA.SG) and Imperial Rome (Italy_Imperial.SG).

For the analyses at the individual level, we used all SNPs, as the coverage of many individuals was already low. To guarantee that using all SNPs did not bias the results, we repeated the analyses at the population level with all SNPs instead of just transversions, and verified that the results remained qualitatively unchanged (Figure S16A). We included first-degree relatives in the individual-level analysis, but omitted the low-coverage individuals (<50k SNPs). For individuals for which the Eastern-EU ancestry proportion was inferred to be negative (Figure 2), we re-ran *qpAdm* with only Southern-EU and Middle Eastern sources.

To evaluate the potential contribution of East-Asians to the ancestry of EAJ, we tested models where the sources were Lebanese, South-Italians or North-Italians, Russians, and Han Chinese (Han were dropped from the reference populations for this analysis). The models had P-values of 1.9·10^−10^ and 1.8·10^−6^ with South- and North-Italians, respectively. When the target was Erfurt-EU, the P-values were 7.5·10^−8^ and 1.8·10^−4^, respectively. Given that the same models for EAJ without Han had plausible P-values (Table S7), we conclude that there is no detectable East-Asian ancestry in EAJ.

To quantify the difference in the Eastern European ancestry between MAJ and Erfurt-ME, we used *qpAdm* to model MAJ as the target of admixture between Erfurt-ME and Russians. We used only transversion SNPs. The model was plausible with P=0.76, with ancestry proportions 87% for Erfurt-ME and 13% for Russians. The model was plausible also with Germans as a source instead of Russians (P=0.74; ancestry proportions 86% for Erfurt-ME and 14% for Germans).

To quantify the relation between Erfurt-ME and Sephardi Jews, we used *qpAdm* to model Erfurt-ME using Turkish Jews and Germans as sources. We again used only transversion SNPs. The model was plausible with P=0.96, with ancestry proportions 97% for Turkish Jews and 3% for Germans. A model with Russians instead of Germans was also plausible (P=0.96; ancestry proportions 96% for Turkish Jews and 4% for Russians).

### 5. Estimating admixture times and the degree of endogamy

#### 5.1. DATES

We attempted to estimate admixture times using *DATES* [48, 49]. As *DATES* cannot infer the dates of multiple admixture events, we focused on the more recent event that likely involved Eastern Europeans. We used Erfurt-EU as the target admixed population, as most of Erfurt-ME individuals lack Eastern European ancestry. We omitted one of each pair of first-degree relatives, keeping the individual with the higher coverage. The source populations were chosen according to the plausible *qpAdm* models, with Russians as one source and Middle Easterners and Southern Europeans as the other source (Lebanese, Syrian, Jordanian, BedouinB, South-Italians, North-Italians, and Sicilian). Since *DATES* can estimate the time of admixture only between two populations, we used equal sample sizes (37 genomes each) from the Middle Eastern and the Southern European sources. The other source was Russians, with 71 genomes. We used the following *DATES* parameters: binsize: 0.001; maxdis: 1; qbin: 10; and lovalfit: 0.45. The estimated admixture time was 22.6 ± 8.1 (Figure S17A).

#### 5.2. Simulations

We used simulations to evaluate the accuracy of *DATES*. The simulated demographic history included two admixture events: the first between Middle Eastern and Southern European sources (35% and 65% ancestry from each source, respectively), and the second with Eastern Europeans (replacing 15% of the gene pool). We simulated two scenarios: one with the admixture events occurring 60 and 10 generations prior to sampling the target genomes, and another with events 70 and 20 generations prior to sampling. We generated the simulated genomes as follows. Each of the three sources included several populations, as listed in Methods Table 1. We phased the source genomes using the Sanger Imputation Service (https://www.sanger.ac.uk/tool/sanger-imputation-service) with the Haplotype Reference Consortium reference panel. To simulate the first admixture event, we randomly selected ten individuals from Southern European source and ten individuals from Middle Eastern source (these individuals were removed in the subsequent *DATES* analyses). We simulated the genomes as mosaics of haplotypes along the 22 autosomal chromosomes. We randomly assigned the source of each segment based on the simulated admixture proportions. We drew the length (in cM) of each segment at random from an exponential distribution with rate *G*/100, where *G* is the time of the admixture event in generations prior to sampling. We generated diploid genomes by pairing two simulated haploid genomes. We simulated ten genomes using this approach. We then used the simulated genomes and ten individuals from the East-EU source (who were removed from the subsequent *DATES* analyses) to simulate the second admixture event in a similar way. We simulated nine genomes, and down-sampled them to form pseudo-haploid data with coverage matching that of Erfurt-EU — the target group in the real *DATES* analysis.

**Methods Table 1.** A list of the populations we used as sources in the admixture analyses.

|  |  |
| --- | --- |
| Middle Eastern populations | Palestinians, Lebanese, Jordanians, Syrians, Egyptians, Bedouin A, Bedouin B, Saudis, Druze |
| Southern European populations | North-Italians, South-Italians, Sicilians, Greek |
| Eastern European populations | Belarusians, Lithuanians, Ukrainians, Russians |

We repeated each simulated scenario 50 times and analyzed the simulated genomes with *DATES*. We used a combined, balanced Middle Eastern and Southern European source, as in the real data analysis. Each of the sources included the populations listed in Methods Table 1. We found that the *DATES* estimates had an upward bias and a very large variance (Figure S17B). Hence, we conclude that *DATES* cannot reliably infer the admixture time between Middle Eastern/Southern European and Eastern European sources.

#### 5.3. The maximal level of gene flow since Erfurt

The *qpWave* test comparing EAJ and MAJ gave P=0.15 (Table S5), consistent with these groups being a clade with respect to reference European populations. To estimate the maximal degree of post-Erfurt gene flow into AJ that would still be consistent with these results, we used simulations. Specifically, we simulated AJ groups that have experienced increasing magnitudes of admixture with Eastern European sources. We then tested, using *qpWave* (with respect to the same European populations, as described in Methods 4.2), which simulated group is still inferred as a clade with modern AJ. The Eastern European sources included 30 samples from the populations listed in Methods Table 1. These samples were removed from the subsequent *qpWave* analyses. The simulated “admixed” AJ included 30 MAJ samples that were generated in Phase 1 of the Ashkenazi Genome Consortium sequencing project, and were not included in the MAJ dataset that was used for the original *qpWave* analyses (see Methods 4.2). For each admixture scenario, we simulated 30 samples, using the same procedure as described in the previous section, but without down-sampling. We used admixture times of *G* = 5, 10, 15, 20 generations. The *qpWave* P-values reported in Table S6 show that the maximal proportion of the AJ gene pool that could have been replaced by East-EU admixture and still remain consistent with being a clade with MAJ is about 2-4%. In our simulations, all gene flow was assumed to occur over a single generation. With continuous gene flow, a replacement of a proportion *m* of the gene pool per generation over 20 generations would lead to a total replacement of 1 − (1 − *m*)^20^ of the total ancestry. Equating to 4% and solving for *m* gives *m* = 1 − (1 − 0.04)^1/20^ = 0.2%.

### 6. The number of EAJ sub-groups

#### 6.1. The gap statistic

The gap statistic method [50] identifies the number of clusters that best fit the data, given a clustering method and a range of possible number of clusters. We used the function fviz_nbclust() from the *factoextra* package in *R*. We used *K*-means to cluster the samples (“kmeans” option with nstart = 25) based on the first two PCs. We set the maximal number of clusters to *K* = 4 and the number of bootstrap samples to 500. The low-coverage samples were not included in this analysis.

As a control, we determined the number of clusters in modern AJ and in Moroccan Jews (from the Human Origins dataset), either separately or jointly, with the results as expected (Figure S18).

#### 6.2. A significance test for the difference between the clusters

The clustering generated by *K*-means with *K* = 2 corresponds to our Erfurt-EU and Erfurt-ME groups. We used the function test_cluster_approx() from the *clusterpval* package in *R* [51] to calculate the P-value for the difference in means between those two clusters. The number of importance samples (“ndraws”) was 10,000.

#### 6.3. Simulations

To study the genetic composition of EAJ from a population genetics perspective, we simulated demographic scenarios with or without substructure. We then tested the similarity between summary statistics of the real Erfurt data and either simulated scenario. In the first scenario, there was a single admixture event between Middle Eastern (50%), Southern European (35%), and Eastern European (15%) sources (based on the model of ref. [52]) that has happened five generations prior to sampling (Methods Figure 2).

**Methods Figure 2.**
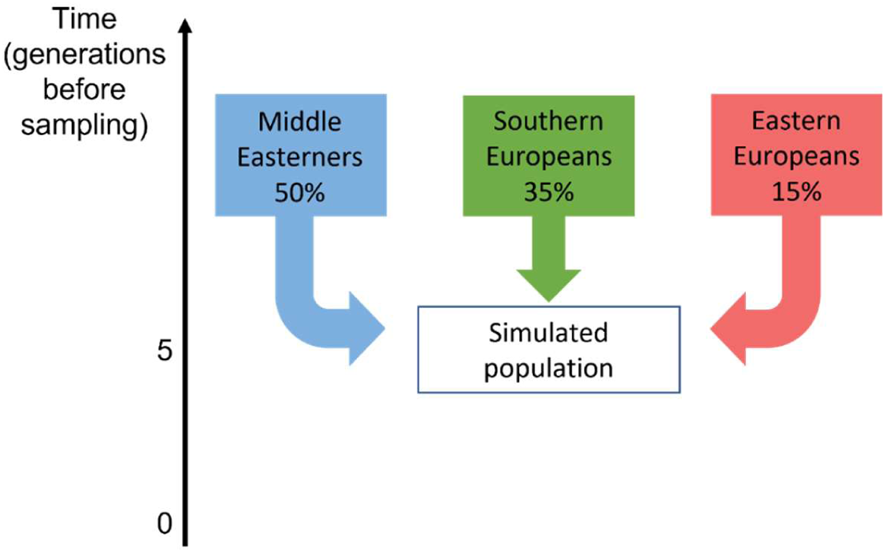
A schematic of an admixture model representing a single EAJ group. Under the model, the EAJ population has experienced a 3-way admixture five generations prior to sampling.

In the second scenario, we simulated two groups. Both groups experienced an admixture event ten generations prior to sampling between Middle Eastern (45%) and Southern European (55%) sources. One of the groups has experienced a second admixture event with Eastern Europeans (15%) five generations prior to sampling (Methods Figure 3).

**Methods Figure 3.**
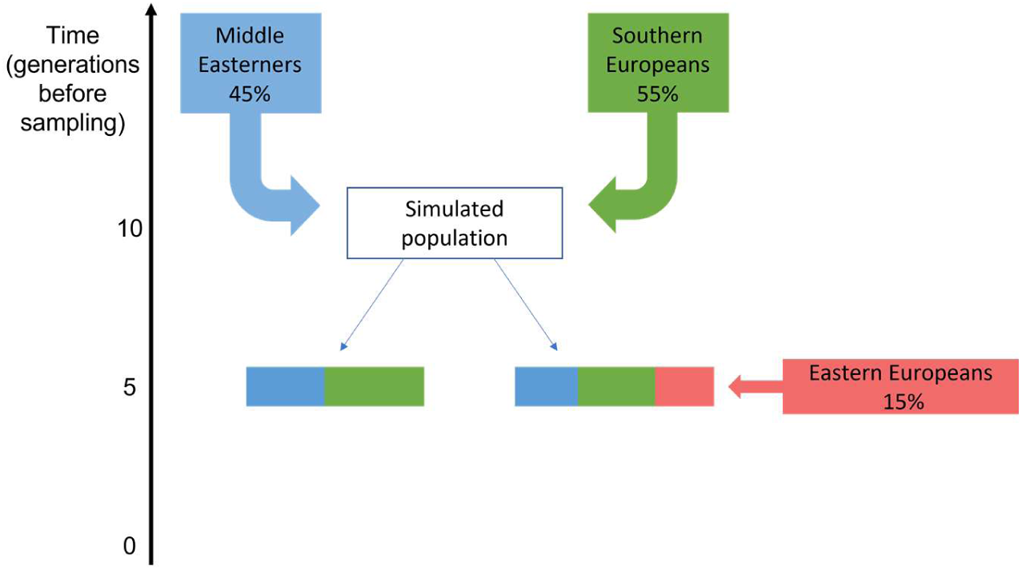
A schematic of an admixture model representing two EAJ groups. In this model, an ancestral population has experienced an admixture event between Southern Europeans and Middle Easterners ten generations prior to sampling. The population then split in two, and one group has experienced an additional admixture event with Eastern Europeans, five generations prior to sampling.

We generated the simulated genomes as described for the *DATES* analysis (Methods 5.2; without South-Italians and Sicilians in the Southern European source and without down-sampling). We simulated 30 genomes for the first demographic scenario (a single group), and 20 genomes for each group in the second (two-group) scenario.

We ran PCA and *qpAdm* analyses on each of the simulated datasets. When *qpAdm* inferred a negative East-EU ancestry proportion, we used the *qpAdm*-reported Middle Eastern and Southern European ancestry proportions had the East-EU ancestry proportion been set to zero. We used the Kolmogorov-Smirnov test (*ks.test* in *R*) to compare the PC1 distribution between the real data and the simulations (Figure S19). We used permutation testing to compare the proportion of individuals without Eastern European ancestry (as inferred by *qpAdm*; Figure S20) between the real and simulated data. In each permutation, we pooled the samples of Erfurt-EU (11 samples), Erfurt-ME (13 samples), and the simulation (30 or 40 samples). We then randomly labelled 24 samples as “Erfurt” and the remaining as “simulated”, and computed the difference in the proportion of individuals without East-EU ancestry between the two sets. The P-value was the fraction of permutations (out of 10k) in which the difference was greater than in the real data.

To validate that coverage does not affect the results, we ran PCA on pseudo-haploid down-sampled genomes from the two-group simulation. We matched the number of genomes and the number of SNPs of the (non-low-coverage) Erfurt-ME and Erfurt-EU samples. The results (Figure S19E) show no qualitative difference in the PCA plot compared to the full genomes.

#### 6.4. Mantel test

We used the Mantel test to investigate the correlation between group affiliation and the distances between the graves in the cemetery. We calculated the distances based on the approximate coordinates of the skeletons’ heads in the cemetery map (Figure S2). For the group affiliation, we set the distance between Erfurt-ME and Erfurt-EU to 1 and the distance within each group to zero. We used the function *mantel.rtest* from the *ade4* package in *R*. We also ran the Mantel test by replacing the group affiliation distance with the distances in the PC1-PC2 space (based on the PCA of Figure 1 of the main text and using Euclidean distances).

### 7. The mitochondrial DNA analysis

#### 7.1. Aligning the K1a1b1a sequences of the modern and ancient samples

To obtain sequence data from modern K1a1b1a carriers, we used *n* = 544 genomes of Phase 2 of the Ashkenazi Genome Consortium sequencing project (after removing related samples) [38]. We first discarded sites with GQ<40 as well as indels. Some sites had heterozygous genotypes (possibly due to heteroplasmy). We encoded these sites as having the alternate allele if the alternate allele was observed at more than 50% of the reads, and otherwise encoded them as having the reference allele. We used *bcftools consensus* [53] with parameters - H A using the rCRS reference sequence to generate an alignment of the sequences of all individuals. We then used *HaploGrep2* [25] to call mitochondrial haplogroups, and focused on the 107 carriers of K1a1b1a. For the ancient samples, haplogroups were previously called (Methods 2.2), and we considered only the 11 K1a1b1a carriers. Their median coverage was 304x (range: 47-447x; Table S1). We discarded indels and otherwise performed no additional filtering. We again used *bcftools consensus* to generate an alignment of the sequences, but this time with the RSRS reference sequence [21].

We noticed that all EAJ carriers had identical sequence except for a single site at position 16223. At that site, samples I13867, I13870, and I14903 had the C allele, while the remaining eight carriers had T. In the modern samples, with the exception of 16223, there were 36 segregating sites: 32 singletons, one doubleton, two variants that appeared in three samples, and one variant that appeared in four.

To determine whether MAJ carriers have significantly more diversity compared to EAJ carriers, we used down-sampling experiments. In each experiment, we sampled at random 11 MAJ carriers, and computed the number of pairwise differences (excluding site 16223). In comparison, the 11 EAJ carriers had no pairwise differences. Over 10,000 runs, the mean number of pairwise differences in MAJ carriers was 0.83 (SD: 0.44). The proportion of runs where MAJ carriers had zero pairwise differences (as in EAJ) was 1.83%. These results are expected given the longer time of the modern samples to their most recent common ancestor (TMRCA) compared to the ancient samples.

#### 7.2. The *BEAST* analysis

To estimate the TMRCA of the K1a1ba1 haplogroup, we used a Bayesian coalescent analysis, as implemented in *BEAST* 2 [54]. The input was an alignment of 11 EAJ carriers and 107 MAJ carriers. We set the dates of the EAJ samples to 650 years ago. For modeling mutations, we used the HKY model with Gamma distributed rates (four categories), and the strict clock model. For the population size prior, we used the coalescent Bayesian skyline with four segments. We also ran BEAST with a coalescent exponential population prior, but convergence was poor and we did not further consider this prior. For the skyline model, we ran 10 chains for 100 million steps each. For each chain, we used 10% of the steps as burn-in, and recorded the parameters every 100,000 steps, resulting in a total of 9000 samples. All other parameters were set to their default values. We combined the samples from all chains using *LogCombiner* and visualized the results using *Tracer* and *FigTree*. The total effective sample size (ESS) for the TMRCA was 3508. The mutation rate (across the entire mtDNA sequence) was estimated as 4.6·10^−8^ per bp per year, broadly in agreement with previous estimates [55, 56]. We show the posterior distribution of the TMRCA in Figure S21.

We present the maximum clade credibility tree based on these runs in Methods Figure 4. As expected given the pattern of polymorphism in 16223, the Erfurt lineages (IDs S1XXXXMT, where X is any digit) coalesce with the modern lineages based on their genotype at 16223, and the TMRCA of the Erfurt samples is the same as that of the modern samples.

**Methods Figure 4.**
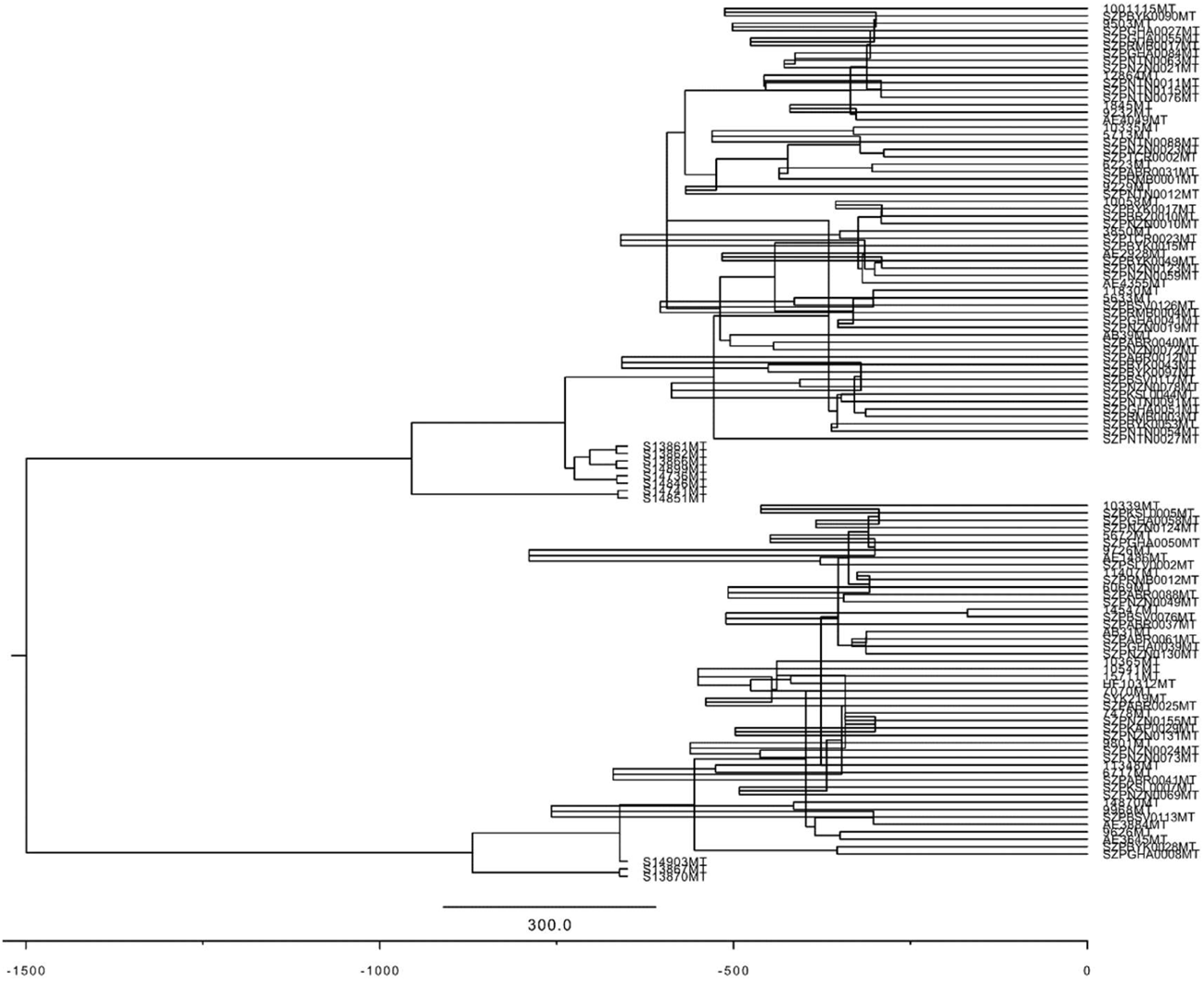
The maximum clade credibility tree of modern and ancient K1a1b1a carriers based on the output of *BEAST*. The tree was visualized using *FigTree*. The x-axis represents the time since the present. The ancient Erfurt genomes were assumed to be sampled 650 years ago.

We show the inferred past population size trajectory in Methods Figure 5. The inferred onset of population expansion is about 750 years ago, consistent with autosomal IBD results [57] (Figure 3A and Figure 3E). However, the large uncertainty associated with the inferred population size (see the 95% highest posterior density interval) does not permit definitive conclusions based on this data alone.

**Methods Figure 5.**
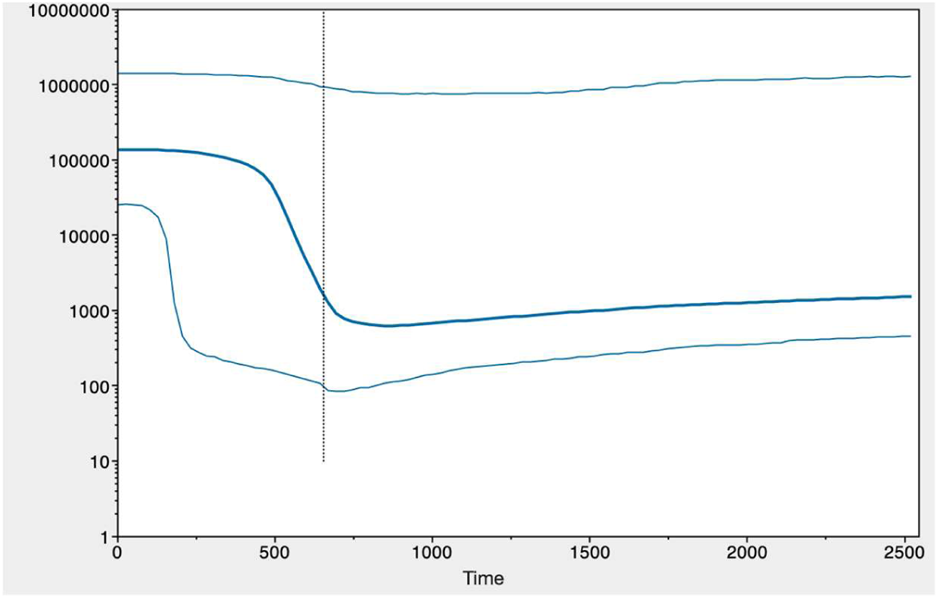
The reconstructed population size history based on the mtDNA sequences of modern and ancient K1a1b1a carriers. We ran *BEAST* on 11 ancient EAJ samples and 107 modern samples carrying the K1a1b1a mtDNA haplogroup, as explained in Methods 7.2 and in Figure S21. The plot shows a *Tracer* screenshot of the inferred population size history. The x-axis represents years before present. The y-axis is the effective population size. The thick middle line is the median estimate, and the two thin lines are the upper and lower bounds of the 95% highest posterior density (HPD) interval. Based on these estimates, the AJ population began to expand about 700-800 years ago. The dotted vertical line is the bottom of the 95% HPD interval for the TMRCA of K1a1b1a.

We finally performed the same *BEAST* analysis on the 107 modern carriers alone. Given that all samples are present-day, the mutation rate cannot be learned from the data itself. We used the value of the mutation rate as estimated in the joint modern-ancient analysis (4.6·10^−8^ per bp per year) to convert the estimated TMRCA to years ago. All other *BEAST* parameters were as in the joint analysis. The estimated median posterior was 1409 years ago, slightly earlier than in the joint analysis. The 95% highest posterior density (HPD) interval was 478-4041 years ago. Therefore, the availability of ancient samples from 650 years ago pushed the estimated TMRCA backwards (P<2.2·10^−16^; two-tailed Wilcoxon test comparing the two posterior distributions). This is expected, given the presence of the polymorphism 16223C/T in the Erfurt carriers, which excludes the possibility that the TMRCA of all carriers has lived prior to their time.

### 8. The founder event

#### 8.1. Detecting runs of homozygosity in the ancient genomes

To identify runs of homozygosity (ROH) within the ancient dataset, we used the Python package *hapROH* version 0.1a8 [58], using 5,008 global haplotypes from the 1000 Genomes Project as the reference panel and the pseudo-haploid genotypes of the ancient genomes as input. As recommended for datasets with genotype data for 1240k SNPs [58], we applied our method to 16 Erfurt individuals covered in at least 400k SNPs and called ROHs longer than 4 cM. We used the default parameters and post-processing of *hapROH*, which are optimized for this ancient DNA data type (described in detail in [58]). We report the sum of the lengths of all ROHs longer than 4cM in each individual in Table S2.

We manually inspected the ROH results by examining the positions of putatively heterozygous sites (Figure S22). Given that the called genotypes are pseudo-haploid (representing the allele of one randomly selected read), information on heterozygosity requires the original sequencing reads. For each SNP, we obtained the read counts for each allele from the processed BAM files using *samtools mpileup* [53]. Due to the low coverage, high quality diploid genotype calls could not be made. We identified putatively heterozygous sites as SNPs with at least one read supporting each of the reference and alternate alleles. While not all heterozygous sites could be detected using this approach (due to the low coverage), their depletion at inferred ROH segments is evident (Figure S22).

We verified that the number of ROH segments did not depend on the coverage. The correlation between the number of segments and the coverage was r=-0.07 (P=0.8). There was also no correlation between the coverage and the number of ROH segments of length <10cM, which are more prone to error (r=-0.01; P=0.96).

To contextualize the ROH results, we repeated the ROH analysis in other ancient populations. We used the Allen Ancient DNA resource (V50.0, Oct 10 2021; https://reich.hms.harvard.edu) and extracted previously published genomes from individuals who lived in the past ≈2000 years. We downloaded the data in *eigenstrat* format. As above, we used *hapROH* with the recommended settings [59] and applied it to pseudo-haploid genotypes. The populations we studied included Hungary Langobard [60]; Germany Early Medieval [45]; Italy Imperial, Late Antiquity (LA), and Medieval [47]; and Denmark Viking [61]. The sum of lengths of ROH segments in these populations is shown in Figure S23.

#### 8.2. Detecting IBD and ROH segments in the modern genomes

We considered sequencing data of *n* = 637 MAJ (from the two phases of the Ashkenazi Genome Consortium project [38, 57], after relatives and duplicates were removed [39]). To detect IBD sharing in MAJ, we used *IBDseq* [62] with default parameters.

We detected runs of homozygosity segments in *n* = 574 genomes (phase 2 data) using the software *bcftools/ROH* [63]. To meaningfully compare ROH in ancient and modern samples, we first down-sampled the modern data to the 1240k SNPs used in the ancient DNA analysis. We used the same sex-averaged genetic map as in the analysis of the ancient data (provided with the *hapROH* reference panel), and set the hidden Markov model transition probabilities to parameters optimized for 1240k data (see [58]; toA = 6.7e-8, toHW=5e-9). To obtain allele frequencies, we used the diploid data of the 574 modern individuals (using the VCF field MLEAF). As in the ancient data, we merged gaps of length up to 0.5 cM between two ROHs (both at least 2 cM long, one at least 4 cM). We manually inspected the ROH results by determining whether inferred ROH segments are depleted of heterozygous sites (Figure S22), considering only bi-allelic SNPs in the 1240k set. We finally removed nine individuals with total ROH length >50 cM (in segments longer than 4 cM), as these individuals likely have closely related parents.

#### 8.3. Inferring the parameters of a single-population model using IBD sharing

Our single-population demographic model is illustrated in Figure 3A. Under the model, the effective population size has been *N_a_* (diploids) until *T_b_* generations ago, at which point it became *N_b_* for *d* generations (the bottleneck). The population size then expanded exponentially, until reaching a present-day population size of *N*_e_. We assume generations are discrete. To infer these five parameters based on IBD sharing data, we used the counts of IBD segments across 11 length bins, equally spaced on a logarithmic scale between 4 to 15 cM. We then searched for the parameters of the demographic model that provided the best fit to the data.

To compute the expected number of segments in each length bin, we used theory from Ringbauer et al. (2017) [59] (see also [64–66]). Consider first two present-day chromosomes of length *L* (Morgan), and fix the coalescence time (i.e., their time to the most recent common ancestor (TMRCA)) to *t* generations before present. The expected number of IBD segments between these two chromosomes with length in the interval [*ℓ*_1_, *ℓ*_2_] is [59]

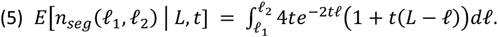

Denote the historical (diploid) size of the population size as *N*(*t*), for *t* = 0,1,2., …. Under our demographic model (Figure 3A),

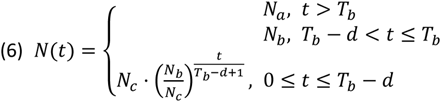

The probability of the TMRCA at a random locus to equal *t* is

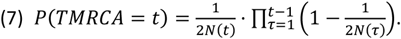

[Eq. (6) is true for any single-population demographic model. It is the probability not to coalesce until and including generation *t* − 1, multiplied by the probability of coalescence (1/2*N*) at generation *t*.] In the regime *t* > *T_b_*, *N*(*t*) = *N_a_* is independent of *t*, and thus the distribution of the TMRCA is

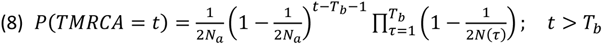

Summing over all *t*, the mean number of IBD segments of length in [*ℓ*_1_, *ℓ*_2_] between two chromosomes of length *L* is

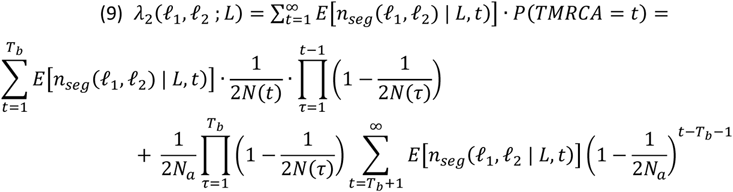

Finally, the mean number of (autosomal) IBD segments of length in [*ℓ*_1_, *ℓ*_2_] between *n* diploid genomes is

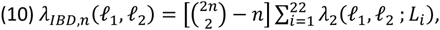

where *L*_i_ is the length of chromosome *i* = 1, …,22 in Morgan. The pre-factor 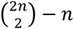 is the number of haplotype pairs when comparing *n* diploid individuals to each other. We used *Mathematica* to find a closed-form solution to the term with the infinite sum in Eq. (9). Eq. (10) thus provides the expected number of segments in each length bin under our demographic model.

Following previous studies [59, 64–66], we assumed that the observed number of segments in each bin is Poisson distributed with the expected mean (Eq. (10)) and independent across bins. This allowed us to write a composite likelihood for the observed segment counts given a proposed demographic model. Denote by *ℬ* the set of bins and by *c*(*ℓ*_1_, *ℓ*_2_) the observed number of IBD segments in the bin [*ℓ*_1_, *ℓ*_2_] (across *n* = 637 modern genomes). The composite likelihood is

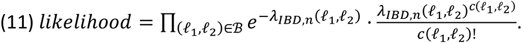

The log-likelihood is (up to an additive constant)

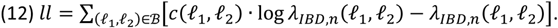

We maximized the log-likelihood with respect to the five model parameters (*N_a_*, *T_b_*, *N_b_*, *d*, *N*_e_) using the function *Deoptim* from the *R* package “DEoptim”. When inferring these parameters, we used the following boundaries to the search space: *N_a_* ∈ [1000, 50,000], *N_b_* ∈ [100, 5000], *T_b_* ∈ [20,60], *d* ∈ [1,30], and *N*_e_ ∈ [105, 107]. We ran the optimizer for 5000 steps after setting the seed to 1, and validated that the inferred parameters remained very similar when starting from other seed values. The inferred parameters of the model are listed in zs’f11 S10, model (A). We also inferred the model parameters after fixing the bottleneck duration to *d* = 1. The estimated parameters are listed in Table S10, model (B).

To compute confidence intervals for the inferred model parameters, we used parametric bootstrapping. In a naïve implementation of the non-parametric bootstrap, we would resample individuals with replacement. However, individuals are not independent with respect to the IBD segments (as segments are shared between pairs), violating the bootstrap assumptions. Additionally, detecting IBD sharing between an individual and itself would be nonsensical. We therefore generated each new bootstrap sample as follows. For each segment length bin, we drew a new count for the total number of segments as a Poisson variable with mean equals to the count in the real data. We generated 100 bootstrap samples, and, for each sample, we inferred all model parameters as for the real data. For each parameter *θ*, we computed the 95% confidence interval for the parameter as 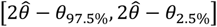 [67], where 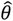 is the estimate based on the real data, and *θ*_2.5%_ and *θ* _7.5%_ are the 2.5- and 97.5-percentiles, respectively, of the estimates across the bootstrap samples. We calculated the 2.5-percentile as the average of the estimates that ranked second and third (out of 100), and similarly for the 97.5-percentile.

#### 8.4. Inference using modern ROH segments

We next attempted to infer the parameters of the single-population model (Figure 3A) using counts of ROH segments in modern genomes. The derivation is exactly as in Eqs. (5) to (12) above, except that in Eq. (10), the pre-factor 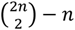 is replaced by *n* (the number of haplotype pairs that would generate ROH is *n*),

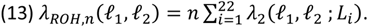

We also assumed *N*(*t*) → ∞ for *t* = 1,2. This represents the fact that two chromosomes in the same individual cannot coalesce in the immediately following generation, as well as that sib-mating is unlikely. We then found the demographic parameters that maximized the composite likelihood as with the IBD data. Across runs, optimization converged to two distinct optima of similar likelihood, likely due to the small amount of data. The first is listed in Table S10, model (C) (*N_b_* = 1295, *T_b_* = 30, and *d* = 11). The other optimum was at *N_b_* = 598, *T_b_* = 25, and *d* = 3. Both models date the end of the bottleneck to around the same time and have similar bottleneck intensities, but they differ in their bottleneck duration.

#### 8.5. Modeling consanguinity in the ancient individuals

The empirical results (Figure 3D) suggest that the inferred demographic model (based on IBD sharing in modern genomes; Table S10, model (A)) underestimates the expected number of ROH segments in the ancient genomes. We hypothesized that this may be a result of consanguinity in the ancient individuals, which could have generated additional ROH segments. We therefore attempted to fit a model where the demographic parameters are as in Table S10, model (A), but a proportion *α* of the ancient individuals are offspring of first cousins.

To determine the expected number of ROH segments of a given length under the consanguinity model, we followed Ringbauer et al. (2021) [58]. For children of *r*^th^ full-cousins, the expected number of ROH segments due to consanguinity in a chromosome of length *L* Morgan is (see also Eq. (5))

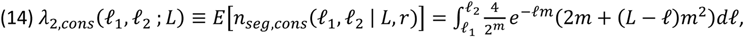

where *m* = 2*r* + 4 is the total number of meioses between the two chromosomes of the child and the most recent common ancestor. For the case of first cousins, where this common ancestor is a great-grandparent, *m* = 6.

The mean number of ROH segments of length in [*ℓ*_1_, *ℓ*_2_] due to consanguinity in *n* genomes of children of first cousins is

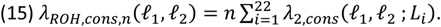

The mean number of ROH segments between two ancient chromosomes of length *L* due to genetic drift, i.e., due to coalescence under the demographic model, is

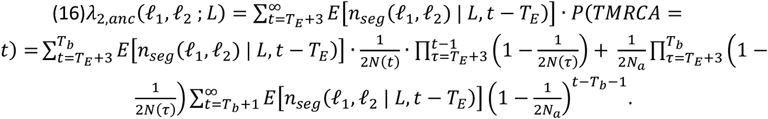

*T*_E_ is the number of generations ago when the Erfurt population has lived. We assumed a generation interval of 25 years, slightly lower than previous studies [68–72], given that early AJ often married extremely young [73]. Given our radiocarbon dating to the 14^th^ century, i.e., about 650 years ago, this gives *T*_E_ = 26. *N*(*t*) is given by Eq. (6). Eq. (16) is the same as Eq. (9), except that no coalescence is possible until *T*_E_ generations ago and that the number of generations to the TMRCA is *t* − *T*_E_. We started the sums at *T*_E_ + 3 to represent the constraint of no sib-mating. The mean number of ROH segments in *n* ancient genomes due to drift is

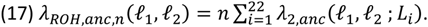

Finally, the total number of ROH segments of length in the interval [*ℓ*_1_, *ℓ*_2_] in the ancient genomes has mean

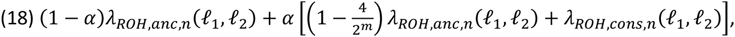

where *λ_ROH,ane,n_* is the expected number of ROH segments due to genetic drift (Eq. (17)), and *λ_ROH_*_,*cons*,*n*_ is the expected number of segments due to consanguinity (Eq. (15)). The term 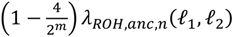 represents ROH in children of first-cousins in genomic regions where the two chromosomes do not coalesce at the shared great-grandparents. We then assumed, as above, that the observed total number of (ancient) ROH segments (across the *n* = 16 ancient genomes) in each length bin follows a Poisson distribution with the given mean. This gave a composite-likelihood similar to Eq. (11).

We then fixed all demographic parameters to their inferred values as in Table S10, model (A) and used the optimization procedure to find the value of *α* that maximized the log-likelihood. Note that we used neither modern IBD nor modern ROH data. For the set of bins *ℬ*, we used (here and in all other models based on ancient ROH) 29 bins equally separated on a logarithmic scale between 4 to 40 cM. This is different from modern data, in that we also considered relatively long ROH segments. The long segments likely appeared because (*i*) the parents of some individuals may have been related, and (*ii*) the individuals lived closer in time to the bottleneck.

The inferred proportion of individuals who were children of first cousins (Table S10, model (D)) was *α* = 0.22, which corresponds to 3-4 individuals out of the total of 16. This estimate is reasonable given the distribution of total ROH lengths (Figure 3C). However, the fit to the ROH counts did not sufficiently improve (Figure 3D), possibly as consanguinity generates predominantly very long segments (mean about 17cM for children of first cousins), whereas the ROH counts were underestimated at shorter lengths. We therefore no longer considered consanguinity in our next models.

#### 8.6. Modeling a narrower or a longer bottleneck

We next hypothesized that the excess of ROH segments in EAJ is due to the EAJ population experiencing a narrower or a longer bottleneck compared to what we inferred based on IBD sharing in MAJ (Table S10, model (A)). In the following, we fixed some of the parameters of the modern-based model (Table S10, model (A)) and inferred the other parameters using ancient ROH data to fit models with a narrower or a longer bottleneck.

For a model with a narrower bottleneck, we fixed the bottleneck starting time to *T_b_* = 41 and inferred the ancestral population size (*N_a_*) and the bottleneck size (*N_b_*) based on the counts of ROH segments in the ancient genomes. We assumed that the population size remained at *N_b_* until the time of the EAJ individuals. In other words, the EAJ population size history has been

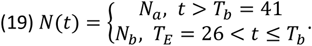

We plugged this expression for *N*(*t*) into Eq. (16) and used Eq. (17) to compute *λ_ROH,ane,n_*(*ℓ*_1_, *ℓ*_2_), the expected number of ROH segments in the ancient genomes under our demographic model (without consanguinity). We again assumed a Poisson distribution for the number of segments in each bin, and used the optimization procedure to find the values of *N_a_* and *N_b_* that maximized the composite-likelihood. [While our focus was on the bottleneck size *N_b_*, and we generally did not attempt the interpret the (highly uncertain) estimate of *N_a_*, we found numerically that allowing *N_a_* to vary improved the fit.]

This above described procedure did not yet use any modern data. Accordingly, we did not infer the values of *d* and *N*_e_, as these do not appear in Eq. (19) and thus do not affect ancient ROH levels. Once we estimated *N_a_* and *N_b_* using the ancient ROH data, we fixed these values (along with *T_b_*, which is fixed to its value from Table S10, model (A)) and estimated *d* and *N*_e_ using modern IBD data, in the same way we inferred the full model (assuming *T_b_* − *d* ≤ *T*_E_). The complete set of inferred model parameters is given in Table S10, model (E). The fit of the model to the ancient ROH data is shown in Figure 3D.

We used a similar procedure to infer the parameters of a model with a longer bottleneck. First, we fixed *N_a_* and *N_b_* to their values from Table S10, model (A), giving the following EAJ population size history,

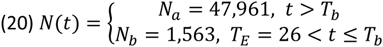

We then used the ancient ROH data to infer the value of *T_b_* (assuming 80 ≥ *T_b_* ≥ 41) by comparing the observed segment count to the expectation based on Eq. (17), as above. We finally used the modern IBD data to infer the parameters *d* and *N*_e_. The inferred parameters are listed in Table S10, model (F), and the fit is shown in Figure 3D. Both a narrower and a longer bottleneck fit the ancient ROH data (in particular the narrower model). However, neither model fit the modern IBD data (Figure S25A).

We note that the more severe bottleneck we inferred in EAJ is not necessarily in contradiction with our observations that EAJ have more diverse ancestry than MAJ. A joint interpretation of these two trends would imply that EAJ had fewer founders compared to MAJ, but that these founders were more diverse in terms of their EU/ME genetic ancestry proportions.

#### 8.7. Joint inference based on modern and ancient data

To identify model parameters that would fit both modern and ancient data, we used the same five-parameter model (Figure 3A) with population size history *N*(*t*) given in Eq. (6), and attempted to infer its parameters using ancient and modern data jointly. Recall that the log-likelihood for the modern IBD data was

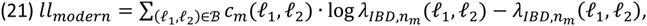

where *c_m_*(*ℓ*_1_, *ℓ*_2_) is the number of IBD segments of length in [*ℓ*_1_, *ℓ*_2_] shared between any pair of haplotypes among *n_m_* = 637 modern genomes, and *λ_IBD_*_,*n*_*_m_*(*ℓ*_1_, *ℓ*_2_) is the expectation based on Eq. (10). Similarly, for the ancient data,

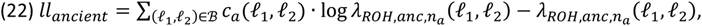

where *c_a_*(*ℓ*_1_, *ℓ*_2_) is the number of ROH segments of length in [*ℓ*_1_, *ℓ*_2_] in any of the *n_a_* = 16 ancient genomes, and *λ_ROH,ane,n_*(*ℓ*_1_, *ℓ*_2_) is the expectation based on Eq. (17). We defined a joint log-likelihood as

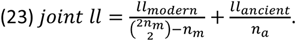

This definition addresses the issue that the number of haplotype pairs is about 50k times larger in the modern IBD data compared to the ancient ROH data. Under Eq. (23), each log-likelihood class (modern IBD/ancient ROH) contributes roughly equally to the log-likelihood. For both IBD and ROH, *ℬ* was 29 bins between [4, 40]cM. We then searched for the maximum likelihood parameters exactly as before, except that we enforced the time of EAJ sampling (*T*_E_ = 26) to be within the bottleneck (i.e., *T_b_* ≥ *T_E_* ≥ *T_b_* − *d*). The inferred parameters are listed in Table S10, model (G). The fit to modern data was still imperfect (Figure S25B).

#### 8.8. Inferring the parameters of a two-population model

To reconcile the demographic models of EAJ and MAJ, we expanded the model to account for substructure in AJ during the Middle Ages. While an expanded model can take various forms, we sought to minimize overfitting, and hence added only a single parameter. In our model, the AJ population split *T_b_* generations ago into two groups. The first represents EAJ, with effective population size *N_b_*. The second had population size *N_a_* − *N_b_*. (This is an arbitrary choice, in order to model different population sizes for the two groups but without further increasing the number of parameters.) The populations then merged *d* generations later, with proportions *f* and 1 − *f*, respectively, and then expanded exponentially until reaching the present population size. Note that we did not explicitly model the substructure within EAJ, again in order not to add parameters, and given that we have modeled substructure in AJ as a whole. The model is illustrated in Figure 3E.

We defined a joint modern-ancient likelihood as in Eqs. (21)-(23). The likelihood based on the ancient ROH data remains the same, as the model is identical to that of Figure 3A from the perspective of the EAJ population. As above, we assumed that the bottleneck must have spanned the time of EAJ, i.e., *T_b_* > *T_E_* > *T_b_* − *d*. For the likelihood based on the modern IBD data, we modified Eq. (9) (for the mean number of segments between a pair of chromosomes of length *L*) as follows,

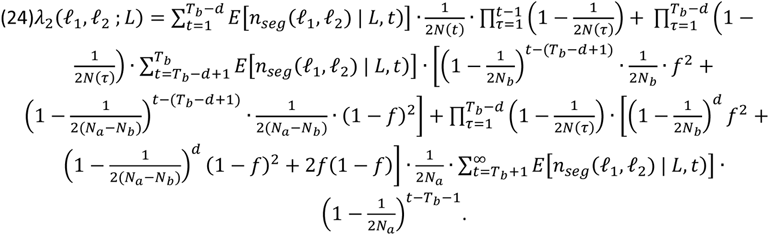

The population size history is

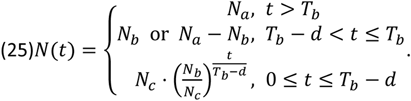

In Eq. (24), for coalescence to occur within the first sub-population, both lineages must descend from that population, which happens with probability *f*^2^, and similarly for the second sub-population (probability (1 − *f*)^2^). For coalescence to happen in the ancestral (pre-split) population, coalescence must not have happened during the bottleneck. This is the case if both lineages descended from the first population (probability *f*^2^) and then there was no coalescence (probability 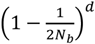), if both lineages descended from the second population followed by no coalescence (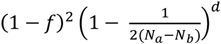) or if each lineage descended from a different population (probability 2*f*(1 − *f*)). We used the same optimization procedure as in the other cases to obtain the maximum likelihood estimate for the six parameters (*N_a_*, *N_b_*, *T_b_*, *d*, *f*, and *N*_e_). To compute confidence intervals, we used parametric bootstrap as for the single population model. Here, we re-sampled segment counts per bin for both modern IBD and ancient ROH from Poisson variables with means as in the real data.

#### 8.9. Model selection

The improved fit of the IBD and ROH data to the two-population model could be due to its increased complexity. To evaluate whether the fit is sufficiently improved to justify the additional parameter, we used parametric bootstrap [74], testing the null hypothesis that the real data comes from the single-population model. We simulated segment length counts under the (five parameter) single-population model. For each simulated dataset, we fit both the single-population and the (six-parameter) two-population model, and we recorded the increase in composite log-likelihood when (over)fitting the more complex model. We then determined whether the increase in likelihood observed in the real data is beyond what is expected when the data is truly derived from the single-population model.

To simulate from the single-population model, we used the best-fit parameters we inferred jointly from the MAJ and EAJ data (Table S10, model (G)). We calculated the expected number of segments in each length bin (29 bins from 4 to 40 cM) for both IBD segments in MAJ (Eq. (10)) and ROH segments in EAJ (Eq. (17)). We then drew a new count at each length bin as a Poisson variable with mean equals to the expected count. For each simulated dataset, we maximized the log-composite-likelihood based on Eq. (23) for either the single-population or the two-population model. Over 100 simulated datasets, the difference in the optimal log-likelihood between the two models, *ll*_two *populations*_ − *ll_single population_*, was in the range [−0.003, 0.09]. For the real data, the log-likelihood difference (based on the models in Table S10, models (G) and (H)) was 0.21. We thus conclude that, with P<0.01, we can reject the hypothesis that the real data derives from the single-population model.

#### 8.10. Simulations of the two-population model

We finally sought to validate, using simulations, that we can use data of the type available to us to accurately infer the two-population model parameters. We used *ARGON* version 1.0 [75] to simulate the demographic model shown in Methods Figure 6 (all population sizes are haploids).

**Methods Figure 6.**
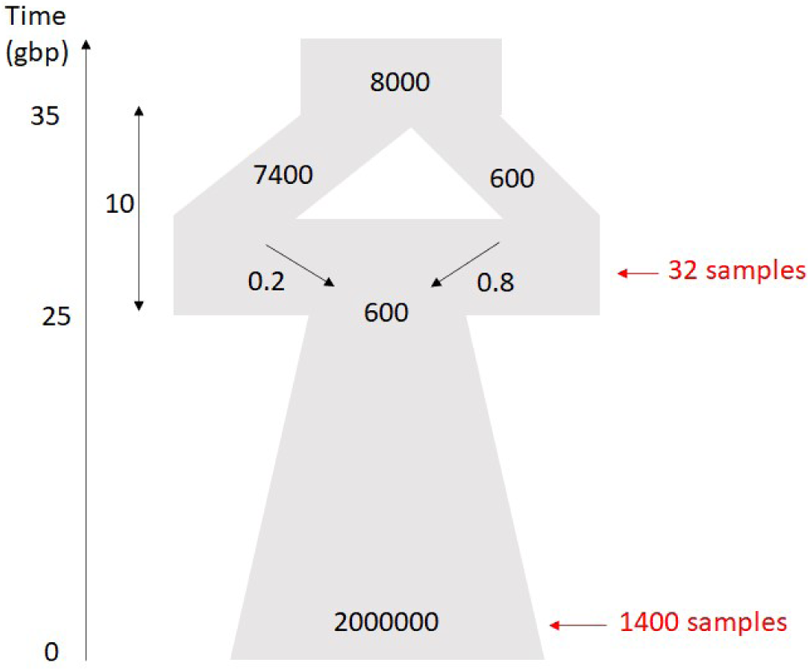
The figure illustrates the demographic model we simulated to test the accuracy of our inference method. All times are in generations before present (gbp). The width of the diagram at different time points is (schematically) proportional to the effective population sizes. The indicated population sizes are in haploid individuals. In our simulations, we sampled either 1400 haploid chromosomes at present, or 32 haploid chromosomes 26 generations before present (red arrows), representing our modern and ancient samples, respectively.

To mimic the extreme imbalance in the real data between the number of modern and ancient observations, we sampled 1400 haploid individuals from the present-day population (“modern” data), and then ran the simulation again and sampled 32 haploid individuals from the right population at the end of the bottleneck (“ancient” data). These sample sizes roughly correspond to our 637 modern genomes and 16 ancient genomes. For each individual we simulated a single chromosome of length 280 Mb with the default recombination rate of 1cM/Mb. The simulator provided ground-truth information on all IBD segments shared between all pairs of either “modern” or “ancient” individuals. [We computed pairwise IBD also for the “ancient” genomes (and not runs of homozygosity), in order to generate sufficient amount of data given that we simulated only a single chromosome.] We then recorded the number of segments per length bin (29 bins between [4–40] cM).

We first used the simulated modern data alone to infer the demographic parameters of the single-population model (Figure 3A). We used the same methods as for the real data. The inferred parameters were *N_a_* = 46.1 ⋅ 10, *N_b_* = 780, *T_b_* = 35, *d* = 12, and *N*_e_ = 1.8 ⋅ 10^6^ (population sizes are in haploid individuals). As we observed for the real data (Figures 3B and 3D), the fit was good for the modern IBD data, but it underestimated the number of ancient ROH segments (Methods Figure 7A). We then used the simulated modern and ancient data jointly to infer all six parameters of the two-population model (Figure 3E). The inferred parameters were very close to their simulated values: *N_a_* = 9.1 ⋅ 10, *N_b_* = 704, *T_b_* = 36, *d* = 13, *f* = 0.84 and *N*_e_ = 3.4 ⋅ 10^6^. The fit was now good for both modern and ancient data (Methods Figure 7B). These results, while not comprehensive, hint that even given the relative scarcity of the ancient ROH data, our method should be able to accuracy infer the parameters of the two-population model.

**Methods Figure 7.**
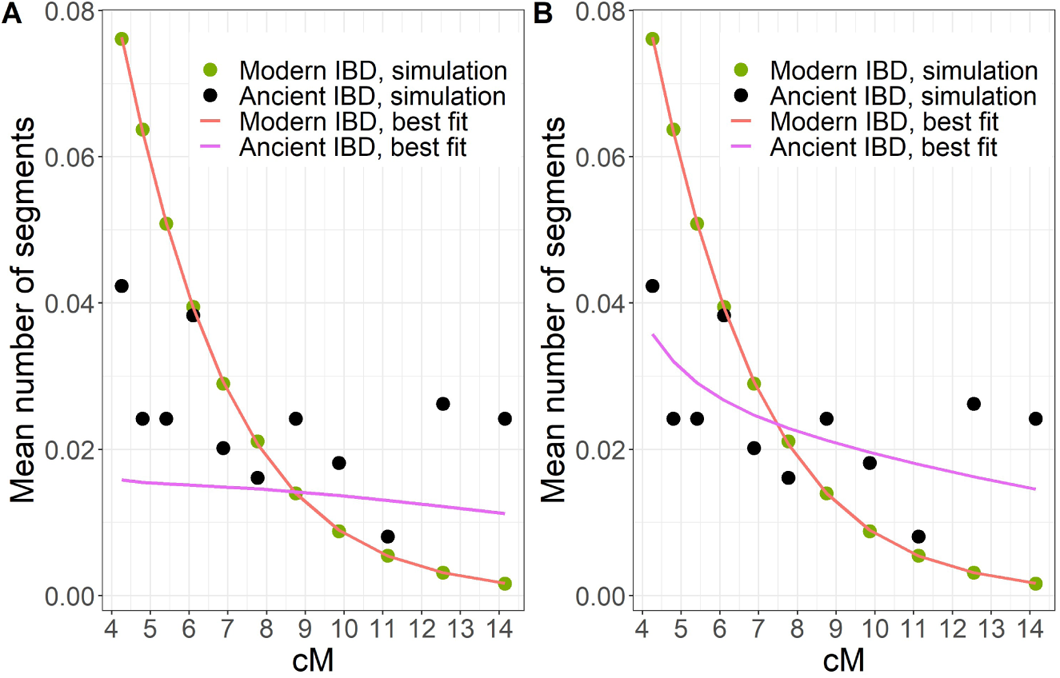
Simulated and fitted IBD segment counts. (A) The mean number of IBD segments per haplotype pair across segment length bins. We simulated those IBD segments based on the demographic model shown in Methods Figure 6. Symbols show simulated mean counts (legend). Lines (legend) show the best fit based on a single-population demographic model (Figure 3A). (B) The same simulated data as in (A), but with the fitted lines based on the two-population demographic model (Figure 3E).

#### 8.11. Presence of Ashkenazi founder alleles in Erfurt

Modern AJ carry dozens of founder pathogenic variants, as well as other alleles that are nearly absent in other populations. Detecting these founder alleles in EAJ would strengthen the case that EAJ have already experienced the Ashkenazi bottleneck. We defined founder alleles as those having minor allele frequency >0.5% in MAJ and <0.01% in non-Finnish Europeans (using *gnomAD* [76]). To exclude variants that may have a Middle Eastern source (which is not covered by *gnomAD*), we used 221 Middle Eastern individuals from the Human Origins dataset (including Palestinians, Saudis, Bedouin A, Bedouin B, Egyptians, Druze, Lebanese, Syrians, and Jordanians), and excluded alleles that appeared more than once among these individuals. Finally, we removed SNPs that were genotyped in less than three EAJ samples (after removing first-degree relatives), leaving a total of 216 SNPs.

Among the EAJ individuals, we excluded all children from families A and B, as well as one individual who was not genotyped in any of the founder SNPs, and was thus uninformative. Within the remaining 29 EAJ individuals, we detected 15 founder alleles in 11 individuals (Table S2). All variants except one appeared in just a single individual. The remaining variant appeared in three individuals, and thus the total number of copies of these alleles was 17.

#### 8.12. Binomial simulations of founder allele counts

To determine whether the number of founder alleles present in EAJ is as expected if EAJ has already experienced the AJ bottleneck, we used binomial simulations. In each run and for each founder allele, we drew an EAJ allele count as a binomial variable with *n* equals to the number of EAJ individuals that were genotyped in that SNP, and *p* equals to the allele frequency in MAJ (based on *gnomAD*). Note that we used the number of EAJ individuals (and not twice the number) as our genotypes are pseudo-haploid. In each run, we recorded the number of alleles (out of 216) that appeared in at least one individual. The distribution of the number of observed alleles across 10,000 runs is shown in Figure S26. The [2.5,97.5]-percentiles of the number of observed alleles were [14, 32].

The number of founder alleles in EAJ may be underestimated due to a “reference allele bias”. To model the bias in our simulations, we assumed that if the real genotype is heterozygous, there is probability 0.55 that the observed allele will be the reference. [A homozygous genotype (alternate or reference) will be observed correctly.] Hence, the probability to observe the alternate allele changes from *p* to *p*^2^ + 0.45 · 2*p*(1 − *p*). When we repeated the simulations with these probabilities, the [2.5,97.5]-percentiles for the number of observed alleles became [12, 29].

The number of founder alleles in EAJ may also be underestimated due to the conditioning on exceeding a given frequency in MAJ. This is because alleles that increased in frequency since ancient times to exceed the cutoff in the modern population will be included, but alleles that decreased in frequency below the cutoff will not. Therefore, our binomial simulations would tend to overestimate the number of alleles that are expected to be present in EAJ. This problem should exacerbate with higher allele frequency cutoffs. Indeed, when we repeated the analysis with a cutoff of 1% (as opposed to 0.5% above), six alleles were observed in EAJ, which was at the lowest range of the expectation based on binomial simulations ([2.5,97.5]-percentiles: [6, 18]). In contrast, when we set the cutoff to 0.1%, 32 alleles were observed in EAJ, compared to simulated [2.5,97.5]-percentiles of [26, 49].

In conclusion across analyses, the number of founder alleles observed in EAJ was consistent with the expectation based on modern AJ allele frequencies.

#### 8.13. Founder alleles in Erfurt sub-groups

*Erfurt-EU and Erfurt-ME.* The proportion of individuals carrying founder alleles was similar between Erfurt-EU (4/9, 44%) and Erfurt-ME (6/13, 46%). However, this result may be confounded by the higher coverage in Erfurt-EU. We therefore used quasi-Poisson regression to model the number of founder alleles carried by an individual as a function of the group affiliation (Erfurt-EU/Erfurt-ME), with the number of covered founder SNPs as an offset. Mathematically,

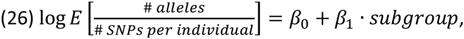

where the sub-group was coded as 1 for Erfurt-ME and 0 for Erfurt-EU. Even after adjusting for coverage, the correlation between the number of carried founder alleles and the group affiliation remained insignificant (Methods Table 2).

**Methods Table 2.** The quasi-Poisson regression for the number of founder alleles vs the Erfurt subgroup affiliation. The model is described in Eq. (26).

|  | Coefficient | Standard error | P-value |
| --- | --- | --- | --- |
| Intercept ($\beta_0$) | -5.36 | 0.52 | 1.62e-09 |
| Erfurt-ME ($\beta_1$) | 0.50 | 0.65 | 0.45 |

*K1a1b1a carriers.* We found that 8/11 (73%) carriers of K1a1b1a also carried at least one founder allele, compared to 3/18 (17%) of carriers of other mtDNA haplogroups (P=0.005, two-tailed Fisher’s exact test). Here too, we accounted for differences in coverage using quasi-Poisson regression with an offset,

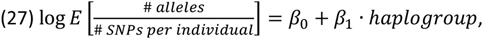

where the haplogroup was coded as 1 for K1a1b1a and 0 for all others. Here, the correlation diminished after accounting for coverage, though the P-value remained less than 0.05 (Methods Table 3).

**Methods Table 3.** The quasi-Poisson regression for the number of founder alleles vs the mtDNA haplogroup. The model is described in Eq. (27).

|  | Coefficient | Standard error | P-value |
| --- | --- | --- | --- |
| Intercept ($\beta_0$) | -5.7424 | 0.4863 | 3.58e-12 |
| K1a1b1a ($\beta_1$) | 1.2631 | 0.5788 | 0.038 |

### 9. Pathogenic founder variants

#### 9.1. Defining the pathogenic founder variants

We started with Supplementary Table 4 from our previous AJ sequencing paper [57]. We excluded 11 variants that were not present in modern AJ (based on *gnomAD*) or had higher frequency in other populations in *gnomAD*, leaving 62 variants. Among these, 47 were present in our reference panel and could thus be imputed. Incidentally, none of the pathogenic variants appeared in the list of founder alleles defined in the previous section, as most of the variants were not genotyped. Of the genotyped variants, four had frequency >0.01% in Europeans, one was not genotyped in the modern Middle Eastern samples, and one did not appear in *gnomAD*.

#### 9.2. Imputation using *PHCP*

To determine the genotypes of SNPs not on the array, we used imputation based on a reference panel of modern AJ whole genomes. Imputation of pseudo-haploid ancient DNA is not supported by current tools, and we therefore developed a method based on our previous *PHCP* model [46]. Briefly, *PHCP* (Pseudo-Haploid ChromoPainter) is an extension of the *ChromoPainter* model [77], which itself is based on the Li-Stephens [78] hidden Markov model (HMM). *PHCP* models a target ancient sequence as a mosaic of modern haplotypes (“donors”). For each SNP, the hidden state of the HMM is a pair of modern haplotypes that the target is “copying” from. The observed (haploid) ancient allele is assumed to derive from each of these two haplotypes with equal probability. Transitions between donor haplotypes along the target sequence are assumed to be due to ancestral recombinations, and emissions model recent mutations and genotyping and other errors that may lead to imperfect copying of the donor haplotypes. The full transition and emission probabilities were described in our earlier publication [46], where we used the population labels of the inferred donors of each target to learn about the target’s ancestry composition.

Here, we used the *PHCP* model for imputation of an ancient target genome. To obtain genotype probabilities, we used the forward-backward algorithm and computed, for each SNP of the target, the marginal posterior probability of each possible pair of donor haplotypes. For SNPs in the reference panel that were not covered in the target or were not on the array, we assigned to the target the marginal probabilities of the nearest covered SNP (in cM). For each SNP, we divided all haplotype pairs into three classes, based on the diploid genotype they imply for the target (AA/AB/BB). For each of the three possible genotypes, we defined their marginal probability as the sum of the marginal probabilities of all pairs of donors in that genotype’s class. In downstream analyses, we used for each SNP the most likely genotype.

To impute the EAJ genomes, we used *n* = 702 MAJ genomes (both phases of the Ashkenazi Genome Consortium sequencing project [38, 57]; without removing relatives or other samples). We considered only autosomal chromosomes. We set the parameters for *PHCP* as *N_e_* = 64.57, *θ* = 0.0014 [46]. To reduce the running time, we used, for each chromosome, a set of 200 donor haplotypes that were most informative for that chromosome. The donor haplotypes were ranked based on the number of SNPs where they share an alternate allele with the target [79] (and scaled by the number of SNPs where they have an alternate allele, as in [79], although we used all available sites and not just the rare variants). The *PHCP* imputation software is available at https://github.com/ShamamW/PHCPImpute.

#### 9.3. Testing the imputation accuracy

*Mendelian inconsistency.* We used the two families to estimate the rate of Mendelian inconsistency in the imputed genomes. The analysis included SNPs both genotyped and fully imputed, because even genotyped SNPs were imputed from haploid to diploid. Given that only one parent was available from each family, Mendelian inconsistency would be observable only when the parent and child carry opposing homozygous genotypes. Thus, in each family, we started with all SNPs imputed as homozygous in the parent, and counted the number of SNPs that were imputed in each child as homozygous to the opposite allele. Overall, we tested three parent-child pairs: mother (I14850) and son (I14853) and mother and daughter (I14898) from family A, and father (I14904) and daughter (I13869) from family B (Table S12). To compare the results to a baseline, we repeated the analysis for the mother from family A and the daughter from family B, and for the mother with an arbitrarily selected high-coverage sample (I13866) from Erfurt-EU (since the two families belong to Erfurt-EU).

*Concordance against masked genotyped variants.* We masked genotypes in the 216 founder SNPs defined above and in three pathogenic variants that were genotyped in our array and were present in at least one EAJ sample (F11/p.E135X, F11/p.F301L, and LRRK2/p.G2019S; Table 1). We then imputed these SNPs and tested the concordance between genotyped and imputed alleles. Among the 219 SNPs that were tested, 9 were not present at the reference panel and were not imputed.

There were overall 20 cases (across all individuals and SNPs) where the ancient (pseudo-haploid) genotype showed the alternate allele. Among these, we correctly imputed at least one alternate alleles in 15 cases. In the remaining five cases, the imputed genotype was homozygous reference, and we thus estimate the false negative rate as 5/20=25%. This is an upper bound, as some of these errors may be false positives in the ancient DNA genotypes. Interestingly, the false negative rate was 15% in Erfurt-ME (2/13) but 50% in Erfurt-EU (3/6), though the number of variants is too small to draw any conclusion (P=0.26; Fisher’s exact test).

We evaluated the false-positive rate as follows. First, we identified all cases, across the 29 individuals that were tested for founder SNPs (i.e., without the children of family A and B and without the individual who was not covered in any of the founder SNPs), and across all masked SNPs, where the pseudo-haploid genotype was the reference allele. We then computed the proportion of these cases where an alternate allele was imputed. (In all such cases, the imputed genotype was heterozygous.) The observed proportion was 13/2541=0.005. This is an upper bound on the false positive rate, because in some cases, the true genotype may have been heterozygous, but only the reference allele was observed. To quantify this, we computed the expected number of cases where the true genotype should have been heterozygous, based on MAJ allele frequency (*gnomAD*), but the observed allele is the reference. Specifically, we multiplied the MAJ frequency of each founder allele by the number of genomes that had the reference allele at this SNP and summed over all SNPs. The expected number of heterozygotes was 24.04, greater than the imputed number of alternate alleles (13). This could be due to (i) false negatives of imputation; or (ii) lower frequency of the founder alleles in EAJ compared to MAJ (Figure S26). Either way, our estimate of the imputation false positive rate (13/2541, or about 1/195) is likely as an upper bound.

*GLIMPSE.* We used *GLIMPSE* (v1.0.0) [80] with *n* = 702 MAJ individuals as the reference panel (as for *PHCP*). Diploid genotype calls were generated using *bcftools mpileup* (v1.10.2) [53]. We imputed all the autosomal biallelic SNPs and indels in MAJ. However, genotype likelihoods were used only for biallelic SNPs (generated by *mpileup*) as input to build the phasing and imputation model. Indels were imputed without genotype likelihood information due to their more severe reference bias.

To evaluate the concordance between *PHCP* and *GLIMPSE*, we compared their output on the pathogenic variants (see Methods 9.1 and Table S11). We first considered eight variants that were not genotyped and that were imputed by *PHCP* as having at least one alternate allele with posterior probability >97%. A carrier of one variant was not run in *GLIMPSE* due to low coverage. For the remaining variants, *GLIMPSE* imputed at least one alternate allele with probability >50% in 6/7 variants. We then considered six variants where *GLIMPSE* imputed an alternate allele with probability >97%. *PHCP* imputed the alternate allele with probability >50% for all such variants.

#### 9.4. High-confidence pathogenic founder variants present in EAJ

After imputing the EAJ genomes with *PCHP* and *GLIMPSE*, 16 pathogenic variants were either genotyped or successfully imputed (Table S11). We defined a set of high-confidence pathogenic variants present in EAJ based on the following criteria: (i) the *PHCP* probability for having at least one alternate allele was >97% and (ii) the alternate allele was detected by *GLIMPSE* with probability >50%. One variant (CLRN1, NP_001182723.1:p.N48K) was detected by *PHCP* with probability >97% in a sample that was not imputed with *GLIMPSE* due to its low coverage (Table S11); we considered this variant as high-confidence. We also defined a set of low-confidence pathogenic founder variants present in EAJ. The criteria defining these variants were: (i) *PHCP* marginal probability >97% and *GLIMPSE* probability <50%; or (ii) *PHCP* probability in the range 50%-97%; or (iii) *PHCP* marginal probability <50% and *GLIMPSE* probability >50% (Table S11).

For each high-confidence variant detected in EAJ, we determined whether the relevant gene was present in pre-conception carrier screening (PCS) panels. We considered four pre-conception sequencing-based panels aimed at the Ashkenazi-Jewish population: Genpath (https://www.genpathdiagnostics.com/hcp/womens-health/carrier-screening/ashkenazi-jewish-cancer-screening/), SEMA4 (https://sema4.com/products/test-catalog/ashkenazi-jewish-carrier-screen/), fulgent (https://www.fulgentgenetics.com/beacon-ashkenazi-jewish-female-carrier-screening) and Baylor genetics (https://geneaware.clinical.bcm.edu/GeneAware/default.aspx). For each variant, we indicated in Table 1 the number of panels in which the gene is included.

### 10. Phenotypes

#### 10.1. A polygenic score for height

We estimated the heights of 14 individuals based on bone measurements (see Methods 1.3; Table S14). For the polygenic score analysis, we excluded the child from family B, as her father was also included. We added to the height of each female 9.84 cm, which is the mean difference between males and females in our sample. For the polygenic score, we used summary statistics from Yengo et al. (2018) [81] without additional adjustments. Our array had 704,830 SNPs overlapping the summary statistics. We calculated the score for each individual using *Plink* version 1.9 [82] (--score) with the “sum” option and otherwise default settings, such that missing genotypes were imputed to their allele frequencies. The mean number of informative SNPs per individual was 340,799 (range 150,420-507,616).

#### 10.2. Other phenotypes

Details on the alleles for lactase persistence, eye color, hair color, and the putative plague risk alleles [83] appear in Table S11. We excluded the children from both families, leaving 30 individuals (60 chromosomes). All seven SNPs were present in our array. To compute allele frequencies in EAJ, we used the most likely genotype based on the *PHCP* imputation results. We computed 95% confidence intervals for the allele frequencies using Wilson’s method, as implemented in the *binconf* function from the *Hmisc* package in *R*. The allele frequencies in modern AJ were obtained from *gnomAD*. For the analysis of the plague putative risk alleles, we compared the EAJ allele frequency to that of MAJ in *gnomAD* using a one-tailed binomial test in the direction of the change observed in Immel et al [83].

## Supplemental Information

### 1. History of early Ashkenazi Jews and the Erfurt Jewish community

#### 1.1. The origins of early Ashkenazi Jews

There are currently two main competing historical theories to explain Ashkenazi Jewish early origins. The first theory holds that AJ are at least partially descendants of Roman-period Diaspora Jews. This theory is supported by dispersed historical and archaeological evidence along the Germanic frontiers of the late Roman Empire. On the basis of the results of the recent Cologne synagogue excavations — a building that the excavator controversially dates to the early Carolingian period — it was argued that there is direct demographic continuity between the scattered late Roman Jewish “proto-Ashkenazic” presence in the region and the Jewish communities of the Rhineland of later times [1, 84].

The second theory regards AJ as a purely medieval formation that did not arise until the 10^th^ century. According to this theory, AJ communities initially arose in the form of just a handful of family groupings in a few episcopal and royal urban centers, and that early AJ were the descendants of Jews from Southern Europe. There was continuous Jewish presence in Southern Europe since Roman times, and an extensive network of intercommunal ties linked these Jewish communities economically, culturally, and demographically to other Jewish communities around the Mediterranean [85–88]. Research suggests that early AJ of Northern Europe were the recipients of Jewish liturgical, legal, mystical, and linguistic practices from medieval Southern Italy.

The available historical evidence does not support a third hypothesis according to which early AJ were primarily descendants of early medieval non-Jewish converts to Judaism known as Khazars — a polyethnic tribal constellation then resident in the Caucasus and adjacent regions [88].

#### 1.2. The medieval Jewish community of Erfurt

The medieval Jewish community in Erfurt was the oldest in Thuringia, and existed between the late 11^th^ century to 1454. The Erfurt old synagogue is the oldest (partly) intact synagogue in Europe [89]. The community practiced Jewish rabbinical law [90]. Erfurt belonged to the territory of the archbishop of Mainz, but was surrounded by territories of different counts and nobles. The Jews in surrounding towns were also part of the Erfurt community, and their deceased were buried in Erfurt [91, 92]. In the second half of 13^th^ century, several families from the region of Franconia (in today’s Northern Bavaria) immigrated to Erfurt and probably to other towns in Thuringia. After 1300, about 30 families or more lived in Erfurt [92].

In 1349, a wave of pogroms (massacres) occurred, and many Jews in Erfurt and other towns in Thuringia were murdered [93–95]. Like in other cities with resident Jewish communities, in Erfurt too anti-Jewish persecutions started even before the arrival of the Black Death [95]. Some families, particularly the wealthy ones, survived in territories in the region where pogroms did not occur, and could even keep parts of their property. It is unknown whether these families lived in Erfurt or in nearby towns before 1349, but in 1354, they belonged to those who resettled in Erfurt [94].

After 1354, the newly founded, “second” community of Erfurt grew to become one of the largest communities in Germany [96]. As the lists of rentals show, about 50 Jewish families lived in Erfurt by the 1370s. The rapid increase in the population between the 1350s and the 1370s was due to migration of several Jewish families from Bohemia, Moravia, and Silesia to Erfurt and nearby towns (see SI 1.3.) [97]. As for the first community, surrounding Jewish settlements were part of the Erfurt community and buried their deceased in Erfurt [98].

Some families left Erfurt in the 1380s and 1390s, whereas after 1400, families from nearby towns moved into Erfurt. The number of Jews in Erfurt after 1407 is unknown, as no lists of rentals remained [91, 92]; but it is known that in 1418, at least 20 families lived in Jewish settlements in the region that were part of the Erfurt community [98]. During the 1430s and 1440s, Jews in some areas of Thuringia were expelled or were forced to leave, and a few moved to Erfurt [98]. In 1453, the city council of Erfurt no longer granted the protection of the Jews. The Jewish families left Erfurt within a year, marking the end the medieval Jewish community [99]. Resettlement of Jewish individuals only occurred in the 19^th^ century in a different part of the city.

#### 1.3. Documented migration from the East into the second Erfurt community

The information on the origin of Jewish families who migrated to Erfurt comes mainly from records of home rentals from 1354 to 1407. Most persons in these records are mentioned with surnames, which often name the town where they lived before [97]. Information in topographic surnames is limited, as surnames can change, and as the time period when a person has lived in the other town could vary. But in some cases, we have independent sources validating the former place of residence. From 1354, and especially in the 1360s, many families moved to Erfurt whose surnames refer to former places of residence in Bohemia, Moravia, and Silesia. For example, several families came from Breslau (Wrocław) after a pogrom in 1360, some after moving to Wrocław from other Silesian towns. After 1400, there are no known cases of families migrating into Erfurt from the East [97, 98].

Towns in Silesia (present-day Poland) from where families moved into Erfurt include Bunzlau/Bolesławiec (one family, first mentioned in the records in 1383), Liegnitz/Legnica (two related families in 1360), Löwenberg/Lwówek Śląski (one person whose family was originally from Brno), Breslau/Wrocław (one family in 1355/6, more families after 1360), Striegau/Strzegom (one family in 1366), Schweidnitz/Świdńica (one person in 1389), and Glatz/Kłodzko (one family in 1380). Towns in Bohemia and Moravia (present-day Czech Republic) from where families moved into Erfurt include the neighboring towns Braunau/Broumov and Náchod (two families in 1360 or later, who moved through Wrocław), Prag/Praha (one family in 1366), Pilsen/Plzeň (one family in 1365), Eger/Cheb (one family in 1359), and Brünn/Brno (one family in 1363, with a son-in-law in Vienna) [97]. We note that one man moved to Erfurt from Poland in 1327 (i.e., in the first community).

### 2. A supplementary discussion on early Ashkenazi Jewish history

#### 2.1. Interpretation of the inferred genetic ancestry

Our results provide new evidence for (although do not definitely prove) the theory of AJ origins in Italy (SI 1.1), given the good fit of *qpAdm* models that had Italy as a source, particularly Southern Italy. Southern Italy is one of the very few places in Europe where there is evidence for Jewish demographic and cultural continuity from the late Roman into the early Medieval period and beyond [87, 100–107]. During this timeframe, the Jewish communities of Southern Italy were at the crossroads of Jewish Mediterranean life. They were in direct contact with the Jewish communities of Byzantine and early Muslim Palestine from whom they received liturgical traditions that they transmitted into Europe and that later turned up in the AJ prayer book. They were also in touch with Jewish communities elsewhere in the Eastern Mediterranean by virtue of the fact that Southern Italy was part of the Byzantine Empire into the late 11^th^ century.

All the evidence currently available indicates that during the Roman and early Medieval periods Jews were highly integrated in Southern Italy. There is historical evidence that there was at least some gene flow between Jews and non-Jews in Southern Italy, because, in the late Roman and early Medieval periods, imperial and ecclesiastical authorities tried to prevent the practice of intermarriage between Jews and Christians, as well as the phenomenon of conversion of non-Jews to Judaism. When, in due course, highly accomplished and connected Jews from Southern Italy started moving north, they were joined by others from central and northern Italy. For example, the Kalonymus family—a Jewish family from Rome, but with roots in Southern Italy—is known to have had major impact on AJ intellectual life in 10^th^-century Mainz and Speyer [87, 108]. This was the multilayered migratory legacy that may be reflected in the Southern European genetic ancestry we observed in our models for the genomes of Erfurt Jews.

Our *qpAdm* models with a South-Italian source suggested that only a small proportion of EAJ ancestry derived from Middle Eastern populations. This may be interpreted to imply that present-day AJ derive only a small proportion of their ancestry from ancient Judaeans; and if so, most AJ ancestry would owe its origin to European converts. While this is one possible explanation, modern Italians themselves have had much higher proportions of ME admixture since at least European Imperial Roman times [47] and this is especially the case in modern Southern Italy [109]. Thus, an alternative explanation for these observations is that the true ME proportion in AJ is higher than in our fitting model, and that the actual contribution of Italians is not as large as suggested by this analysis. Under this scenario, good *qpAdm* fits are obtained when using Southern Italians as sources simply because Southern Italians are a modern population that harbors a relatively high proportion of ME ancestry with less impact from additional immigration waves that subsequently affected ME populations and may make modern ME populations relatively poor proxy sources for the ME ancestry in AJ. If this alternative explanation is right, the true ME proportion could be higher than in our fitting models, even higher than the ≈44% for the models using Northern Italians. At present, we believe both types of scenarios are plausible, along with scenarios that involve features of both. Co-analysis of ancient DNA data from the Middle East and the Italian peninsula from the periods of Antiquity and the early Medieval period would make it possible to distinguish them.

Our genetic data suggest that some Erfurt individuals had elevated levels of European ancestry, likely Eastern European-related. A possible explanation is the documented migration into the second Erfurt Jewish community from Bohemia, Moravia, and Silesia (SI 1.3). However, this requires that Jews living in these areas had previously admixed with local non-Jewish populations. Partly supporting this hypothesis may be the presence of names of Slavic origin among medieval Jewish women in Bohemia, particularly as it stands in contrast to naming practices common among Jews in medieval times [110, 111]. Finally, the genetic data suggested a high degree of endogamy in AJ through the last ≈700 years. Historical evidence indicates that the social practice of intermarriage between Jews and Christians was frowned upon by medieval Jewish and Christian authorities [112, 113]. Our genetic results suggest that in practice there was indeed very little gene flow into the Jewish community since this period.

#### 2.2. Timing demographic events in Ashkenazi history

Our modeling of shared haplotypes dated the onset of the AJ bottleneck to ≈40-45 generations ago, or approximately about 1000-1200 years ago. This period is well before the time in the late 12^th^ century when the persecution of Jews in the Rhineland became endemic. The appearance of a bottleneck in the early stages of the AJ community formation could reflect the historical evidence that the original AJ settlers comprised only a few dozen families, which were not always welcome and lacked the benefit of a fully developed Jewish community [114].

Our models dated the onset of expansion of AJ to about 20-25 generations ago, or approximately about 500-700 years ago. This confirms historical research pointing towards a gradual demographic growth within the Jewish community in German lands. The growth is hard to quantify numerically, but, especially from the 1300s onwards, it appears to have been substantial, considering the rapid increase in the number of towns that accommodated Jewish communities [115].

In this work, we were unable to reliably estimate the dates of the historical admixture events of AJ in Europe. Our previous work inferred a minor post-bottleneck gene flow event from Eastern Europeans based on a depletion of EU ancestry in IBD segments [52] (as such segments are expected to descend from ancestors who lived during the bottleneck). However, with a model of a prolonged bottleneck (about 20 generations; Table S10), such a depletion may be observed also if the admixture event had happened late during the bottleneck. Our previous work estimated that admixture between Middle Eastern and European sources in AJ history occurred about 30 generations ago [52]. This date may be associated with the admixture event with Eastern Europeans. Unfortunately, our EAJ genomes did not provide additional insight, as we found that a state-of-the-art tool for admixture time inference (*DATES*) provided unreliable results under simulations of AJ-like history (Figure S17B).

## Supplemental Figures

**Figure S1.**
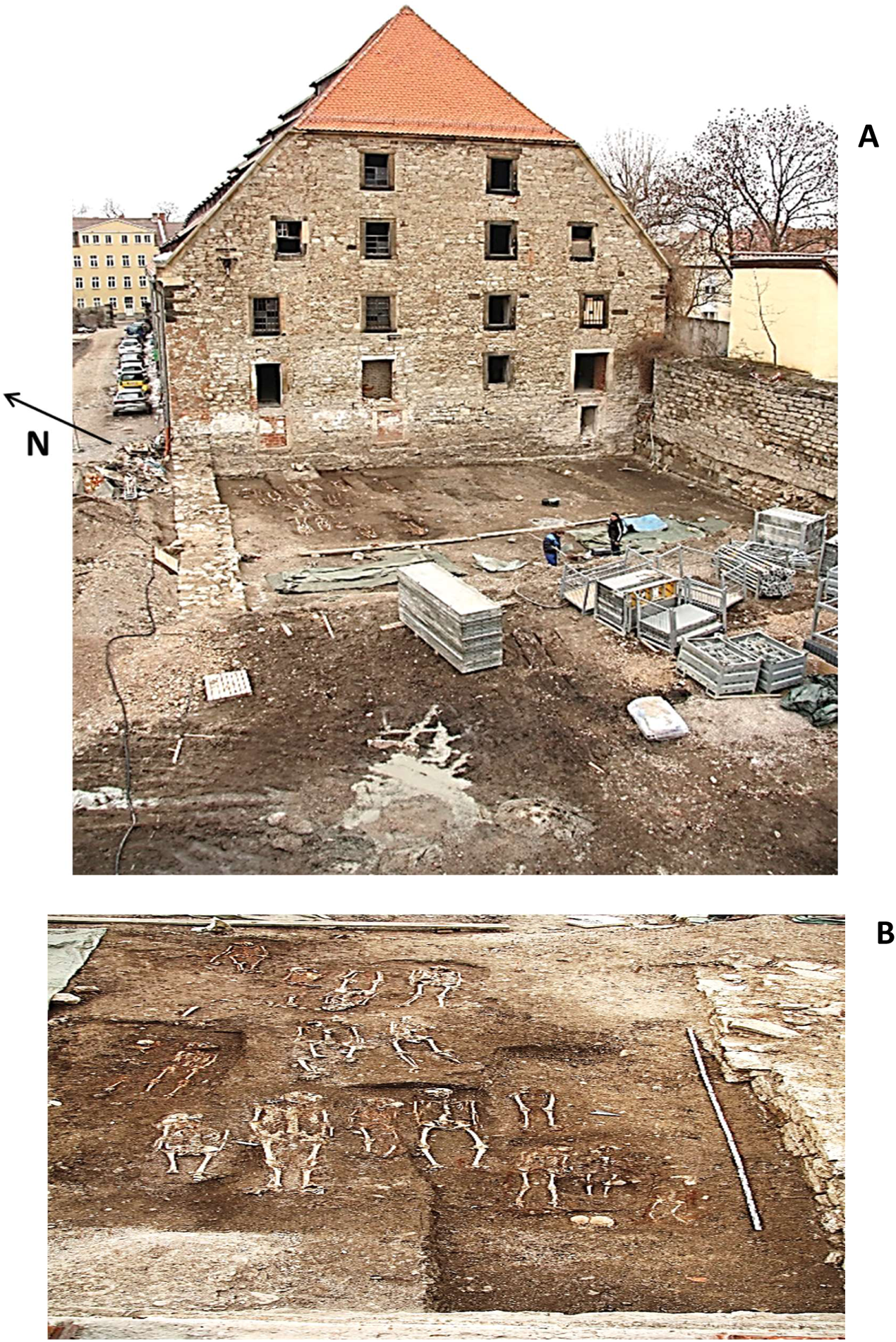
The archeological site. (A) The excavation of the medieval Jewish cemetery in Erfurt, Germany. The large structure behind (to the east of) the excavation site is a granary (the “Kornhofspeicher”) that was built in the 15^th^ century on top of the cemetery and now serves as a garage. Behind the granary is Moritzstraße (distant building to the left), which delimits the area of the original cemetery to the east. To the right of the site is the wall of the old town of Erfurt, which bounded the cemetery from the south. To the left (north) is an outer city wall that was built later. The area between the walls underwent salvage excavations before the construction of a ramp in 2013. The cemetery likely extended further north and west beyond the area of the excavation: the main part of the cemetery was possibly north of the city’s fortifications. (B) Skeletons that were discovered in the excavation (view from the granary).

**Figure S2.**
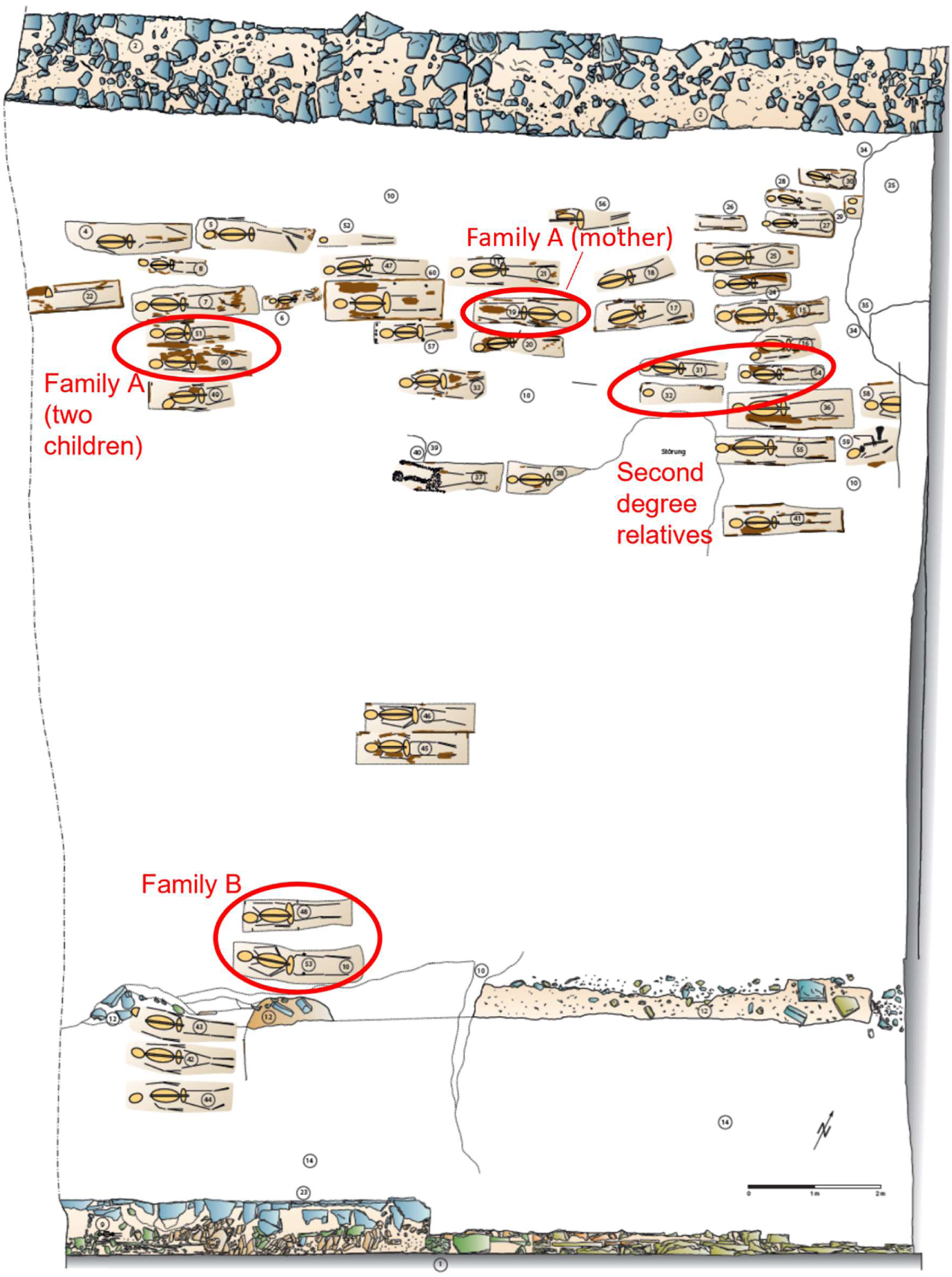
The cemetery’s map. A map of the medieval Jewish cemetery of Erfurt along with grave numbers. Family members are marked in red ellipses. The first city wall (shown on the right-hand side of Figure S1A) is marked as 1 at the bottom of the map. The outer city wall, only preserved below the surface, is marked as 2 at the top of the map. Number 12 on the map is probably the filling of an older moat belonging to the first city wall, numbers 10 and 14 are layers of earth, and 34/35 is a recent ditch (20^th^ century).

**Figure S3.**
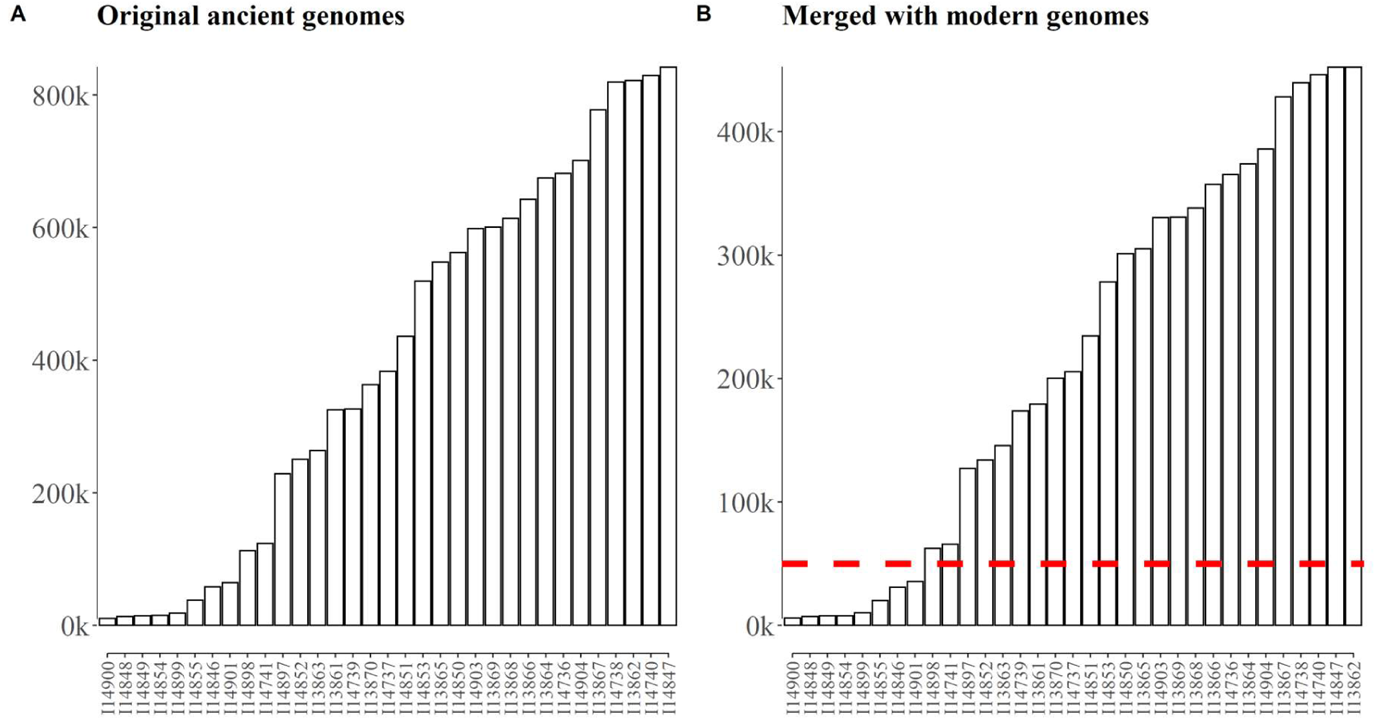
The number of genotyped SNPs in each Erfurt sample. (A) In the original ancient genomes, the mean and median number of SNPs were 402k and 383k, respectively (autosomes only). (B) After merging with the Human Origins dataset, the mean and median number of SNPs in the Erfurt samples were 219k and 205k, respectively. The horizontal dashed line indicates the cutoff defining the low-coverage samples (Figure S8).

**Figure S4.**
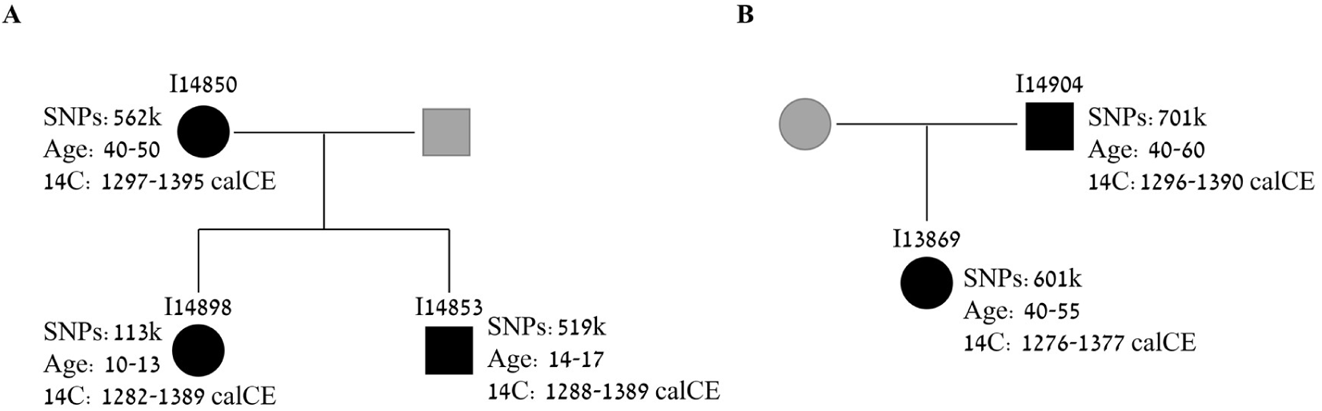
First-degree relatives. The figure shows the pedigrees of the two families we identified. Black symbols represent genomes we genotyped; gray symbols represent inferred family members. Circles: females; squares: males. For each sample, we indicate the sample ID, the number of genotyped SNPs, the estimated age, and the ^14^C date. (A) Family A, with a mother, a son, and a daughter. (B) Family B, with a father and a daughter.

**Figure S5.**
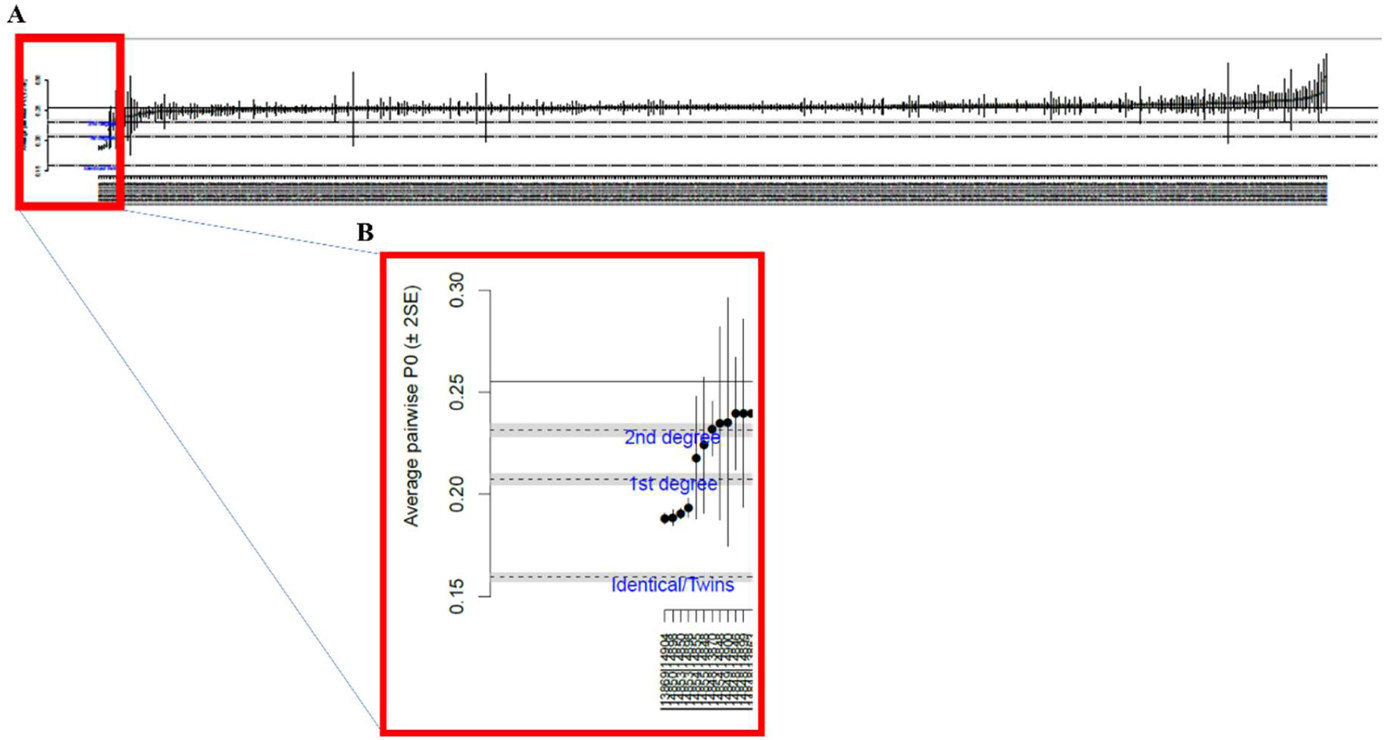
Detecting additional relatives. The figure shows the output of *READ* [29]. Each vertical line corresponds to one pair of individuals. The main figure shows all 33*32/2=528 possible pairs. The inset shows the first few pairs, seven of them with a point estimate of either first- or second-degree relationship. The y-axis shows the mean proportion (across 1Mb genomic windows) of non-matching alleles between pairs of individuals (P0). The horizontal solid line corresponds to the median P0 in the sample. The horizontal dashed lines correspond to the cutoffs for first-degree relatives, second-degree relatives, and unrelated individuals, and the horizontal gray bars correspond to their 95% confidence intervals [29]. The vertical lines for each pair represent two standard errors of the mean (across genomics windows). Two pairs of individual, I14855 and I14854, and the same I14855 and I14848, were estimated to be second-degree relatives, although the confidence intervals also included a first-degree relationship or no relationship. The value of P0 for I14854 and I14848 was slightly above the cutoff for a second-degree relationship. All three samples had low coverage (<40k SNPs).

**Figure S6.**
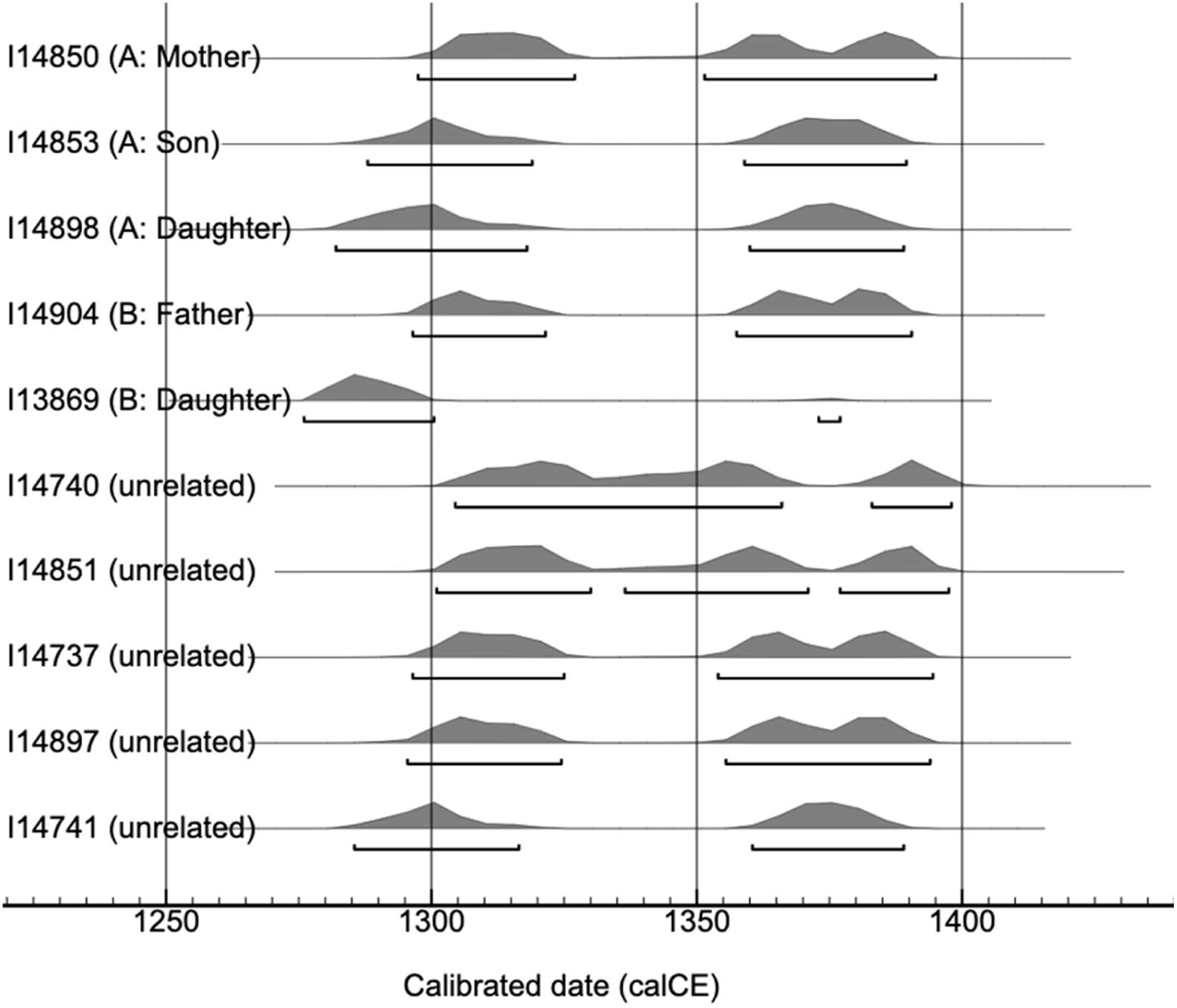
The distribution of estimated dates of the Erfurt samples based on ^14^C. The figure was generated by OxCal for ten samples that underwent radiocarbon dating. For each individual, the underlying bars denote time intervals that have a combined probability of 95.4%. The estimated dates are almost equally likely to predate or postdate the 1349 pogrom. One individual – I13869 – was inferred to be much more likely to date to the 13^th^ century. However, there was a small amount of probability density in the late 14^th^ century, and this individual is the daughter of I14904 (Figure S4), who is more likely to date to the 14^th^ century. This implies that I13869 might also date to the 14^th^ century, despite this event having a smaller probability.

**Figure S7.**
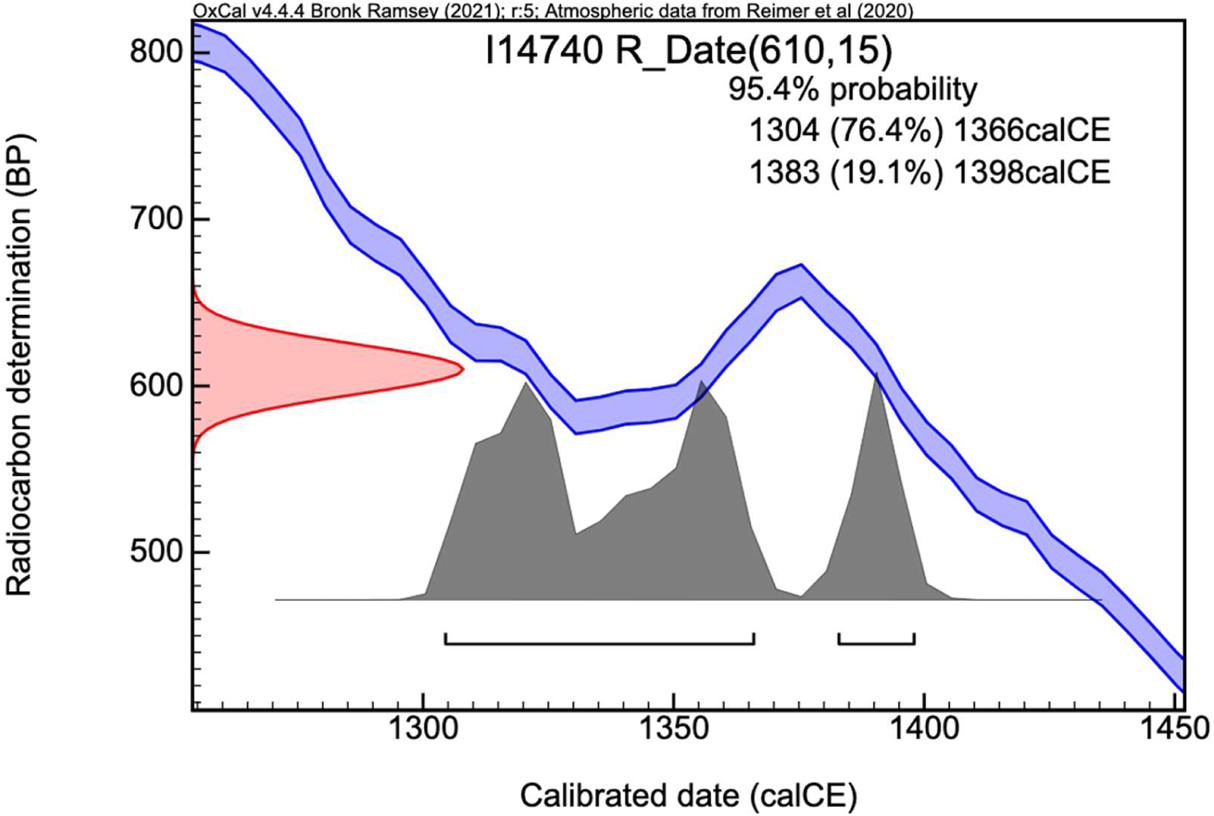
The radiocarbon calibration curve for a representative sample. We show a screenshot from OxCal for sample I14740 (see also Figure S6). The figure demonstrates the wiggle in the calibration curve throughout the 14^th^ century, which prohibits the definitive dating of the sample to before or after the 1349 pogrom.

**Figure S8.**
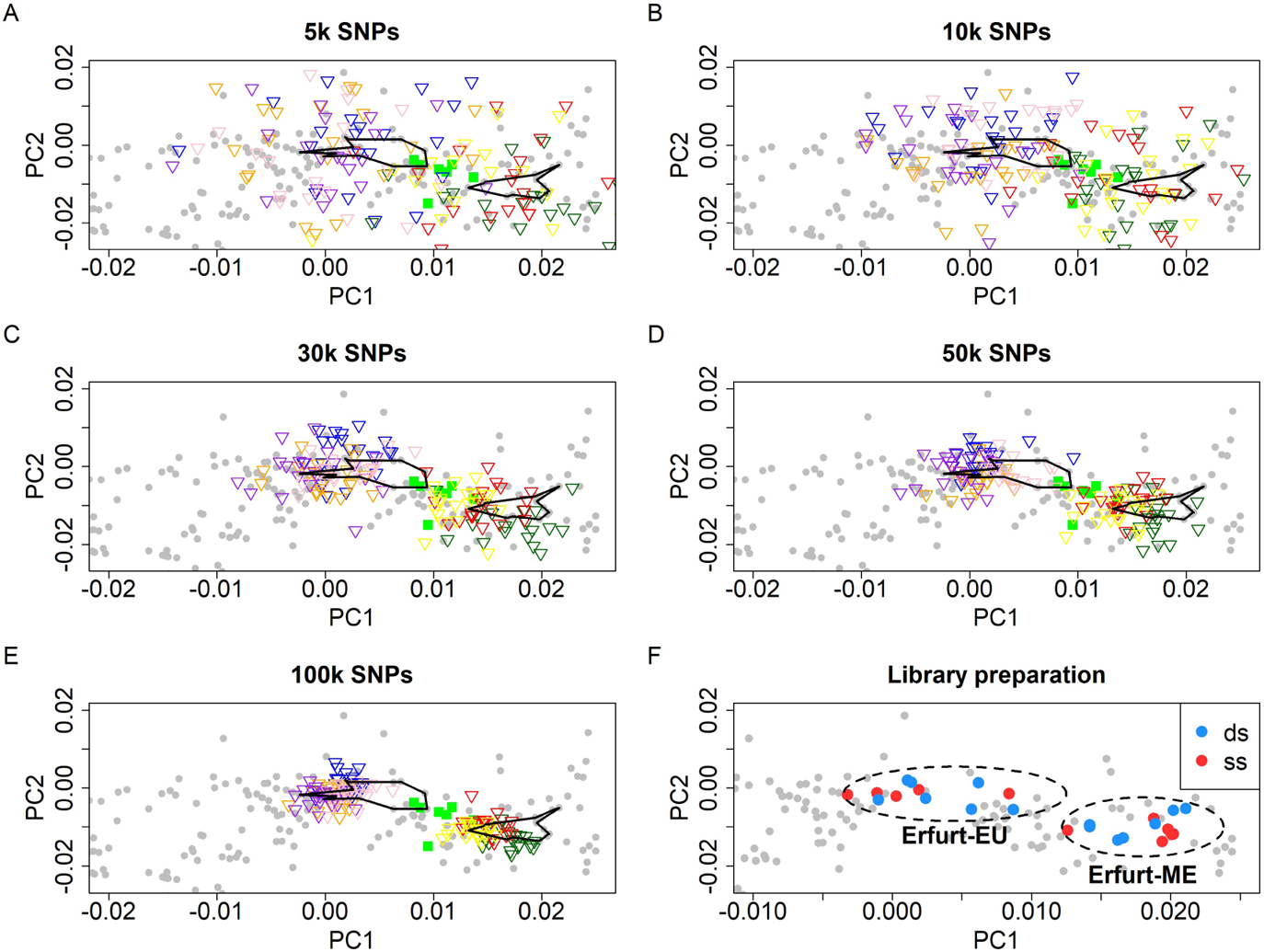
The effect of the coverage of the Erfurt samples on the PCA results. (A) – (E) To determine the minimal number of SNPs for a reliable PC projection, we down-sampled seven high-coverage EAJ (four of Erfurt-EU and three of Erfurt-ME) to different levels of coverage (5k, 10k, 30k, 50k, and 100k SNPs) and examined their location in PC space relative to the original samples. For each sample and for each coverage level, we generated 20 down-sampled copies, and projected them onto the West-Eurasian PC space, as in the main text. The down-sampled genomes are plotted as triangles colored based on their original sample ID. MAJ are plotted as filled green squares, and the original EAJ genomes are marked by two polygons corresponding to Erfurt-EU (to the left of MAJ) and Erfurt-ME (to the right of MAJ). All PCA plots are zoomed-in versions of the PCA in the main text. In downstream analyses of the PCA results, we only used samples with coverage 50k and above. At this coverage level, down-sampled genomes remain reasonably close to their original positions, and there is no overlap between samples originally designated as Erfurt-EU or Erfurt-ME. (F) A zoomed-in version of the PCA of the main text with EAJ samples marked by their type of library preparation (ss: single-stranded; ds: double-stranded). The difference in PC1 coordinates between the two treatments was not significant (P=0.95, two-tailed t-test).

**Figure S9.**
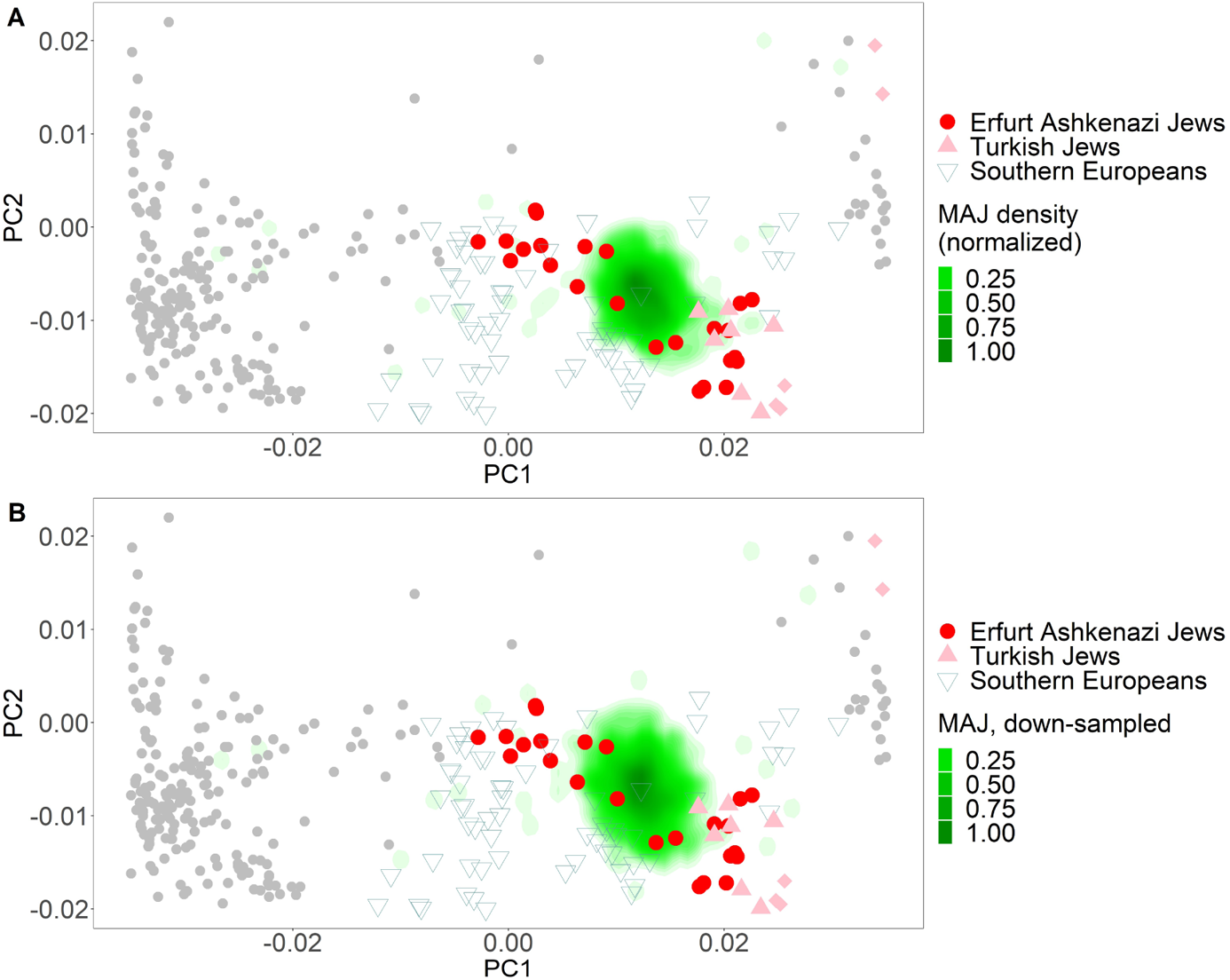
Projection a large modern AJ sample on the principal components space. (A) The plot is similar to the inset of Figure 1 of the main text, with two differences. (1) We projected both the Erfurt samples and the modern Ashkenazi samples on the principal components (PCs) that were learned using the Human Origins (HO) dataset. As in the main text, we learned the PCs using all West-Eurasian samples, but the figure only shows a subset of the space relevant for studying within-AJ structure. (2) We did not include in the analysis the seven modern AJ samples that were part of the HO dataset. For the modern AJ samples, we used the Ashkenazi Genome Consortium samples (n=544), down-sampled to the ≈470k HO SNPs. As the modern AJ sample is very large, we do not plot individual points but rather the density of points (Methods 3.1). As in Figure 1, the Erfurt samples show higher variability on the PC1 axis (the European/Middle Eastern cline) than modern AJ. (B) To demonstrate that the higher variability in EAJ is not due to their lower coverage compared to MAJ, we further down-sampled each MAJ sample to match the SNPs covered in an EAJ sample. We implemented that by arbitrarily ordering (randomly selected) 525 MAJ and the 25 non-low-coverage EAJ samples, and sequentially matching MAJ and EAJ samples, cycling over the EAJ samples until covering all modern samples. For each down-sampled MAJ genome, we further used a single, randomly-selected allele. Down-sampling did not qualitatively change the results.

**Figure S10.**
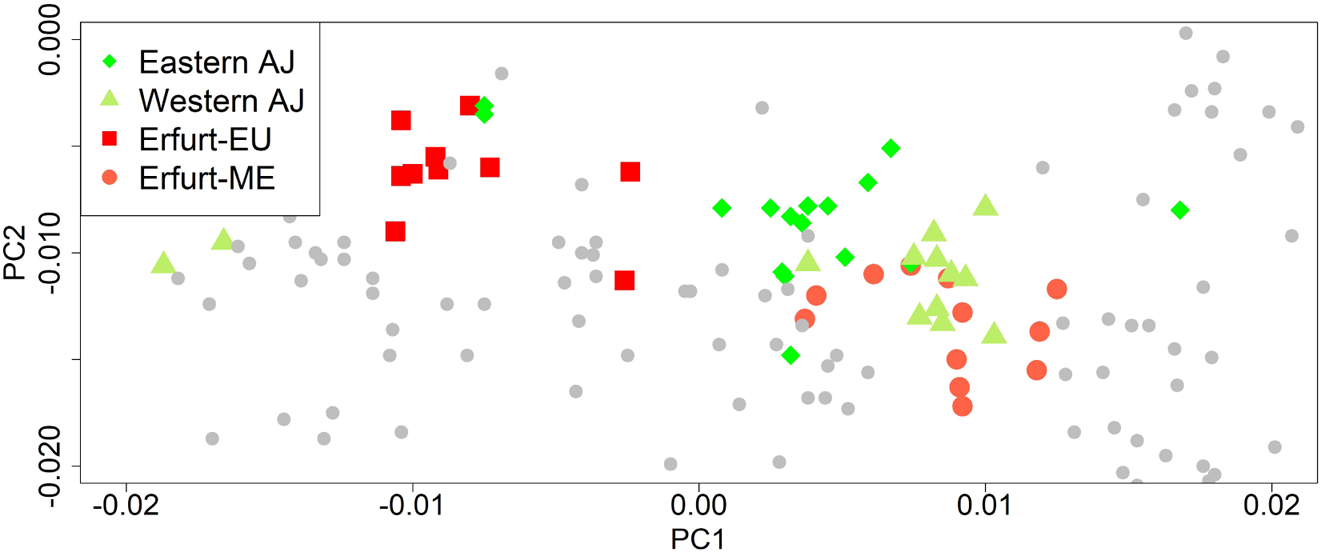
PCA with Eastern and Western modern AJ. To test whether the two subgroups of Erfurt correspond to modern AJ of Eastern European or Western European origin, we merged the Erfurt data with that of Behar et al. (2013) [40]. The merged dataset included about 246k SNPs. We ran the PCA using West-Eurasian samples, but the figure shows only a subset of the space relevant for studying within-AJ structure. Both EAJ and MAJ were projected on the PC plane. The results show that Western MAJ overlap with Erfurt-ME and that Eastern MAJ have slightly more EU ancestry, although both groups cluster primarily next to Erfurt-ME.

**Figure S11.**
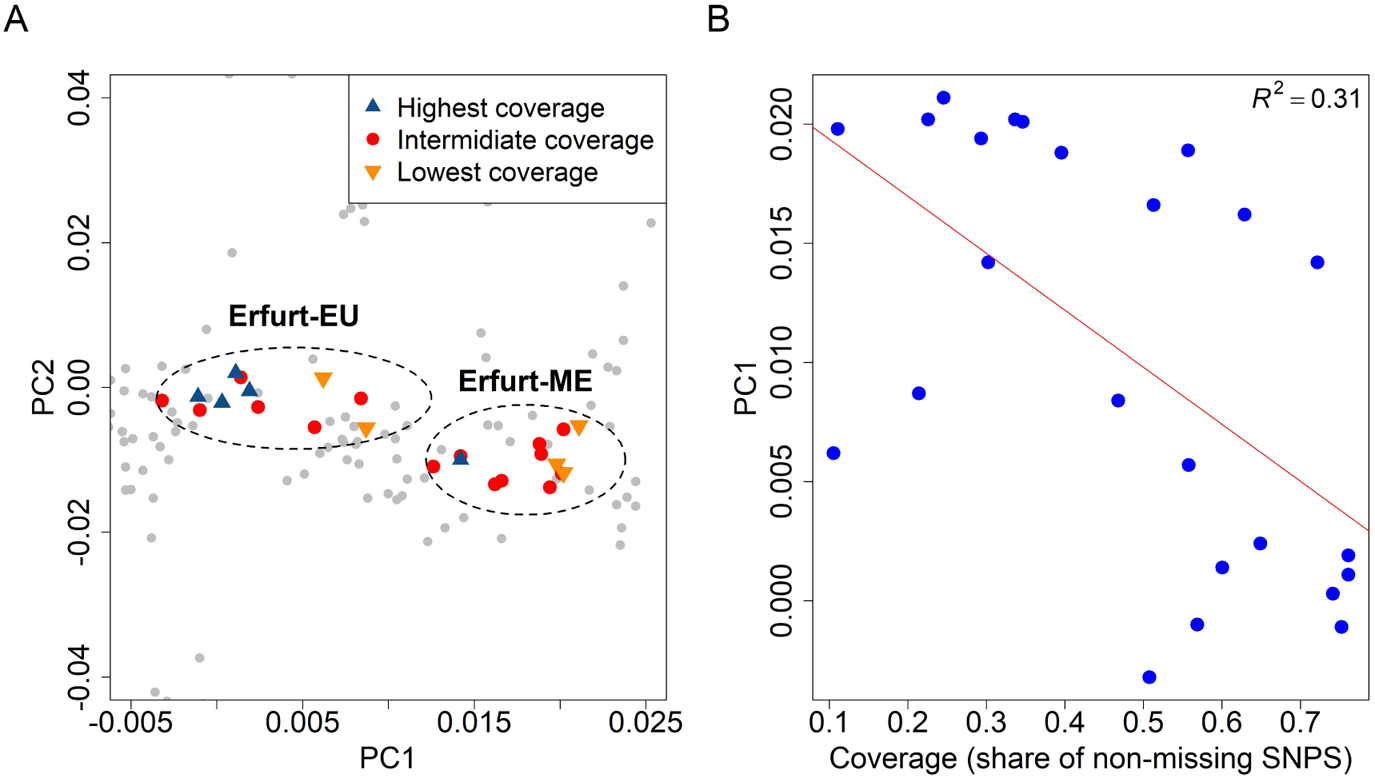
The association of the sequencing coverage with the PCA placement. (A) The figure shows a zoomed-in version of the PCA of the main text, where Erfurt samples are labeled by their coverage. The five individuals with the highest and lowest coverage are marked as “highest coverage” and “lowest coverage”. [The PCA plot does not include the eight samples with <50k SNPs. In other words, the “lowest coverage” samples are the lowest only among the samples that were used in the PCA.] Four of the five highest coverage samples are part of the Erfurt-EU group. (B) The PC1 coordinate vs the coverage level, along with a regression line.

**Figure S12.**
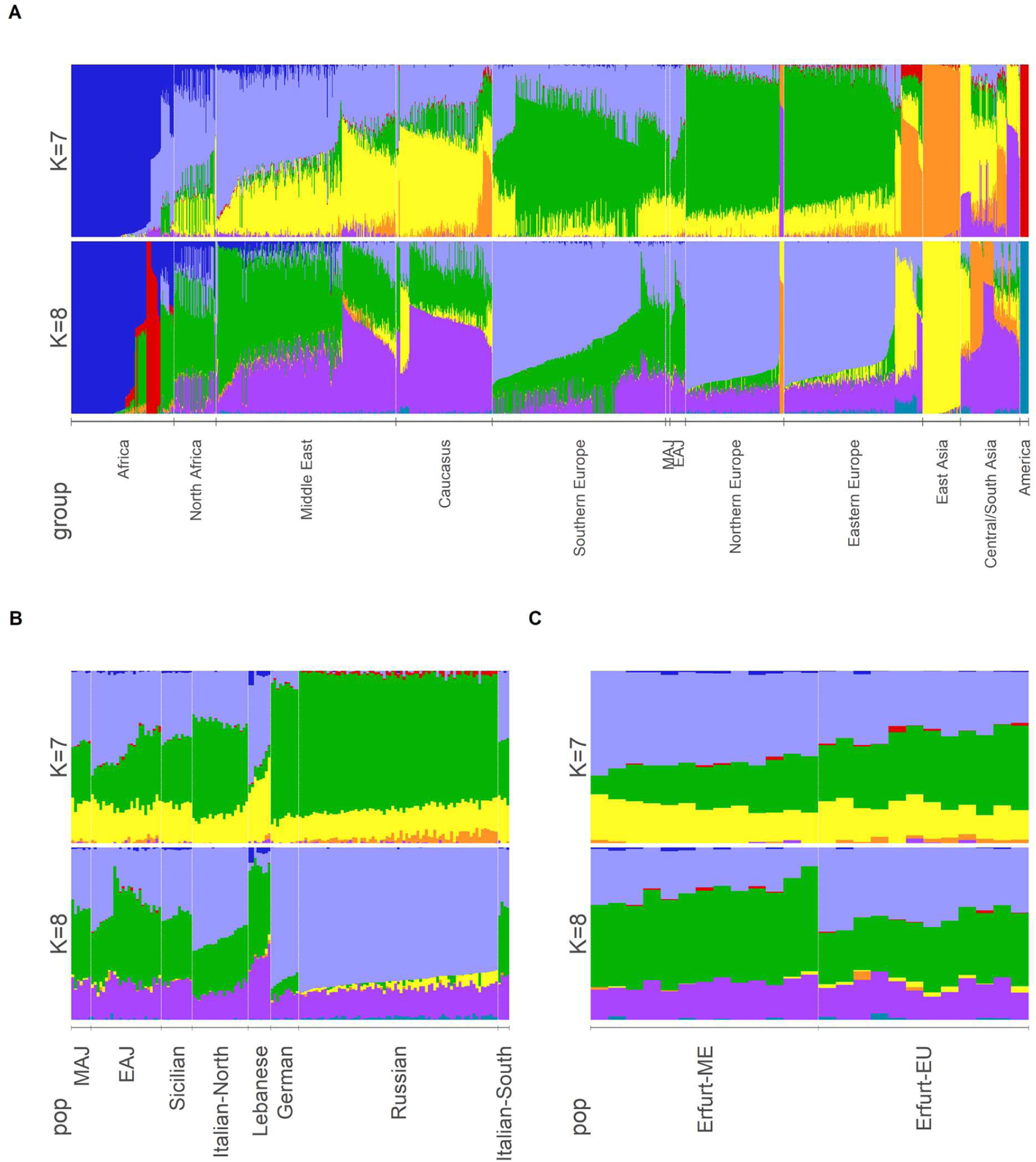
ADMIXTURE results. (A) ADMIXTURE results for all populations, grouped by regions, for *K* = 7 and *K* = 8 ancestral components. We also show zoom-in on populations relevant for our study (B) and on EAJ alone (C). In (C), we divided EAJ into Erfurt-EU and Erfurt-ME and sorted the samples by the European component (green) in *K* = 7.

**Figure S13.**
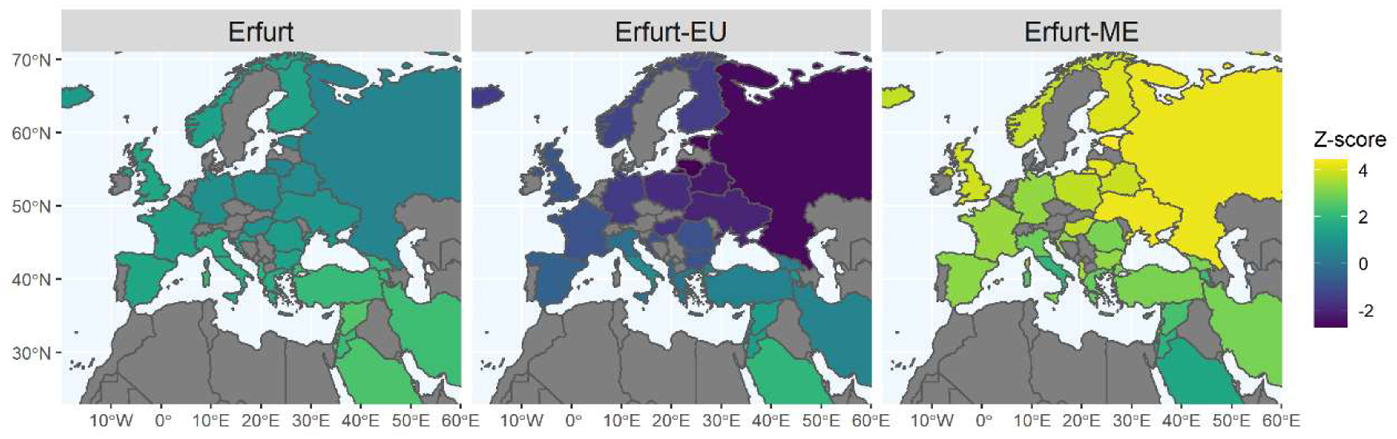
Tests of the form f_4_(MAJ, EAJ; X, chimp). X represents any non-Jewish West-Eurasian population. Each country on the map was colored based on the Z-score for deviation from zero of the f_4_-statistic when replacing X with the local population. Gray represents countries that were not tested. In the middle and right columns, EAJ were replaced with Erfurt-EU and Erfurt-ME, respectively.

**Figure S14.**
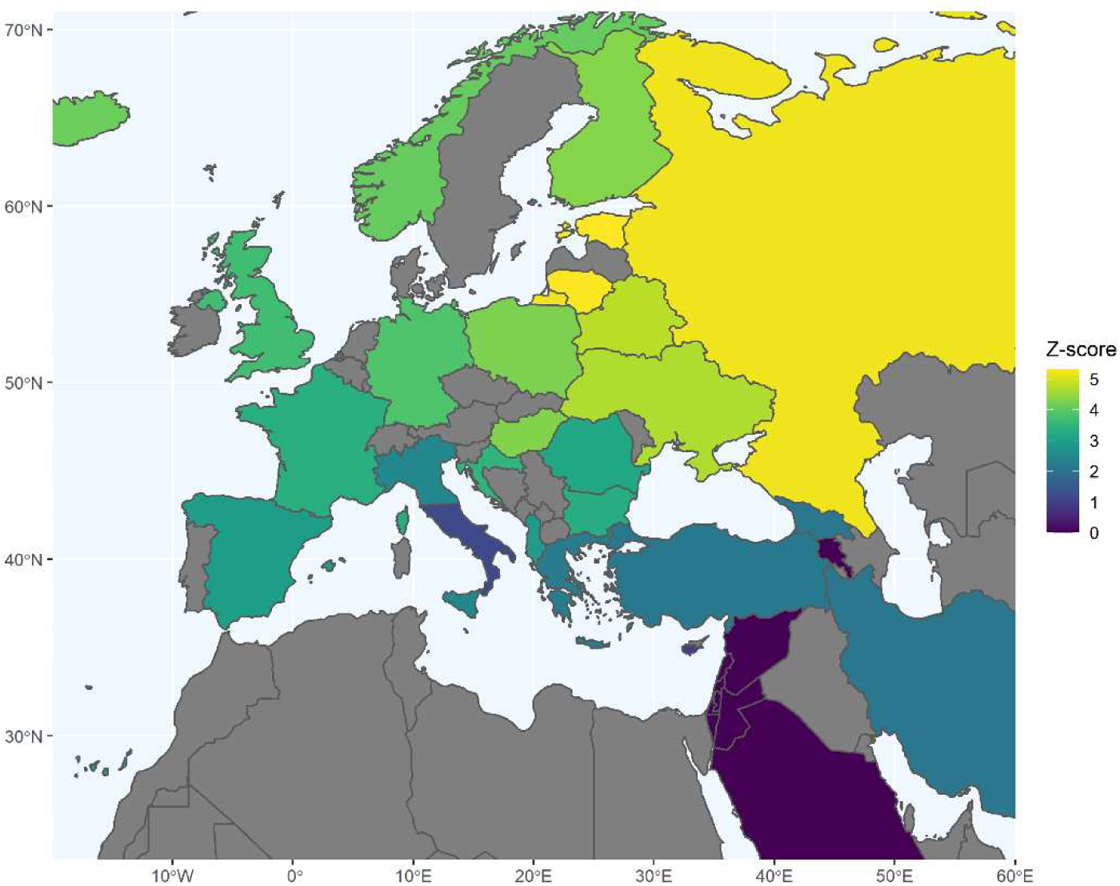
Z-scores for deviation from zero of tests of the form f_4_(Erfurt-EU, Erfurt-ME; X, chimp). X represents any non-Jewish West-Eurasian population. Each country in the map was colored based on the Z-score of the f_4_ test when replacing X with the local population. Gray represents countries that were not tested.

**Figure S15.**
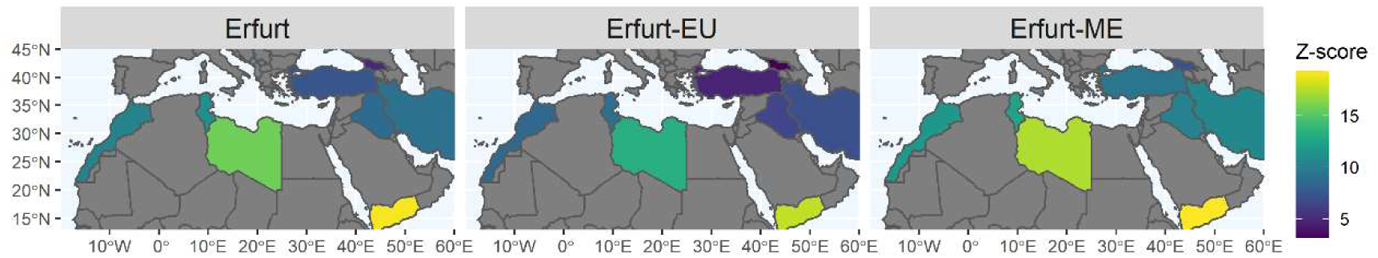
Z-scores for deviation from zero of tests of the form f_4_(MAJ, X; EAJ, chimp). X represents Jewish non-Ashkenazi populations. The location of each Jewish population on the map is represented by its origin in the diaspora. In the middle and right columns, EAJ were replaced with Erfurt-EU and Erfurt-ME, respectively. In all analyses, the Z-score was positive and >3, indicating that MAJ are the closest Jewish population to EAJ.

**Figure S16.**
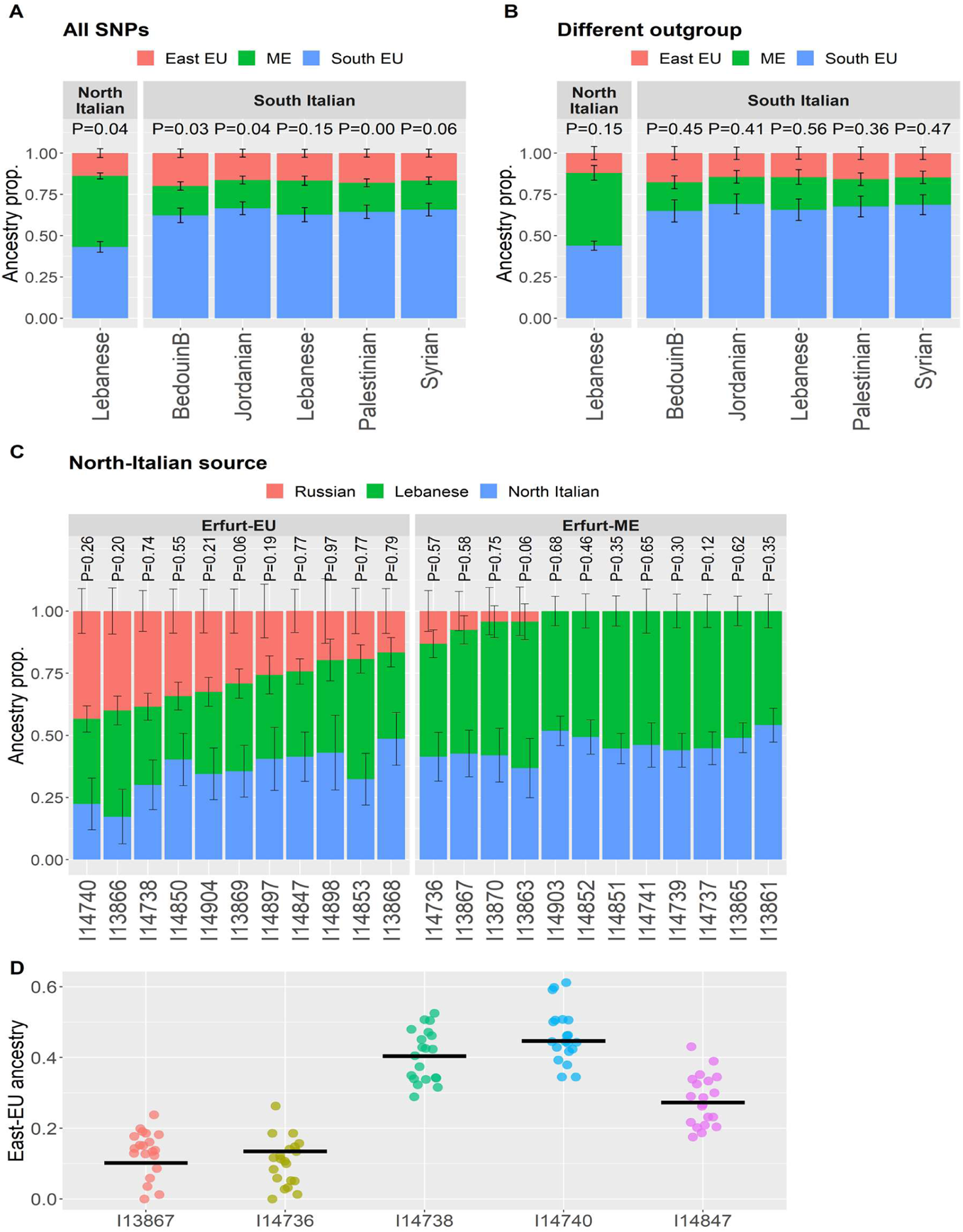
Robustness tests for *qpAdm*. Panels (A) and (B) are the same as in Figure 2A of the main text, showing models for the ancestry of EAJ, with the following changes. (A) We used all SNPs instead of only transversions. (B) We used the Ami population as the outgroup instead of Mbuti. The plots present the models that were plausible in the main analysis. Panel (C), which is analogous to Figure 2B, shows a model for the ancestry of single individuals (labeled by their IDs). The sources were Russians, Lebanese, and North-Italians (instead of South-Italians in the main text). Two individuals could not be modeled using these sources and are not presented. As in the main analysis, Erfurt-EU individuals have a substantial East-EU component that is missing from most Erfurt-ME individuals. The individual-level models were estimated using all SNPs. (D) We sought to determine whether the absence of Eastern European ancestry in some Erfurt-ME individuals might be due to their lower coverage (Figure S11). We used five high-coverage samples (two Erfurt-ME and three Erfurt-EU) and down-sampled each genome 20 times to 100k random SNPs, as in Figure S8. In the plot, we compare the East-EU ancestry proportion inferred by *qpAdm* in the original samples (horizontal black lines) to those inferred in their down-sampled versions (colored dots). Two samples from Figure S8 were not used: one had no East-EU ancestry, and one could not be modeled using the given sources. We used all SNPs and the same sources as in Figure 2B: South-Italians, Lebanese, and Russians. The results show that the inferred proportion of East-EU ancestry is reasonably robust to down-sampling.

**Figure S17.**
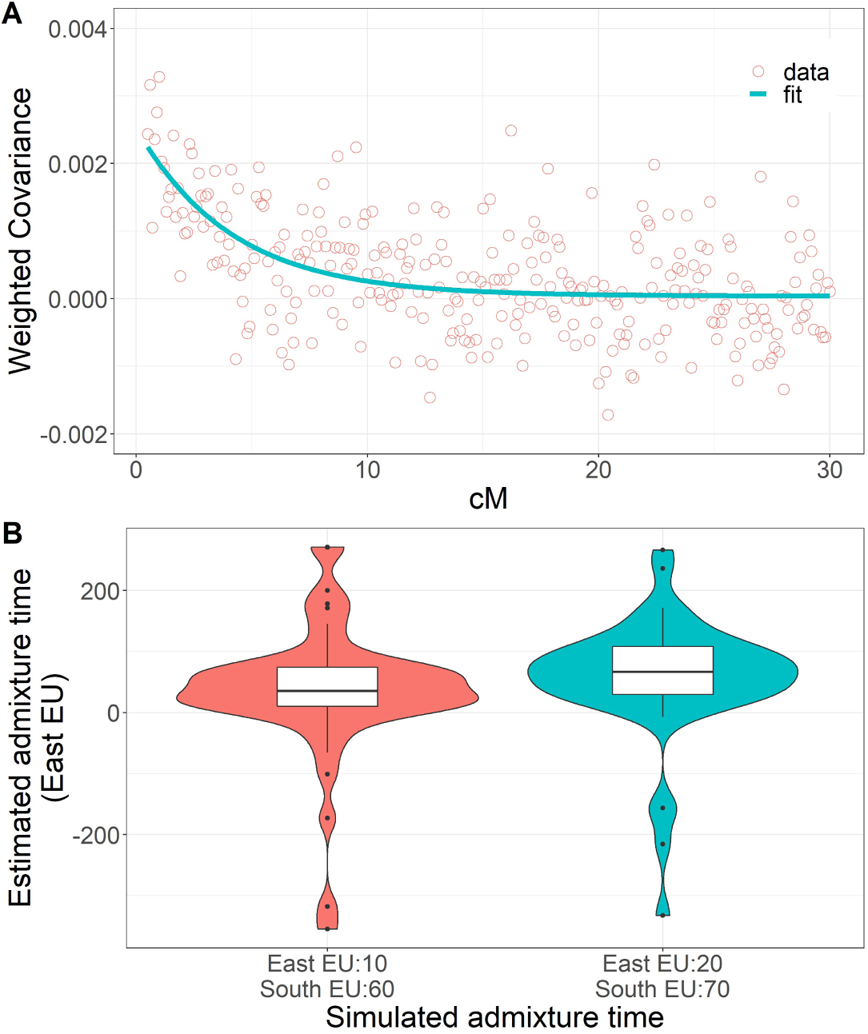
*DATES* results. **(A)** We used *DATES* [48, 49] to estimate the admixture time between AJ ancestors and Eastern Europeans. The first source population was Russians and the second was Middle Easterners + Southern Europeans. We used Erfurt-EU as the admixed population, given that (based on the *qpAdm* models of Figure 2B of the main text), Erfurt-ME samples have little East-EU ancestry. The coral circles represent the observed weighted covariance (described in [48, 49, 116]) between the genotypes of SNPs in each genetic distance apart. The blue line represents the least squares fit to an exponential decay. Based on the decay rate, it is estimated that admixture occurred 22.6 ± 8.1 generations before the time when the Erfurt-EU individuals lived. **(B)** We tested the performance of *DATES* on genomes simulated under two admixture events: the first between Middle Easterners (35%) and Southern Europeans (65%), and the second with Eastern Europeans (15%). In the first scenario we examined, the older admixture time was 60 generations ago and the recent admixture was 10 generations ago. In the second scenario, the admixture events happened 70 and 20 generations ago, respectively. For both scenarios, we matched the sample size and the number of SNPs to those of the real Erfurt-EU data (Methods 5.2) and repeated the simulations 50 times. For each set of simulated genomes, we then used *DATES* (with the same parameters as in the real data analysis) to estimate the time of the admixture event with Eastern Europeans (Methods 5.2). The x and the y axes represent the true and inferred admixture times, respectively. The plot shows the densities of the *DATES* estimates under the two scenarios. In the box plots, the bold horizontal line represents the median, the borders of the box are the first and third quartiles, and the vertical lines extend to the most extreme value no more distant than 1.5x the inter-quartile range from the quartiles. We omitted from the plot one data point for which the inferred admixture time was 1,915 generations ago. The distribution of the estimated admixture times is extremely wide under both scenarios, suggesting that *DATES* cannot reliably infer the admixture time for the real Erfurt-EU data.

**Figure S18.**
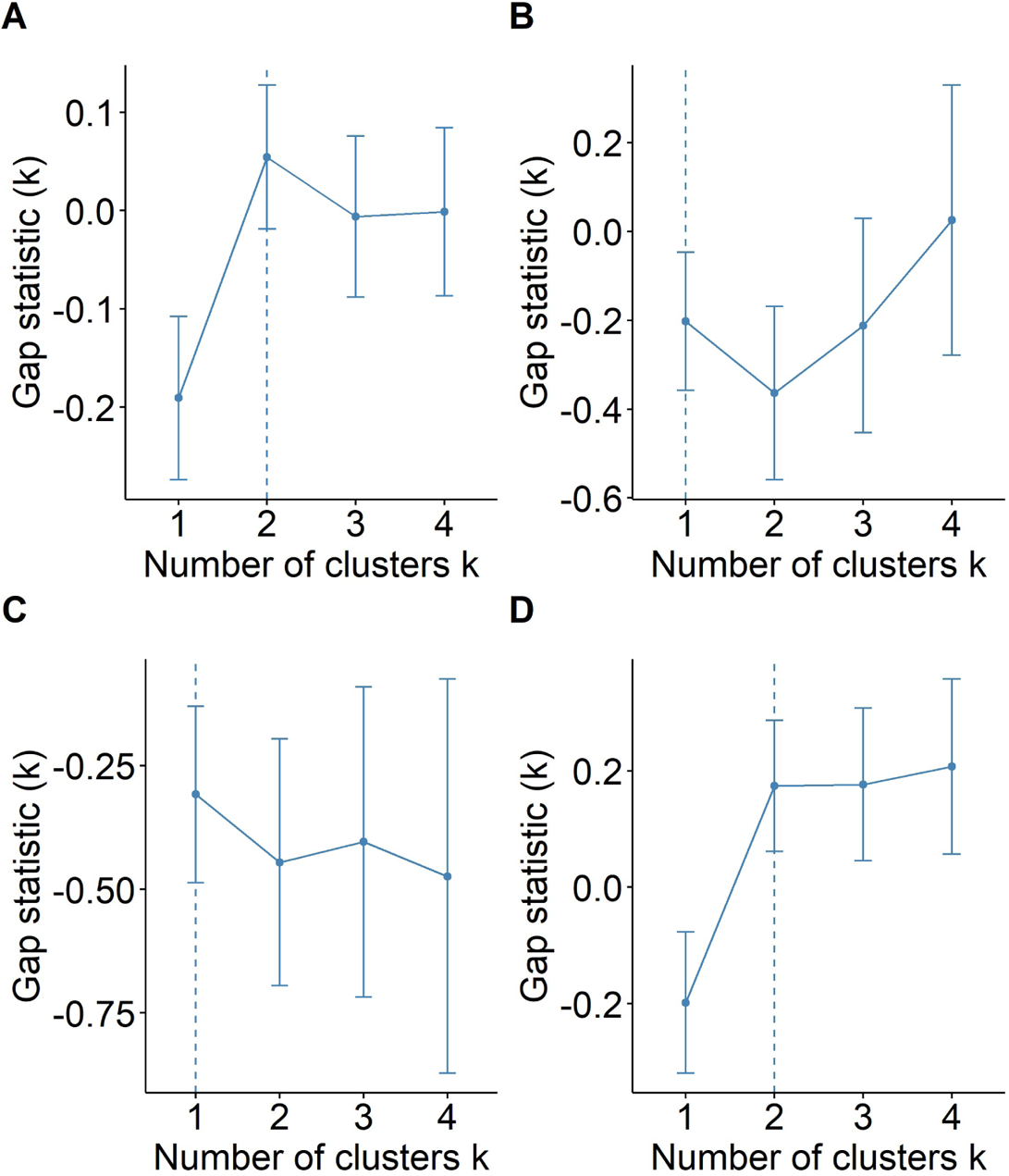
The number of clusters in EAJ based on the first two PCs. We used the gap statistic (Methods 6.1) to infer the optimal number of clusters (dashed line) for EAJ (A). As a control, we also inferred for the optimal number of clusters for modern AJ ((B); one cluster expected); Moroccan Jews ((C); one cluster expected); and modern AJ and Moroccan Jews together ((D); two clusters expected).

**Figure S19.**
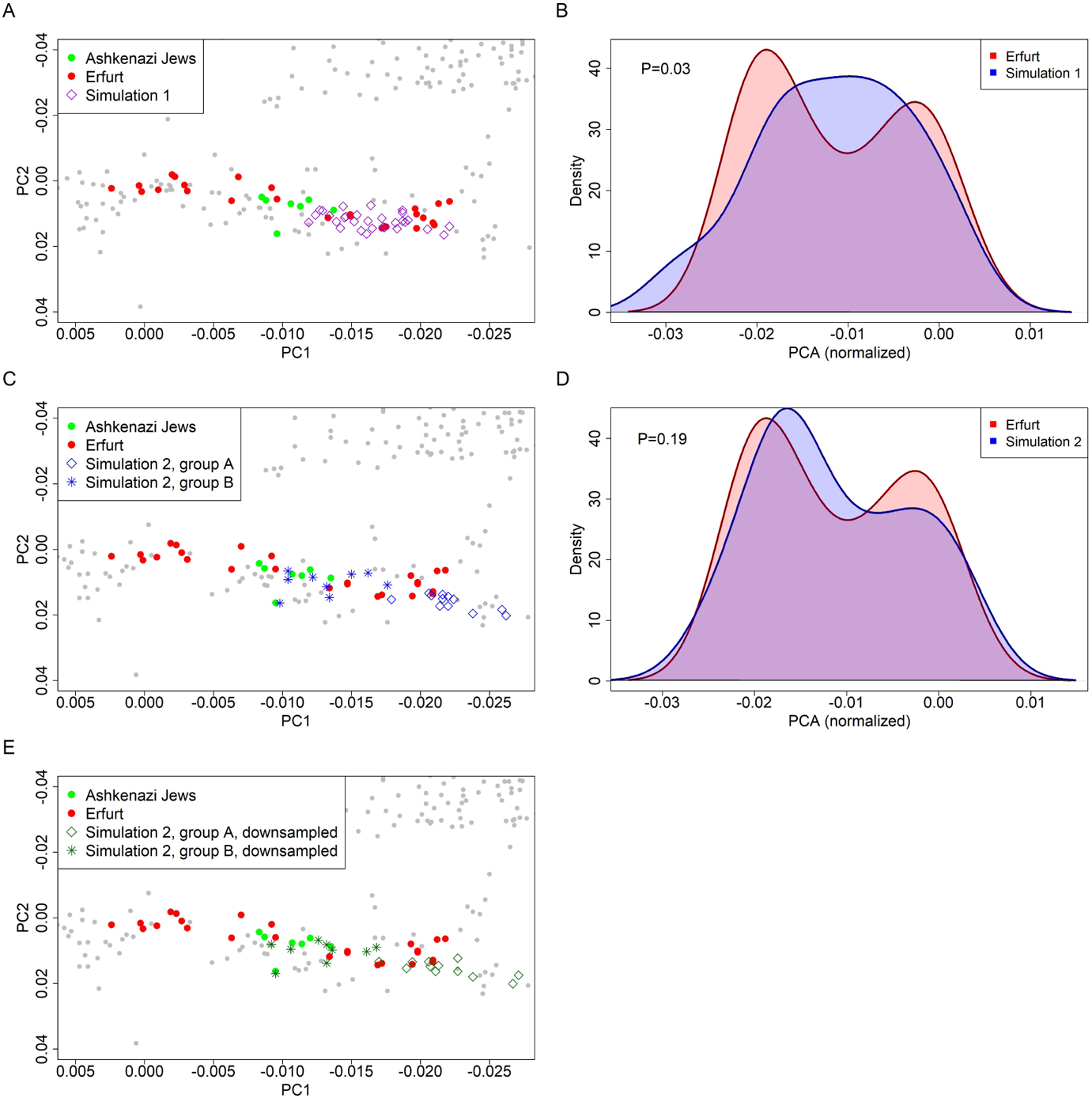
PCA of genomes simulated under two admixture scenarios and a comparison to EAJ. (A) PCA results for simulations of a single group that has experienced recent admixture between Middle Eastern, Southern European, and Eastern European sources. Both simulated genomes and EAJ genomes were projected on the PC plane. (B) A comparison between the distributions of PC1 of the simulated genomes (from (A)) and EAJ. The distribution of the simulated data was shifted and scaled to match the mean and variance of the EAJ data. The EAJ distribution is bimodal, and does not fit the simulated data. (C) PCA results for simulations of two groups, one that has experienced admixture between Middle Eastern and Southern European sources, and one that had additional admixture with Eastern Europeans. (D) A comparison between the distributions of PC1 of the simulated genomes (from (C)) and EAJ. The distribution of the simulated data was shifted and scaled to match the mean and variance of the EAJ data. Here, the distribution of the simulated data is also bimodal. (E) PCA results for the two-group simulation after down-sampling the simulated genomes to match the coverage of the Erfurt samples (Methods 6.3). The results are qualitatively similar to those of (C).

**Figure S20.**
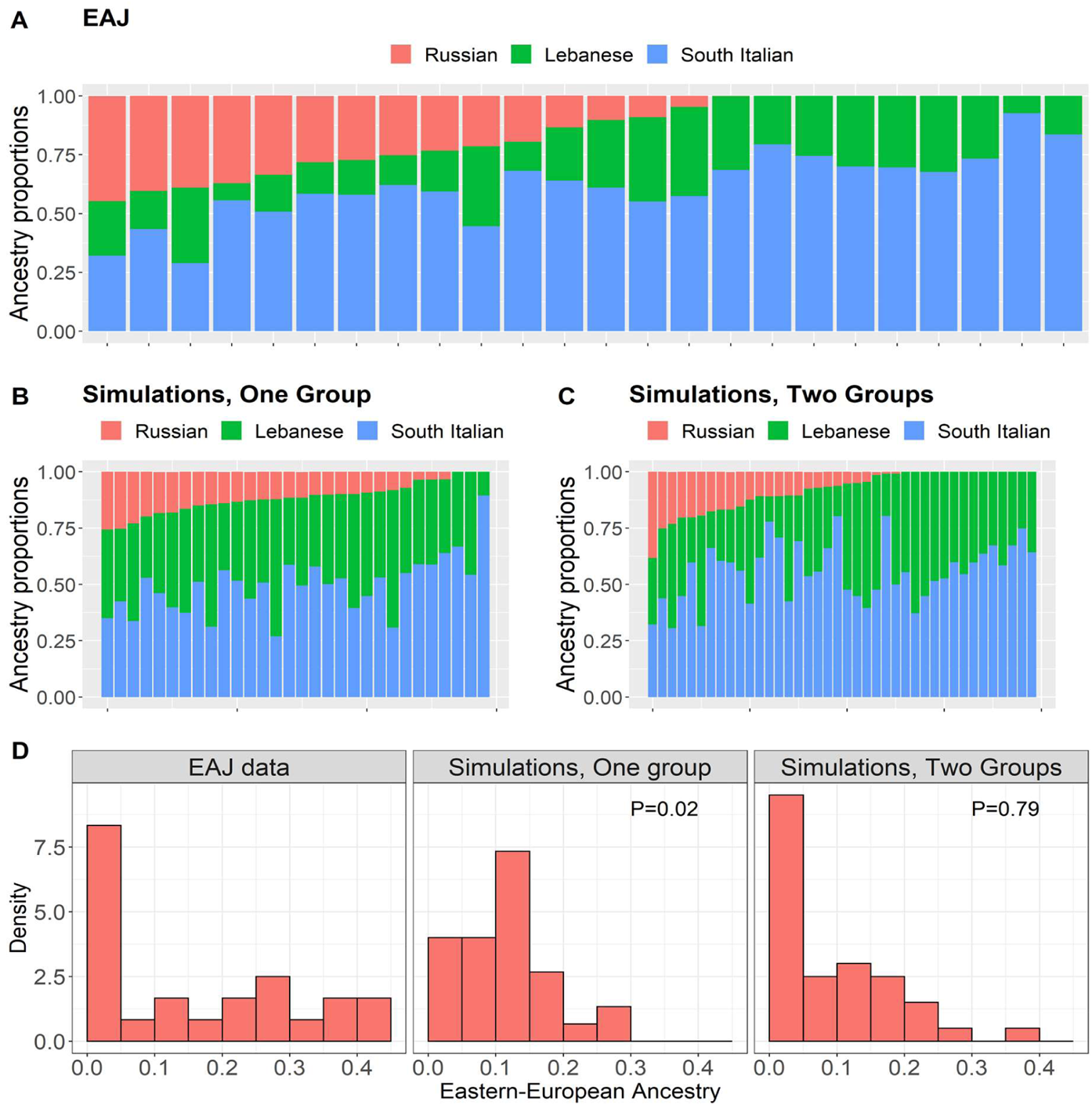
*qpAdm* results for genomes simulated under two admixture scenarios and a comparison to EAJ. (A) *qpAdm* results for the real Erfurt data with Lebanese, South-Italians, and Russians as sources. This panel is identical to Figure 2B of the main text. (B) *qpAdm* results for the single-group simulations. (C) *qpAdm* results for two-group simulations. (D) The distribution of the Eastern European ancestry proportions in the real EAJ data, in the single-group simulations, and in the two-group simulations. The proportion of individuals with no East-EU ancestry in the real EAJ data is significantly different from that of the single-group simulation (P=0.01; permutation test, randomly shuffling the labels of simulated and real data points), but not from that for the two-group simulation (P=0.78).

**Figure S21.**
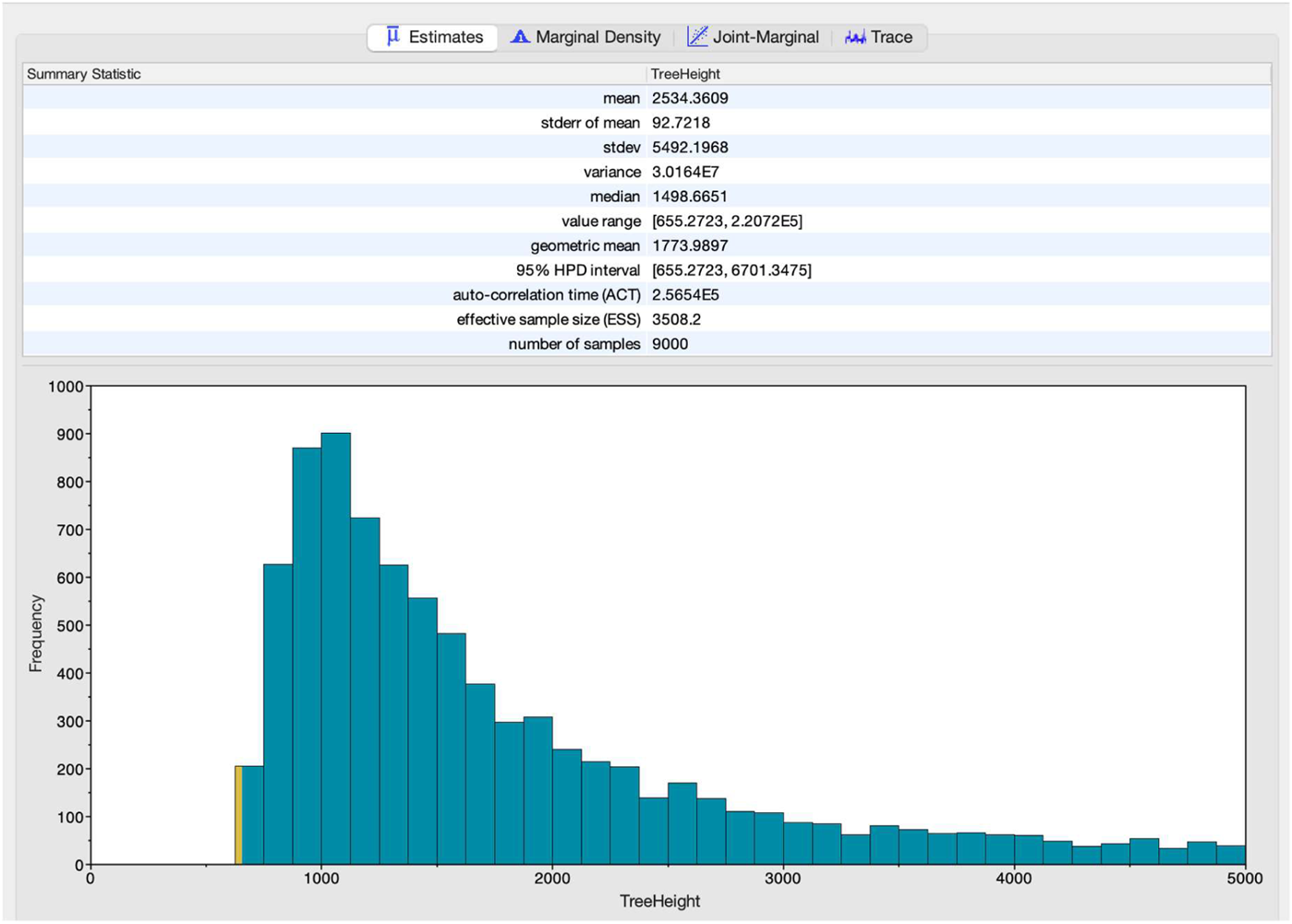
The posterior distribution of the mtDNA tree height (time to the most recent common ancestor (TMRCA)) based on ancient and modern K1a1b1a carriers. The plot shows a screenshot of the *Tracer* software showing the output of the *BEAST* analysis, as described in Methods 7.2. Briefly, we used an alignment of the mtDNA sequence of 11 EAJ and 107 MAJ K1a1b1a carriers. We ran *BEAST* with a strict clock, Gamma distributed mutation rates, and a skyline population size prior. The effective sample size (ESS) was 3508, and the total number of samples from the posterior was 900. The median posterior TMRCA was 1499 years ago, with a 95% highest posterior density (HPD) interval 655-6701 years ago. Other characteristics of the distribution are shown above the plot.

**Figure S22.**
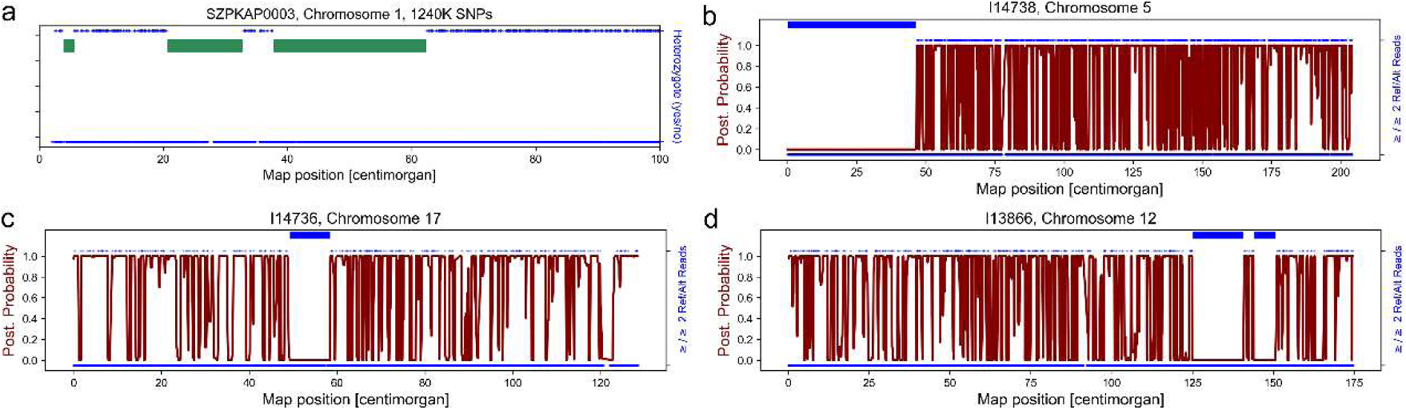
Visual inspection of runs of homozygosity (ROH) segments in modern and ancient genomes. (A) The inferred ROH segments (green bars, called with *bcftools/ROH*; Methods 8.2) along a subset of chr1 in one modern AJ individual. We considered only bi-allelic SNPs included in the “1240” SNP panel. Blue dots at the top (bottom) of the panel show the positions of heterozygous (homozygous) sites. It can be seen that the inferred ROH segments are depleted of heterozygous sites. (B-D) The inferred ROH segments (blue bars, inferred using *hapROH*; Methods 8.1) in three chromosomes from three Erfurt individuals. The red lines show the posterior (“post.”) probability estimated by *hapROH* that a SNP is in a non-ROH state given the data. The blue dots show sites that were covered by at least one read (Methods 8.1). A dot is plotted at the top of the panel whenever the reads covered both alleles, which suggests heterozygosity. The inferred ROH segments are again depleted of these putatively heterozygous sites.

**Figure S23.**
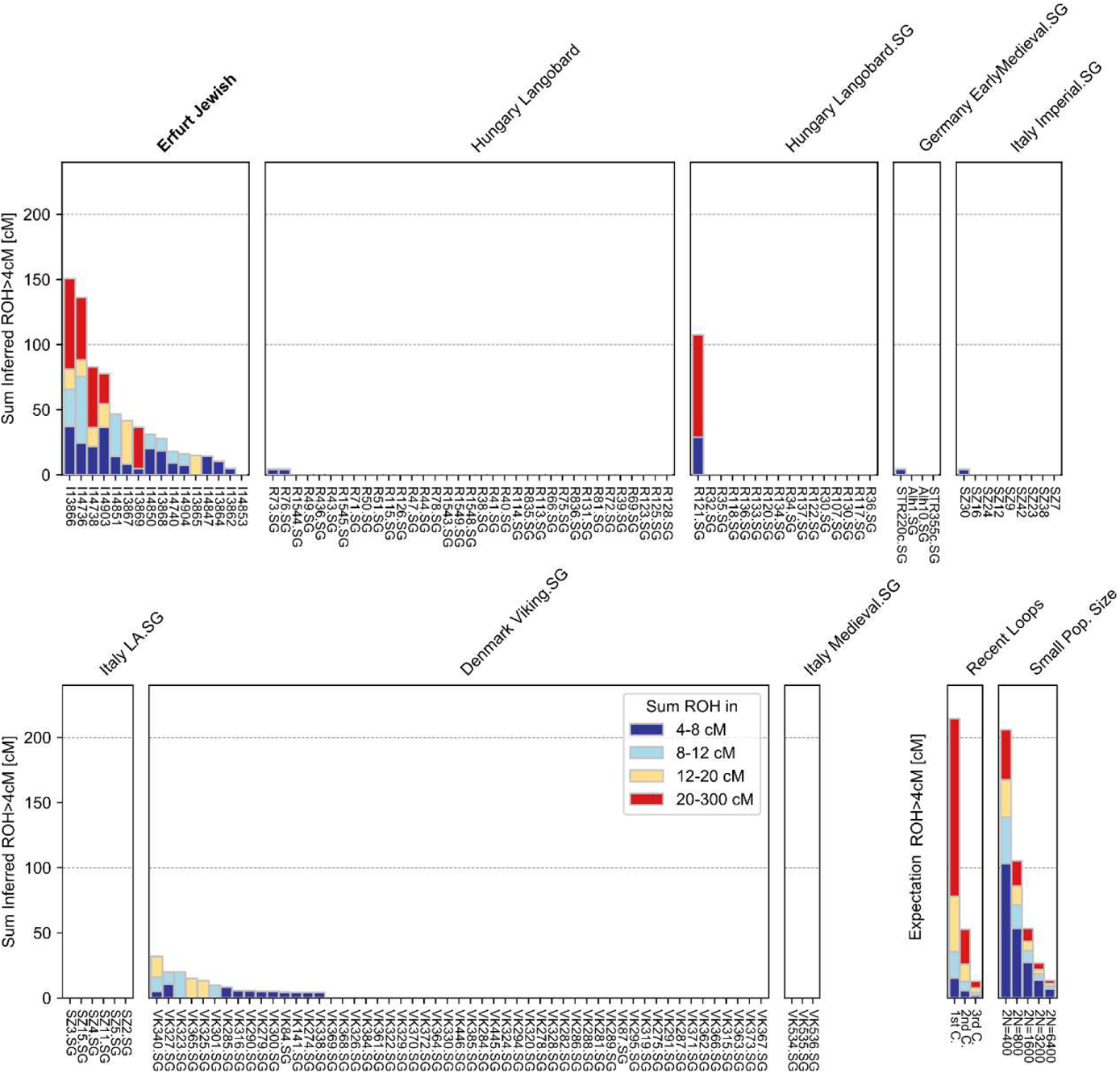
A comparison of the abundance of runs of homozygosity (ROH) across ancient European populations from the past two millennia. Each bar represents one individual, and individuals are grouped by population labels. We show the sum of the lengths of ROHs in four length bins (see legend). On the bottom right, we demonstrate the expected sum of ROH lengths for individuals whose parents are close relatives (first, second, and third cousins; “recent loops”), as well as for individuals from a population of a given constant effective size (*N* is in number of diploid individuals; “small pop. size”). See [58] for details. The Erfurt samples have substantially longer ROHs compared to all other populations. SG: shotgun sequencing. The Hungary Langobard data (SNP enrichment and SG) is from [60]. The Germany Early Medieval data is from [45]. The Italy Imperial, Late Antiquity (LA), and Medieval data is from [47]. The Denmark Viking data is from [61].

**Figure S24.**
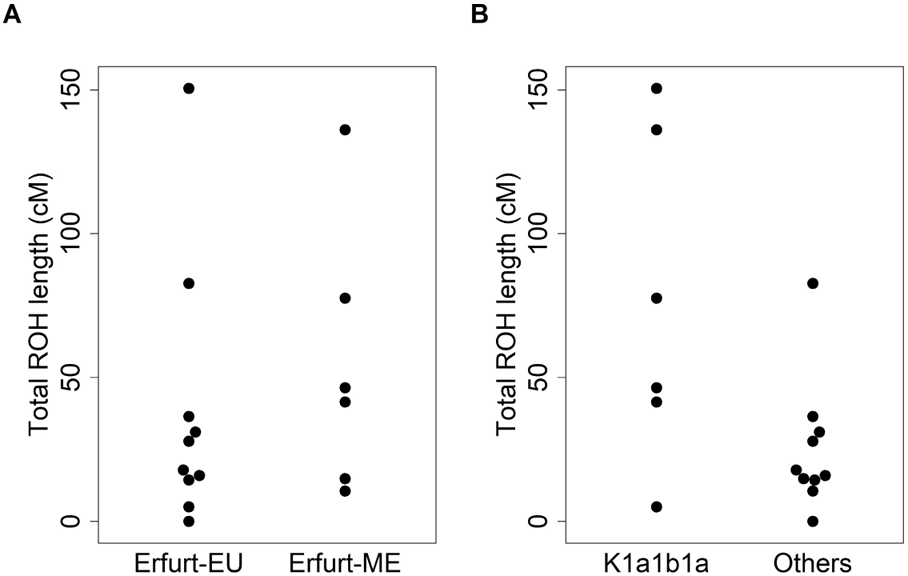
ROH segments across EAJ sub-groups. For each of 16 high-coverage (>400k SNPs) EAJ genomes, we computed the total length of ROH segments longer than 4 cM. (A) The total ROH length (cM) per genome in Erfurt-EU and Erfurt-ME individuals. The distribution is similar between the groups (P=0.43; two-tailed Wilcoxon test). (B) The total ROH length per genome was greater in K1a1b1a carriers compared to all other samples (P=0.03; one-tailed Wilcoxon test).

**Figure S25.**
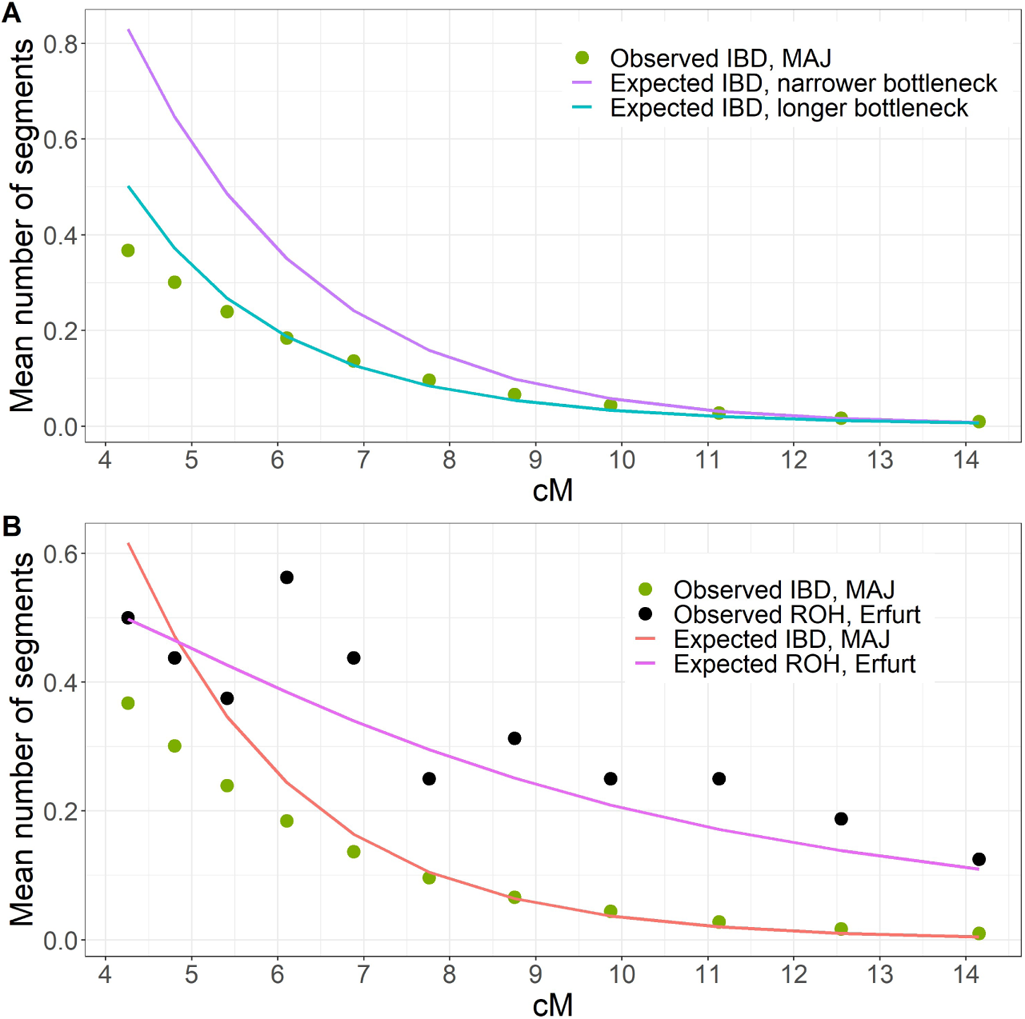
Demographic models that did not fit the modern IBD data. (A) The mean number of IBD segments (per pair of haploid genomes) across length bins in modern AJ is shown in circles. The purple and teal lines show the expected counts (Methods 8.6) as predicted by models having a narrower or a longer bottleneck, respectively (Figure 3D; Table S10, models (E) and (F)), as compared to the model inferred using modern IBD. These models, in particular the narrower bottleneck model, did not fit the modern data well. (B) We plot the observed number of IBD segments in MAJ and ROH segments in EAJ in green and black circles, respectively. The expectations based on the single-population joint-likelihood model, as described in Table S10, model (G), are shown in red and pink lines, respectively. The expected number of short IBD segments in MAJ is overestimated by the model.

**Figure S26.**
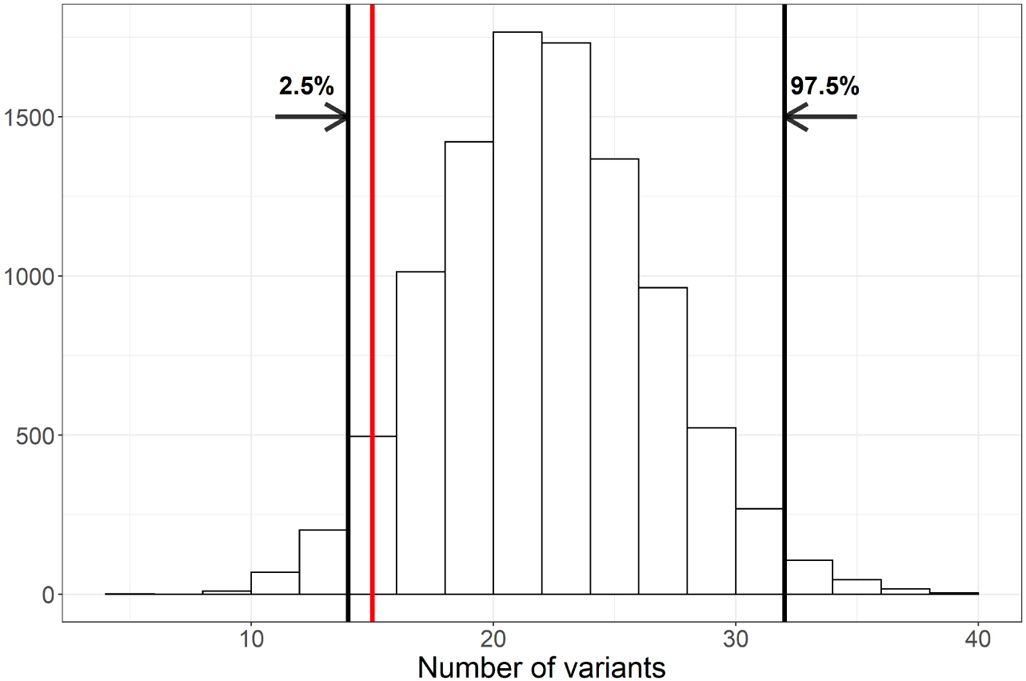
Simulations for the expected number of founder alleles in EAJ under modern allele frequencies. In each iteration and for each founder SNP, we drew a minor allele count as a binomial with *n* equals to the number of (pseudo-haploid) EAJ individuals that were genotyped in that SNP and *p* equals the allele frequency in MAJ (from *gnomAD* [76]). The figure shows the distribution of the number of founder SNPs with at least one minor allele across 10,000 runs. We show the 2.5- and 97.5-percentiles in black vertical lines, and the observed number in EAJ in red.

**Figure S27.**
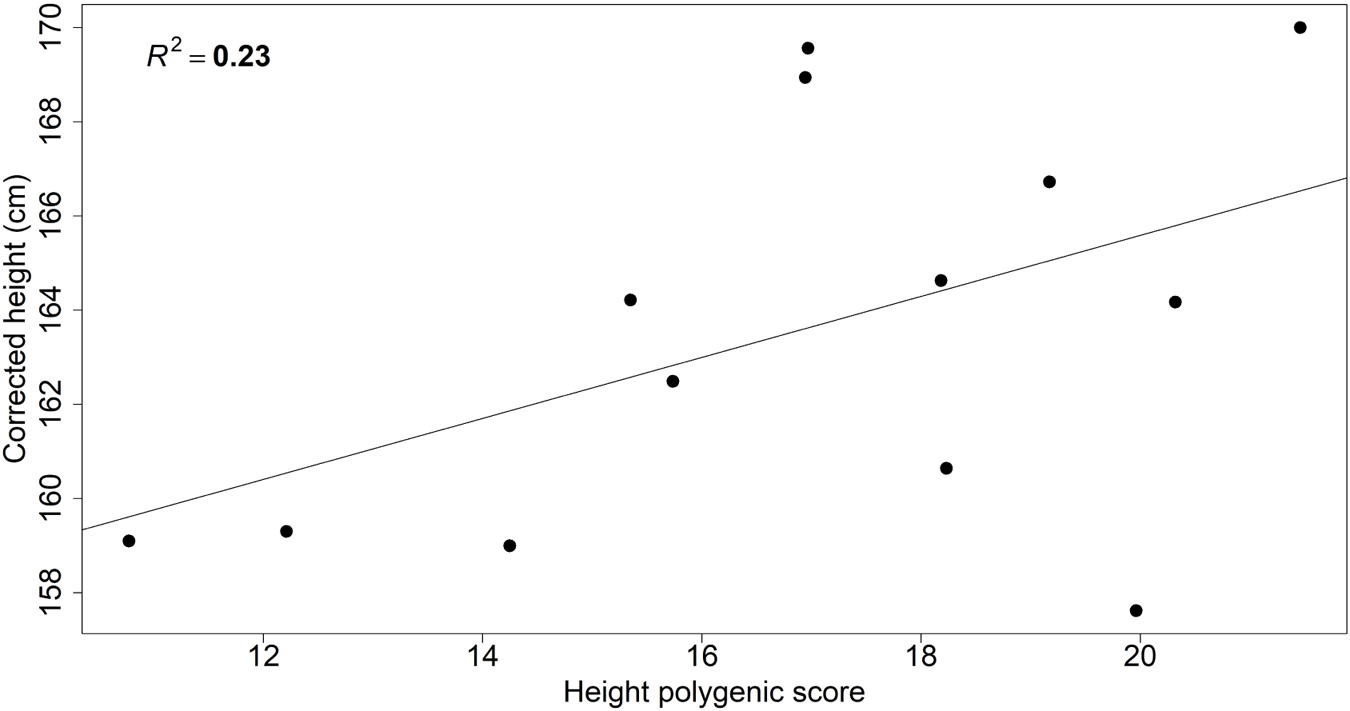
Correlation between height and polygenic score. For each individual, we plot the estimated height (mean over all available estimates, shifted up by 9.84 cm for females (the empirical mean difference between the sexes); Table S14; Methods 10.1) and the polygenic score for height based on summary statistics from [81] (Methods 10.1). We also plot the linear regression line. The proportion of variance in height explained by the score was 23% (*r* = 0.48, 95% CI: [−0.10,0.81]).

## Supplementary Tables

**Table S1. Quality control metrics for the Erfurt samples.** See Excel file.

**Table S2. The characteristics of the Erfurt samples.** See Excel file.

**Table S3. Details of the radiocarbon dating analyses.** See Excel file.

**Table S4. Pathogens detected in DNA sequences from the Erfurt samples.** See Excel file.

**Table S8. The results of the isotope analysis.** See Excel file.

**Table S11.** Properties of the Ashkenazi Jewish pathogenic variants. See Excel file.

**Table S5.** qpWave results. Each line presents the P-value for one *qpWave* test. The reference European populations were modern Russian, Norwegian, French, Spanish, Bulgarian, and Italian_North, with Primate_Chimp as an outgroup (first right population). The reference Middle Eastern populations were BedouinA, Lebanese, Jordanian, and Druze, with Primate_Chimp as an outgroup. Entries with P>0.05 are highlighted in dark green, and entries with 0.01≤P≤0.05 in light green. The only case with P>0.05 with respect to European populations is when the left populations are Erfurt Ashkenazi Jews (EAJ) and modern Ashkenazi Jews (MAJ). When Erfurt is replaced by Erfurt-ME or Erfurt-EU, the P-value is smaller, reflecting the differences in ancestry between each EAJ sub-group and MAJ. In the other tests, we replaced MAJ or EAJ with Sephardi (Turkish) Jews or with non-Jewish Italians and Germans. P-values were very small except when comparing Erfurt-ME and Sephardi Jews.

| Left population 1 | Left population 2 | P-value – European populations | P-value – Middle Eastern populations |
| --- | --- | --- | --- |
| Erfurt Ashkenazi Jews | Modern Ashkenazi Jews | 0.15 | 0.38 |
| Erfurt-EU | Modern Ashkenazi Jews | 3.2e-06 | 0.14 |
| Erfurt-ME | Modern Ashkenazi Jews | 0.031 | 0.63 |
| Erfurt Ashkenazi Jews | Turkish Jews | 1.8e-10 | 0.18 |
| Erfurt-EU | Turkish Jews | 4.0e-18 | 0.085 |
| Erfurt-ME | Turkish Jews | 0.0081 | 0.60 |
| Turkish Jews | Modern Ashkenazi Jews | 2.9e-14 | 0.39 |
| Erfurt Ashkenazi Jews | Germans | 2.1e-81 | 0.00029 |
| Germans | Modern Ashkenazi Jews | 8.4e-187 | 2.0e-8 |
| Erfurt Ashkenazi Jews | South-Italians | 0.00020 | 0.030 |
| South-Italians | Modern Ashkenazi Jews | 8.1e-7 | 0.012 |

**Table S6.** Determining the degree of endogamy in AJ in the past ≈700 years. We used simulations to quantify the maximal degree of gene flow from Eastern Europeans into a group of modern AJ such that this admixed group will remain consistent with being a clade with unadmixed AJ. Our unadmixed group was the modern AJ dataset used for the original *qpWave* analyses. For the admixed group, we used *n* = 30 modern AJ genomes that were not used in the original analysis (sample size selected to match the size of the Erfurt sample; Methods 5.3). In the admixed group, we replaced a given proportion of the genome (columns) with haplotypes from Eastern European sources (Methods 5.3). The haplotype lengths were determined based on the assumed admixture times (rows; Methods 5.3). Each entry in the table shows the P-value for a *qpWave* test comparing the admixed and unadmixed groups with respect to European populations (as in Table S5; Methods 5.3). Cells with P>0.05 are highlighted in green. The results suggest an upper bound of about 2-4% on the degree of Eastern European gene flow separating modern and Erfurt AJ.

|  |  | Proportion of Eastern European ancestry |  |  |  |  |
| --- | --- | --- | --- | --- | --- | --- |
|  |  | 1% | 2% | 4% | 6% | 8% |
| Admixture time (generations before present) | 20 | 0.36 | 0.34 | 0.23 | 0.034 | 0.0014 |
|  | 15 | 0.33 | 0.091 | 0.0010 | 1.7e-07 | 2.2e-10 |
|  | 10 | 0.040 | 0.040 | 0.0060 | 4.9e-05 | 2.6e-07 |
|  | 5 | 0.21 | 0.087 | 0.0018 | 3.5e-06 | 2.1e-09 |

**Table S7.** qpAdm P-values for models with EAJ as the target group. In all models, EAJ was the target group and there were three source populations: Middle Eastern, Southern European, and Eastern/Western European. In the first six columns, the third source was Russians. In the last two columns, the source was Germans. The rows represent the Middle Eastern source in each model and the columns represent the Southern European source. Models with P>0.05 are highlighted in green, and models with P>0.01 in light green. The two columns under “transversions SNPs” present the results that were described in the main text, where we used only transversions to avoid bias due to ancient DNA damage. The next two columns (“all available SNPs”) present the results of the same models when all SNPs were used. The next two columns (“different outgroup”) present the results of the same models when we set Ami as the outgroup instead of Mbuti (using only transversions). The last two columns (“West-EU source”) use transversions and Mbuti as an outgroup. Overall, the models with South-Italian, Syrian/Lebanese, and Russian sources were plausible in all tests. Generally, a South-Italian source was more plausible than a North-Italian source, a Western European source was not plausible, and Lebanese, Syrian, Jordanian and Bedouin B were the more likely Middle Eastern sources.

| Middle Eastern source | Transversion SNPs (main analysis) |  | Robustness test: all available SNPs |  | Robustness test: different outgroup |  | West-EU source |  |
| --- | --- | --- | --- | --- | --- | --- | --- | --- |
|  | North Italian | South Italian | North Italian | South Italian | North Italian | South Italian | North Italian | South Italian |
| **Druze** | 0.0016 | 0.0031 | 6.9e-9 | 9.4e-10 | 0.0027 | 0.12 | 2.8e-5 | 0.0025 |
| **Egyptian** | 1.4e-14 | 5.4e-6 | 0 | 4.1e-17 | 2.3e-12 | 0.027 | 4.5e-24 | 9.5e-10 |
| **Bedouin A** | 1.7e-7 | 0.0042 | 2.9e-16 | 2.0e-6 | 2.8e-6 | 0.15 | 4.7e-17 | 7.7e-8 |
| **Bedouin B** | 0.022 | 0.21 | 0.00068 | 0.026 | 0.057 | 0.45 | 1.3e-6 | 0.00038 |
| **Palestinian** | 0.0021 | 0.058 | 3.7e-6 | 0.0020 | 0.0074 | 0.36 | 2.3e-11 | 1.5e-5 |
| **Lebanese** | 0.086 | 0.32 | 0.037 | 0.15 | 0.15 | 0.56 | 0.0058 | 0.020 |
| **Jordanian** | 0.0039 | 0.13 | 0.00014 | 0.039 | 0.014 | 0.41 | 2.8e-07 | 0.00049 |
| **Syrian** | 0.019 | 0.21 | 0.0064 | 0.061 | 0.053 | 0.47 | 0.00012 | 0.0027 |

**Table S9.** Ashkenazi Jewish mitochondrial DNA founder haplogroups. Previous studies showed that modern AJ carry four founder haplogroups, accounting for 40% of all Ashkenazi haplogroups [117]. We compared the frequencies of these haplogroups in modern AJ [118] and Erfurt. We excluded the two children of family A since their mother is also in the sample. The frequency of K1a1b1a in Erfurt (35%) was significantly higher than in modern AJ (P=0.041; two-tailed binomial test (binom.test in R)). The frequency of K1a1b1a in Erfurt-ME (7/13=54%) was higher than the frequency in Erfurt-EU (2/10=20%), but given the small sample sizes, the difference was not statistically significant (P=0.20; two-tailed Fisher’s exact test). The combined count of N1b2 (also called N1b1b1), K1a9, and K2a2a1 carriers (3/31; 9.7%) was lower in EAJ than expected based on modern AJ frequencies (20%), but again the result was not statistically significant (P=0.18; two-tailed binomial test).

| Haplogroup | Freq. in modern AJ | Freq. in Erfurt | Freq. in Erfurt-ME | Freq. in Erfurt-EU | Freq. in low-cov samples |
| --- | --- | --- | --- | --- | --- |
| K1a1b1a | 20% | 11/31 | 7/13 | 2/10 | 2/8 |
| N1b2 | 9% | 1/31 | 1/13 | 0/13 | 0/8 |
| K1a9 | 6% | 2/31 | 1/13 | 0/10 | 1/8 |
| K2a2a1 | 5% | 0 |  |  |  |

**Table S10.** Demographic models for the AJ founder event. The table presents the parameters we inferred for various demographic models of AJ history. In all models, the population has been of constant size *N_a_* diploids until *T_b_* generations before present, and the population has grown exponentially starting *T_b_* − *d* generations ago and until reaching present size *N*_e_. In the single-population models (all models except (H); Figure 3A), the population size has been *N_b_* for *d* generations, starting *T_b_* generations ago. In the two-population model ((H); Figure 3E), the population has split *T_b_* generations ago into one population of size *N_b_* (representing EAJ) and another of size *N_a_* − *N_b_*. After *d* generations, these two populations merged with proportions *f* and (1 − *f*), respectively. In the consanguinity model (D), we assume that a proportion *α* of the EAJ samples were born to parents who were first cousins. In all models, we estimated the parameters numerically by maximizing the composite-likelihood of observing the given number of segments in each length bin (Methods 8.3). In some models, the maximum likelihood was obtained when some parameters were at the boundary of the search space (e.g., *N_a_* = 50*k* and *N*_e_ = 10^7^). For models (A) and (H), we computed 95% confidence intervals (shown under each point estimate) using parametric bootstrap by resampling the segment counts per bin (Methods 8.3). In models (A) and (B), we inferred the demographic parameters based on modern IBD sharing (Methods 8.3). In (A), we inferred the bottleneck duration (*d*) from the data, while in (B), we fixed it to 1. In the modern ROH model (C), we inferred the model parameters based on ROH segments observed in modern AJ (Methods 8.4). In the consanguinity model (D), we fixed all parameters that were inferred in model (A), and used ROH segments in the EAJ samples to infer *α*, the proportion of EAJ individuals whose parents were first cousins (Methods 8.5). In the narrower bottleneck model (E), we fixed *T_b_* from model (A), and inferred the population sizes *N_a_* and *N_b_* based on ROH segments in EAJ. We then fixed *N_a_* and *N_b_* to their inferred values, and inferred *d* and *N*_e_ using modern IBD (Methods 8.6). In the longer bottleneck model (F), we fixed *N_a_* and *N_b_* from model (A), and inferred *T_b_* based on ROH in EAJ. As in (E), we then inferred *d* and *N*_e_ using the modern IBD data (Methods 8.6). In the joint likelihood model (G), we inferred the demographic parameters based on both ancient and modern data (Methods 8.7). Finally, in the two-population model (H), we inferred all parameters based on both ancient and modern data, including *f*, the proportion of modern AJ lineages that descend from EAJ (Methods 8.8).

|  | A | B | C | D | E | F | G | H |
| --- | --- | --- | --- | --- | --- | --- | --- | --- |
|  | Modern IBD,<br>variable<br>bottleneck<br>length | Modern IBD,<br>single-<br>generation<br>bottleneck | Modern<br>ROH | Consanguinity | Narrower<br>bottleneck | Longer<br>bottleneck | Joint<br>likelihood | Two-population<br>model |
| $N_a$ | 47,961<br>[45,922-72,585] | 10,010 | 4,671 | 47,961 | 5,042 | 47,961 | 50,000 | 50,000<br>[50,000-97,465.5] |
| $N_b$ | 1,563<br>[1,363.5-1,750.5] | 480 | 1,295 | 1,563 | 529 | 1,563 | 905 | 627<br>[355-958] |
| $N_c$ | 363,484<br>[-3,831,093.0<br>-545,656.5] | 157,498 | $10^7$ | 363,483 | $10^7$ | 169,092 | $10^7$ | 1,652,527<br>[-6,694,946-2,456,011] |
| $T_b$ | 41<br>[39-43] | 31 | 30 | 41 | 41 | 74 | 51 | 46<br>[36-58] |
| $d$ | 20<br>[15-24] | 1 (fixed) | 11 | 20 | 15 | 48 | 25 | 22<br>[9-35.5] |
| | | | | $\alpha = 0.22$ | | | | $f = 0.52$<br>[0.41-0.99] |

**Table S12.** Evaluating the imputation accuracy of PHCP using Mendelian inconsistency. We tested the rates of Mendelian inconsistency in the imputed genomes of families A and B and compared to the inconsistency in unrelated individuals. The table presents the number of covered SNPs (before imputation) in all individuals, the number of homozygous SNPs in the imputed genome of the parent (this includes genotyped SNPs, as these were also imputed from haploid to diploid), and the percentage of SNPs that are homozygous to the opposite allele in the child (out of the number of homozygous SNPs in the parent). In family A, Child 1 is the son (I14853) and Child 2 is the daughter (I14898). The unrelated individuals are the mother from family A (I14850) and the daughter from family B (I13869; “Child 1”), or another unrelated individual (I13866; “Child 2”).

|  | Number<br>of<br>covered<br>SNPs<br>Parent | Number of<br>imputed<br>homozygous<br>SNPs<br>Parent | Number<br>of<br>covered<br>SNPs<br>Child 1 | Proportion<br>of<br>inconsistent<br>SNPs (%)<br>Child 1 | Number<br>of<br>covered<br>SNPs<br>Child 2 | Proportion<br>of<br>inconsistent<br>SNPs (%)<br>Child 2 |
| --- | --- | --- | --- | --- | --- | --- |
| **Family A** | $562 \cdot 10^3$ | $15.52 \cdot 10^6$ | $519 \cdot 10^3$ | 0.23% | $113 \cdot 10^3$ | 0.59% |
| **Family B** | $701 \cdot 10^3$ | $15.52 \cdot 10^6$ | $601 \cdot 10^3$ | 0.19% | | |
| **Mother**<br>**of family**<br>**A vs**<br>**unrelated** | $562 \cdot 10^3$ | $15.52 \cdot 10^6$ | $601 \cdot 10^3$ | 2.13% | $643 \cdot 10^3$ | 2.15% |

**Table S13.** Carriers of pathogenic founder variants. We detected eight carriers of at least one high-confidence pathogenic variant. The proportion of Erfurt-ME individuals who were carriers (7/13, 54%) was higher than in Erfurt-EU (1/9, 11%, after removing the children of families A and B; P=0.07; two-tailed Fisher’s exact test). The proportion of K1a1b1a carriers who were pathogenic variant carriers (4/11; 36%) was higher than in carriers of other mtDNA haplogroups (4/19; 21%; P=0.42; two-tailed Fisher’s exact test).

| ID | Variants | Sex | Subgroup | mtDNA Haplogroup |
| --- | --- | --- | --- | --- |
| I13861 | GBA, NM_000157.3:c.84dupG<br>BRCA1, NM_007294.3:c.68_69delAG | F | Erfurt-ME | K1a1b1a |
| I13865 | F11, NP_000119.1:p.F301L | M | Erfurt-ME | L2a1l2a |
| I13870 | F11, NP_000119.1:p.E135X<br>G6PC, NP_000142.2:p.R83C | M | Erfurt-ME | K1a1b1a |
| I14736 | CFTR, NP_000483.3:p.G542X | F | Erfurt-ME | K1a1b1a |
| I14737 | ACADS, NM_000017.2:c.319C>T | F | Erfurt-ME | N1b1b1 |
| I14739 | LRRK2, NP_940980.3:p.G2019S<br>MEFV, NP_000234.1:p.V726A | F | Erfurt-ME | H5c2 |
| I14897 | CLRN1, NP_001182723.1:p.N48K | F | Erfurt-EU | J1c1 |
| I14903 | DHDDS, NM_001243564.1:c.124A>G | F | Erfurt-ME | K1a1b1a |

**Table S14.** Height estimates. The heights of 14 individuals were estimated based on eight long bone measurements and the anatomical method (Methods 1.3). For the polygenic score analysis, we used the mean over all estimates available for each individual.

| ID | Humerus<br>right | Humerus<br>left | Radius<br>right | Radius<br>left | Femur<br>right | Femur<br>left | Tibia<br>right | Tibia<br>left | Anatomically |
| --- | --- | --- | --- | --- | --- | --- | --- | --- | --- |
| I13868 | 163.2 | . | . | . | 164.6 | 166.1 | . | . | . |
| I14847 | 159.3 | . | . | . | . | . | . | . | . |
| I13864 | . | 167.3 | . | . | . | 164.6 | 162.5 | 162.3 | . |
| I13862 | 165.6 | 162.7 | 163.8 | . | 164.2 | 164.8 | 164.2 | . | 164.2 |
| I13866 | . | . | . | . | 159.1 | 160.1 | 157.1 | 160.1 | . |
| I14904 | . | 169.3 | . | . | 170.4 | 171.7 | 169.2 | 169.4 | . |
| I14850 | 146.1 | 146.1 | 150.1 | 148.4 | . | . | . | 148.2 | . |
| I13863 | . | . | . | . | 160.6 | 161.0 | 158.3 | 159.0 | . |
| I13861 | . | . | . | 150.8 | . | . | . | . | . |
| I14852 | . | . | . | . | . | 158.8 | . | 159.4 | . |
| I13867 | 149.4 | . | 150.4 | 149.8 | 149.7 | 148.7 | 148.4 | 147.7 |  |
| I14903 | 155.5 | . | 158.4 | 158.1 | 159.2 | 158.0 | 154.0 | 155 | 152.3 |
| I13869 | 157.1 | . | 158.1 | . | 157.5 | 155.3 | . | 157.1 | . |
| I14739 | 152.2 | . | . | 151.8 | 153.0 | 153.6 | . | . | . |

